# A Bacterial Bile Acid Metabolite Modulates T_reg_ Activity through the Nuclear Hormone Receptor NR4A1

**DOI:** 10.1101/2021.01.08.425963

**Authors:** Wei Li, Saiyu Hang, Yuan Fang, Sena Bae, Yancong Zhang, Gang Wang, Megan D. McCurry, Munhyung Bae, Eric A. Franzosa, Curtis Huttenhower, Lina Yao, A. Sloan Devlin, Jun R. Huh

## Abstract

Bile acids act as signaling molecules that regulate immune homeostasis, including the differentiation of CD4^+^ T cells into distinct T cell subsets. The bile acid metabolite isoallolithocholic acid (isoalloLCA) enhances the differentiation of anti-inflammatory regulatory T cells (Treg cells) by facilitating the formation of a permissive chromatin structure in the promoter region of the transcription factor forkhead box P3 (*Foxp3*). Here, we identify gut bacteria that synthesize isoalloLCA from 3-oxolithocholic acid and uncover a gene cluster responsible for the conversion in members of the abundant human gut bacterial phylum Bacteroidetes. We also show that the nuclear hormone receptor NR4A1 is required for the effect of isoalloLCA on Treg cells. Moreover, the levels of isoalloLCA and its biosynthetic genes are significantly reduced in patients with inflammatory bowel diseases, suggesting that isoalloLCA and its bacterial producers may play a critical role in maintaining immune homeostasis in humans.

## INTRODUCTION

The mammalian gut is a unique organ that harbors trillions of commensal bacteria (Sender et al., 2016) as well as a large number of immune cells that are segregated from the resident microbiota by a single layer of epithelial cells. The close proximity of microbial and host cells creates a challenge for the gut-residing immune cells, which must protect the host against harmful pathogens while maintaining tolerance to commensal microbes (Whibley et al., 2019). While it is known that immune cells shape the composition of gut microbiota (Pandiyan et al., 2019), a growing number of studies now indicate that gut microbes regulate the development and function of host immune cells. Furthermore, these studies incidate that alterations in the gut microbial community can lead to pathological conditions including disrupted barrier function and heightened intestinal inflammation (Lloyd-Price et al., 2019; Michail et al., 2012; Munoz et al., 2019).

Regulatory T (Treg) cells are a subset of CD4^+^ T helper cells that are abundant in the gut lamina propria. The transcription factor forkhead box P3 (FOXP3) promotes the differentiation of Treg cells (Fontenot et al., 2003; Hori et al., 2003), which play a critical role in promoting gut immune homeostasis (Sakaguchi et al., 2008). The abnormal regulation of these cells is closely associated with autoimmune and inflammatory diseases including colitis (Maul et al., 2005; Sakaguchi, 2005). Evidence indicates that Treg cells are regulated by the gut microbiota, suggesting that gut bacteria can directly influence adaptive immunity (Arpaia et al., 2013; Furusawa et al., 2013; Mazmanian et al., 2008). Indeed, both commensal strains (Atarashi et al., 2013; Sefik et al., 2015) and bacterially derived metabolites such as short-chain fatty acids (Arpaia et al., 2013; Smith et al., 2013) and polysaccharide A (Mazmanian et al., 2008) have been shown to modulate Treg differentiation.

Secondary bile acids are another group of bacterial metabolites that function as T cell modulators. Primary bile acids are steroidal compounds that are synthesized from cholesterol in the liver and secreted into the gut lumen after a meal, where they facilitate uptake of dietary fatty acids and vitamins (Ridlon et al., 2006). Once these compounds reach the lower gastrointestinal (GI) tract, they are chemically modified by the resident microbiota to form a class of metabolites called secondary bile acids (Ridlon et al., 2006). A growing body of evidence indicates that both primary and secondary bile acids affect host physiology, including metabolism and immune response (Fiorucci and Distrutti, 2015; Pols et al., 2017; Song et al., 2020; Thomas et al., 2008; Vavassori et al., 2009). Indeed, prior work has demonstrated that either groups of bile acids or the specific secondary bile acids modulate the function and differentiation of Treg cells (Campbell et al., 2020; Hang et al., 2019; Song et al., 2020). For example, we recently demonstrated that the bile acid metabolite isoallolithocholic acid (isoalloLCA), an isomer of the secondary bile acid lithocholic acid (LCA), enhances the differentiation of naïve T cells into Treg cells both in vitro and in vivo (Hang et al., 2019). LCA is one of the most abundant secondary bile acids, with mean concentrations of ∼160 μM in human cecal contents (Hamilton et al., 2007). While LCA is exclusively produced by gut bacteria (Ridlon et al., 2006), the biosynthetic pathway for the production of the isomeric compound isoalloLCA is unknown. This bile acid is absent from germ-free B6 mice (Hang et al., 2019), indicating that a microbiome is necessary for the production of this immunomodulatory metabolite and suggesting that gut bacteria may play a direct role in the synthesis of this compound.

Moreover, the way in which isoalloLCA enhances Treg cell differentiation is not fully understood. The Treg cell-modulating activity of isoalloLCA was found to depend on an enhancer of the *Foxp3* gene, the conserved noncoding sequence (CNS) 3 (Hang et al., 2019). This mechanism is unique amongst small molecules that promote Treg differentiation. The bacterial metabolites butyrate and isodeoxycholic acid (isoDCA) increase *Foxp3* gene expression and Treg cell differentiation in a CNS1-dependent manner (Arpaia et al., 2013; Campbell et al., 2020). While our previous work indicated that isoalloLCA increases H3K27 acetylation at the *Foxp3* promoter region, the host factor directly responsible for isoalloLCA-dependent upregulation of *Foxp3* gene expression is unknown.

In this work, we sought to determine whether human gut bacteria can biosynthesize isoalloLCA and to elucidate the mechanism by which isoalloLCA enhances Treg cell differentiation. Here, we identify species of Bacteroidetes, a phylum of bacteria that is abundant in the human gut, that produce isoalloLCA. Producer strains possess an inducible operon required for isoalloLCA synthesis *in vitro* and *in vivo* in germ-free (GF) mice monocolonized with Bacteroidetes species. We also demonstrate that isoalloLCA induces the differentiation of naïve T cells to Treg cells through the nuclear hormone receptor NR4A1. IsoalloLCA treatments result in the increased binding of NR4A1 at the *Foxp3* locus, leading to enhanced *Foxp3* gene transcription. Notably, both the levels of isoalloLCA and the abundance of isoalloLCA-producing bacterial genes are significantly reduced in human inflammatory bowel disease (IBD) cohorts compared to healthy controls. This study reveals a mechanism by which commensal bacteria regulate immune tolerance in the gut and suggests that isoalloLCA and the bacteria that produce this bile acid may temper inflammatory responses in the human GI tract.

## RESULTS

### Bacteroidetes Species Produce IsoalloLCA from the Gut Bacterial Metabolite 3-oxoLCA

To identify individual gut bacterial strains that produce isoalloLCA, we performed a screen of 990 gut bacterial isolates obtained from human stool samples (Paik, Yao et al., 2020, under review) (Figure S1A). Single colonies were incubated for up to 96 hours with either LCA or 3-oxolithocholic acid (3-oxoLCA) (100 μM), two bile acids found in the human gut (Hamilton et al., 2007; Hang et al., 2019) that we hypothesized could be precursors to isoalloLCA (Figure 1A).

**Figure 1.**
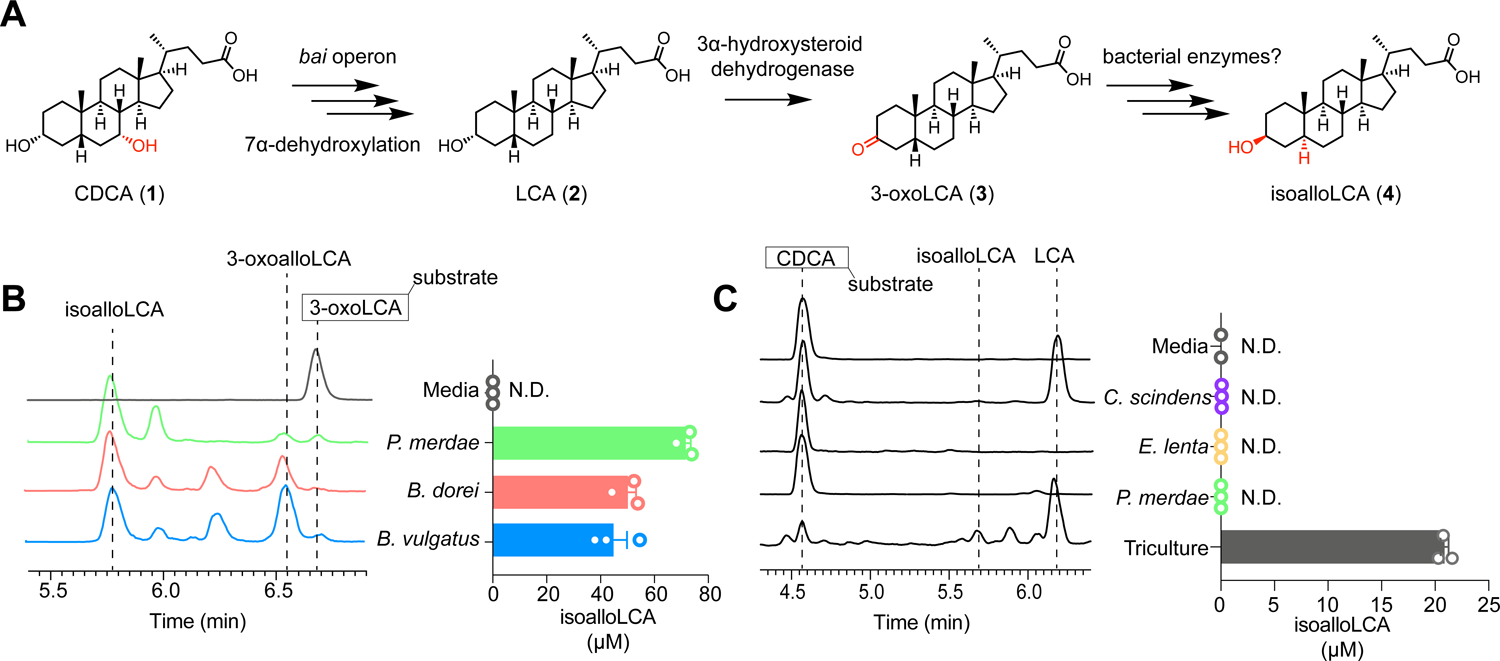
Identification of gut bacteria that produce isoalloLCA from the gut bacterial metabolite 3-oxoLCA. (A) Proposed biosynthetic pathway for the conversion of the primary bile acid chenodeoxycholic acid (CDCA) (1) to isoalloLCA (4) by human gut bacteria. Bacteria that produce isoalloLCA have not yet been identified. (B) Representative UPLC-MS traces (left) and quantification of isoalloLCA production (right) by bacteria from a screen of human isolates following incubation for 96 hours with 3-oxoLCA (100 μM) (n = 3 biological replicates per group, data are shown as the mean ± SEM, N.D. = not detected). (C) Representative UPLC-MS traces (left) and quantification of isoalloLCA production (right) showing that a triculture of *Clostridium scindens* ATCC 35703, *Eggerthella lenta* DSM 2243 and *Parabacteroides merdae* ATCC 43184 converted CDCA to isoalloLCA while monocultures of these bacteria did not produce isoalloLCA. *C. scindens* ATCC 35703 contains *bai* operon genes that can transform CDCA to LCA (Ridlon et al., 2006), while *E. lenta* DSM 2243 contains a 3α-HSDH, Elen_0690, that transforms LCA into 3-oxoLCA (Paik, Yao et al., 2020, under review) (n = 3 biological replicates per group, data are shown as the mean ± SEM, N.D. = not detected).

IsoalloLCA production was quantified using ultra-high performance liquid chromatography-mass spectrometry (UPLC-MS), and isoalloLCA-producing species were identified using 16S rRNA sequencing (Clarridge, 2004; Matsuki et al., 2004). While we did not identify any species that can directly convert LCA to isoalloLCA, we identified 16 bacterial species from 11 bacterial genera that converted 3-oxoLCA to isoalloLCA (*Bacillus*, *Bacteroides*, *Bifidobacterium*, *Catenibacterium*, *Collinsella*, *Eggerthella*, *Lachnospira*, *Lactobacillus*, *Parabacteroides*, *Peptoniphilus*, *Mediterraneibacter*) (Table S1). The top producers from the screen were *Parabacteroides merdae* P1-D8 (mean 72 μM), *Bacteroides dorei* P6-H5 (mean 50 μM), and *Bacteroides vulgatus* P6-D10 (mean 45 μM) (Figure 1B). These species all belong to the phylum Bacteroidetes, one of the two most abundant phyla in the human gut (Arumugam et al., 2011). The type strains of these bacterial species, *P. merdae* ATCC 43184, *B. dorei* DSM 17855 and *B. vulgatus* ATCC 8482, also converted 3-oxoLCA to isoalloLCA (Figure S1B), further verifying the results of the screen.

Human gut bacteria, specifically *Clostridium scindens* and its close genetic relatives, convert the primary bile acid chenodeoxycholic acid (CDCA) (1) to the secondary bile acid LCA (2) through enzymes encoded by the bile acid-inducible (*bai*) operon (Figure 1A) (Funabashi et al., 2020; White et al., 1980). Furthermore, in recent work we showed that a variety of gut bacteria from the phyla Actinobacteria and Firmicutes convert LCA (2) to 3-oxoLCA (3) through the action of 3α-hydroxysteroid dehydrogenase (3α-HSDH) enzymes (Paik, Yao et al., 2020, under review). Given our finding that gut bacteria convert 3-oxoLCA (3) to isoalloLCA (4), we hypothesized that multiple species of gut bacteria could be acting together to convert the host-produced bile acid CDCA to isoalloLCA. To test this hypothesis, we incubated CDCA (100 μM) with *C. scindens* alone, the 3α-HSDH-containing bacterium *Eggerthella lenta* alone, *P. merdae* ATCC 43184 alone, or a tri-culture of all three organisms. We observed isoalloLCA production from the triculture but not from any of the monocultures (Figure 1C). These data suggest that collaborative bacterial metabolism may contribute to the production of isoalloLCA.

### Identification of an IsoalloLCA-producing Gene Cluster in Bacteroidetes Species

We hypothesized that the biosynthetic pathway from 3-oxoLCA to isoalloLCA may involve the sequential actions of three enzymes: a 5β-reductase to oxidize the C3 ketone of 3-oxoLCA (3) to an enone (5), a 5α-reductase to reduce the enone (5) back to a ketone with inverted (α) stereochemistry at the C5 position to produce 3-oxoallolithocholic acid (3-oxoalloLCA) (6), and a 3β-HSDH to reduce the ketone (6) to a 3β alcohol (4) (Figure 2A, top red pathway). While the *bai* operon contains a 5β-reductase that acts on bile acids (baiCD), and 3β-HSDHs have been shown to produce iso-bile acids, 5α-reductases that act on bile acids have not yet been identified (Andersson and Russell, 1990; Devlin and Fischbach, 2015; Eminovic et al., 2001; Kang et al., 2008; Ridlon et al., 2010). Moreover, it is also possible that after enone (5) formation by the 5β-reductase, a 3β-HSDH could act first to form Δ4-isoLCA (7), and then a 5α-reductase could reduce the allylic alcohol to form isoalloLCA (4) (Figure 2A, bottom blue pathway). To validate our biosynthetic hypothesis, we incubated *P. merdae* ATCC 43184 and *B. dorei* DSM 17855 with 3-oxoLCA substrate and quantified the levels of putative pathway intermediates by UPLC-MS. In addition to isoalloLCA (4), 3-oxo-Δ4-LCA (5) and 3-oxoalloLCA (6) were produced, while Δ4-isoLCA (7) was not detected (Figures 2B and S2A). The lack of production of Δ4-isoLCA supports our initial hypothesis that 3-oxoLCA is converted to isoalloLCA by sequential 5β-reductase, 5α-reductase, and 3β-HSDH activities.

**Figure 2.**
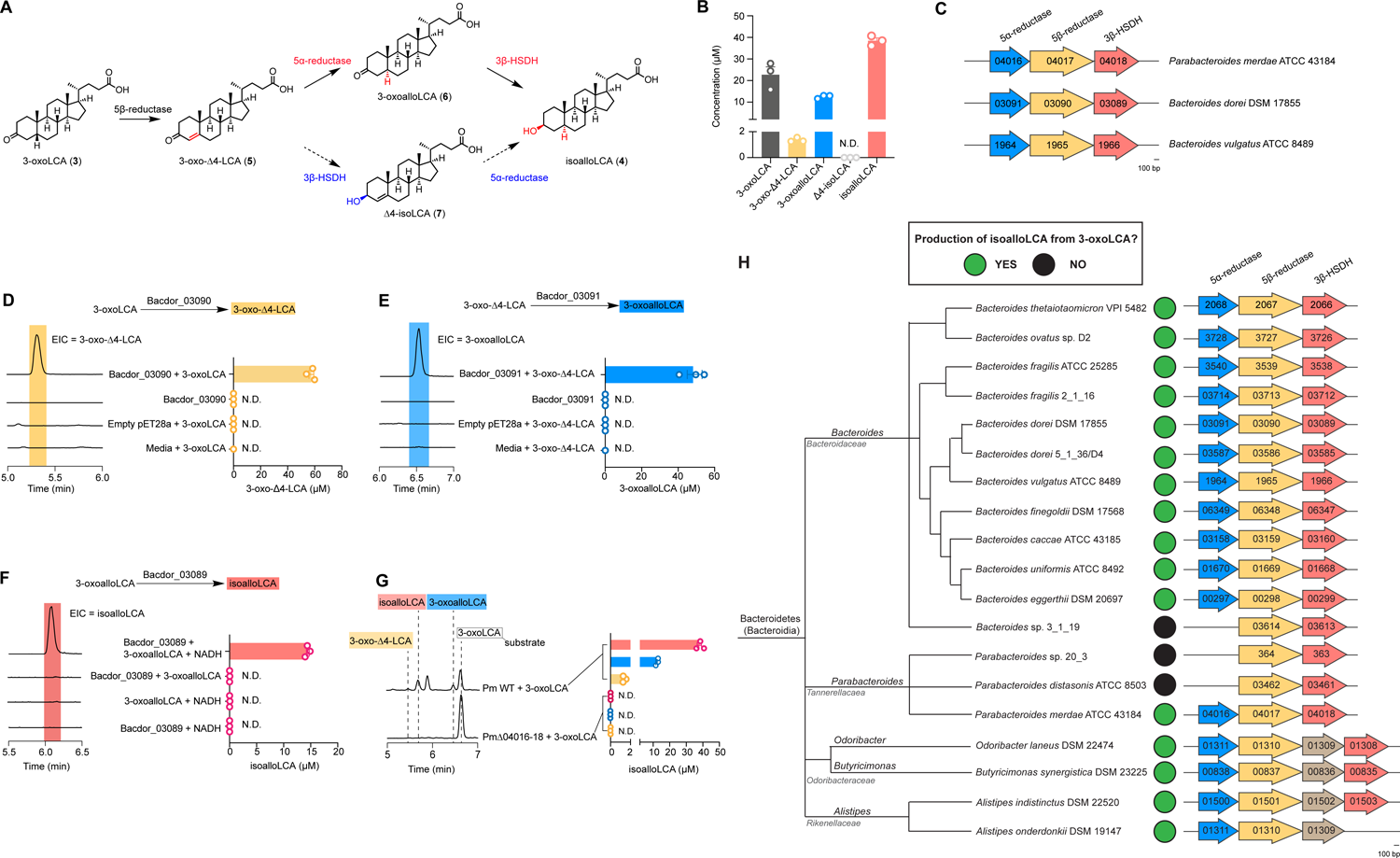
A gene cluster in Bacteroidetes converts 3-oxoLCA to isoalloLCA. (A) Proposed biosynthetic pathways for the conversion of 3-oxoLCA (3) to isoalloLCA (4). Subsequent UPLC-MS analysis of *Parabacteroides merdae* ATCC 43184 cultures incubated with 3-oxoLCA (Figure 2B) detected the proposed intermediate 3-oxo-Δ4-LCA (5) as well as 3-oxoalloLCA (6) from the top (red) pathway, but not Δ4-isoLCA (7) from the bottom (blue) pathway, supporting the hypothesis that subsequent actions of a 5β-reductase, a 5α-reductase, and a 3β-HSDH are responsible for the conversion of 3-oxoLCA to isoalloLCA by Bacteroidetes species. (B) The bile acid metabolites 3-oxo-Δ4-LCA, 3-oxoalloLCA, and isoalloLCA were detected by UPLC-MS in cultures of *P. merdae* ATCC 43184 incubated with 100 μM 3-oxoLCA for 96 hrs, while Δ4-isoLCA was not detected (n = 3 biological replicates per group, data are shown as the mean ± SEM, N.D. = not detected). (C) BLASTP searches revealed a three-gene cluster in the isoalloLCA producers *Bacteroides dorei* DSM 17855, *P. merdae* ATCC 43184, and *B. vulgatus* ATCC 8489. Bacdor_03091, Parmer_04016 and Bvu_1964 encode putative a 5⍺-reductase, Bacdor_03090, Parmer_04017 and Bvu_1965 encode a putative 5β-reductase, and Bacdor_03089, Parmer_04018 and Bvu_1966 encode a putative 3β-HSDH (see Table S2). (D-E) Representative UPLC-MS traces (left) and quantification of bile acid product (right) showing conversion of 100 μM 3-oxoLCA to 3-oxo-Δ4-LCA and 100 μM 3-oxo-Δ4-LCA to 3-oxoalloLCA by cell lysates of *E. coli* heterologously expressing Bacdor_03090 (**d**) and Bacdor_03091 (**e**), respectively. Extracted ion chromatograms (EICs) for 3-oxo-Δ4-LCA (m/z 371) (**d**) and 3-oxoalloLCA (m/z 373) (**e**) are shown (n = 3 biological replicates per group, data are shown as the mean ± SEM, N.D. = not detected). (F) Representative UPLC-MS traces (left) and quantification of isoalloLCA (right) showing conversion of 20 μM 3-oxoalloLCA to isoalloLCA by purified Bacdor_03089. EICs for isoalloLCA (m/z 375) are shown (n = 3 biological replicates per group, data are shown as the mean ± SEM, N.D. = not detected). (G) Representative UPLC-MS traces (left) and quantification of bile acids (right) showing *P. merdae* ATCC 43184 wild-type (Pm WT) produces isoalloLCA as well as the intermediates 3-oxo-Δ4-LCA and 3-oxoalloLCA from 3-oxoLCA (100 μM), while the gene cluster knockout strain of *P. merdae* (PmΔ04016-18) does not produce detectable amounts of these metabolites (n = 3 biological replicates per group, data are shown as the mean ± SEM, N.D. = not detected). (H) In vitro testing of Bacteroidetes strains containing complete or partial gene clusters homologous to the cluster identified in *Bacteroides dorei* DSM 17855, *P. merdae* ATCC 43184, and *B. vulgatus* ATCC 8489 revealed additional isoalloLCA producers. The presence of a 5α-reductase can be seen as a predictor of overall function. Phylogenetic tree of gut Bacteroidetes strains was adapted from (Karlsson et al., 2011).

We next performed BLASTP searches using a 5β-reductase, baiCD, from *Clostridium scindens* (Kang et al., 2008; Ridlon et al., 2010), a 5α-reductase, SRD5A1, from *Homo sapiens* (Eminovic et al., 2001), and a 3β-HSDH, Rumgna_00694, from *Ruminococcus gnavus* (Devlin and Fischbach, 2015) as query sequences against the genomes of the type strains of isoalloLCA-producing bacteria (*P. merdae* ATCC 43184, *B. dorei* DSM 17855 and *B. vulgatus* ATCC 8482). We identified a gene cluster that encodes homologs of all three putative enzymes in the pathway (Figure 2C, Table S2). Gene expression analysis by RT-qPCR revealed that the candidate 3β-HSDH homolog was significantly upregulated in *P. merdae* ATCC 43184 and *B. dorei* DSM 17855 in the presence of the substrate 3-oxoLCA compared to control-treated cultures, suggesting that this operon is inducible (Figure S2B).

To confirm the function of these candidate genes, we reconstituted this pathway *in vitro* by heterologously expressing individual genes from the *B. dorei* DSM 17855 cluster and incubating them with the relevant substrates. Bacdor_03090 converted 3-oxoLCA to 3-oxo-Δ4-LCA, while Bacdor_03091 converted 3-oxo-Δ4-LCA to 3-oxoalloLCA, indicating that these genes encode a 5β-reductase and 5α-reductase, respectively (Figures 2D, 2E and S2C). However, *E. coli* appeared to have native 3β-HSDH activity that could convert 3-oxoalloLCA to isoalloLCA (Figure S2D). We therefore purified the enzyme encoded by Bacdor_03089 and determined that this enzyme converted 3oxoalloLCA to isoalloLCA and is a 3β-HSDH (Figures 2F and S2E). To verify the function of the gene cluster in a native producer, we generated an in-frame deletion mutant of the entire candidate gene cluster in *P. merdae* ATCC 43184 (Figure S2F). The mutant strain did not produce either isoalloLCA or pathway intermediates previously shown to be produced by the wild-type strain (Figure 2B), demonstrating that this three-gene cluster is required for isoalloLCA production (Figure 2G).

Having identified a biosynthetic gene cluster for isoalloLCA production, we next sought to uncover more gut bacterial producers of isoalloLCA. Using BLASTP searches, we identified a group of species in the phylum Bacteroidetes that possess the cluster (Table S3). We then verified the ability of a subset of these species to produce isoalloLCA from 3-oxoLCA (Figure 2H). We also found that while strains lacking the 5α-reductase homolog in the cluster were not able to produce isoalloLCA, strains lacking the 3β-HSDH homolog, or possessing a putative 17β-HSDH inserted before the 3β-HSDH homologin the cluster, were still able to produce isoalloLCA (Figure 2H). These data suggest that the cluster does not need to be contiguous in order to produce isoalloLCA, and that there may be other 3β-HSDH genes elsewhere in the genomes of Bacteroidetes producers that have yet to be identified.

### NR4A1 Is Required for the iTreg-promoting Activity of IsoalloLCA

As previously reported, isoalloLCA, but not other LCA derivatives, strongly enhanced the differentiation of Treg cells (Hang et al., 2019). Furthermore, compared to isoalloLCA, the intermediate metabolites 3-oxo-Δ4-LCA and 3-oxoalloLCA have either no effect or greatly reduced effects on *in vitro*-induced Treg (iTreg) differentiation (Figures S3A-S3C).

To examine overall transcriptomic and genomic changes upon isoalloLCA treatment, we performed RNA-Seq and ATAC-Seq analysis of naïve CD4^+^ T cells cultured under non-polarizing TH0 conditions. Like TGFβ, isoalloLCA induced a set of Treg signature genes, including *Foxp3*, *Ctla4*, and *Il2ra* (Figure 3A), consistent with its Treg-enhancing activity. Compared to the DMSO-treated control, ATAC-Seq identified a few changes in the numbers of accessible chromatin peaks in isoalloLCA-treated cells (Figure 3B). Analysis of transcription factor binding motifs that were over-represented in the increased peaks showed enrichment of motifs for several families (Figure 3C, Table S4). Most suggestively in this context, the most significantly overrepresented motif was the NR motif that binds the NR4A family of nuclear receptors. This result is of interest because the NR4A family of nuclear receptors had been shown to influence Treg cell development (Fassett et al., 2012; Sekiya et al., 2011; Sekiya et al., 2013). We decided to test if isoalloLCA requires NR4A1 in order to enhance iTreg differentiation. Indeed, isoalloLCA treatment-associated ATAC-Seq peaks in the *Foxp3* promoter region were no longer enriched in NR4A1 deficient (KO) cells compared to isoalloLCA-treated control (CTL) wild-type (WT) cells (Figure 3D). Moreover, the effect of isoalloLCA on iTreg differentiation was significantly diminished in NR4A1 deficient cells under both TH0 and iTreg conditions (Figures 3E, 3F, S4A, and S4B). In contrast, NR4A1 deficiency did not compromise the iTreg-enhancing effects of TGFβ or retinoic acid (RA) (Figures 3E and 3F). Consistent with previous findings(Hang et al., 2019), the effects of isoalloLCA are mediated through the induction of mitochondrial ROS (mtROS), and induction of mtROS by mitoParaquat (mPQ) is sufficient to enhance iTreg differentiation (Figures S4A-S4D). While NR4A1 deficiency reduced the effect of mPQ (Figures S4A and S4B) on iTreg differentiation, it did not affect the production of mtROS by isoalloLCA (Figures S4C and S4D), suggesting that NR4A1 acts downstream of mtROS production (Figure S4E). Taken together, these data indicate that the iTreg-enhancing effects of isoalloLCA are dependent on the transcription factor NR4A1.

**Figure 3.**
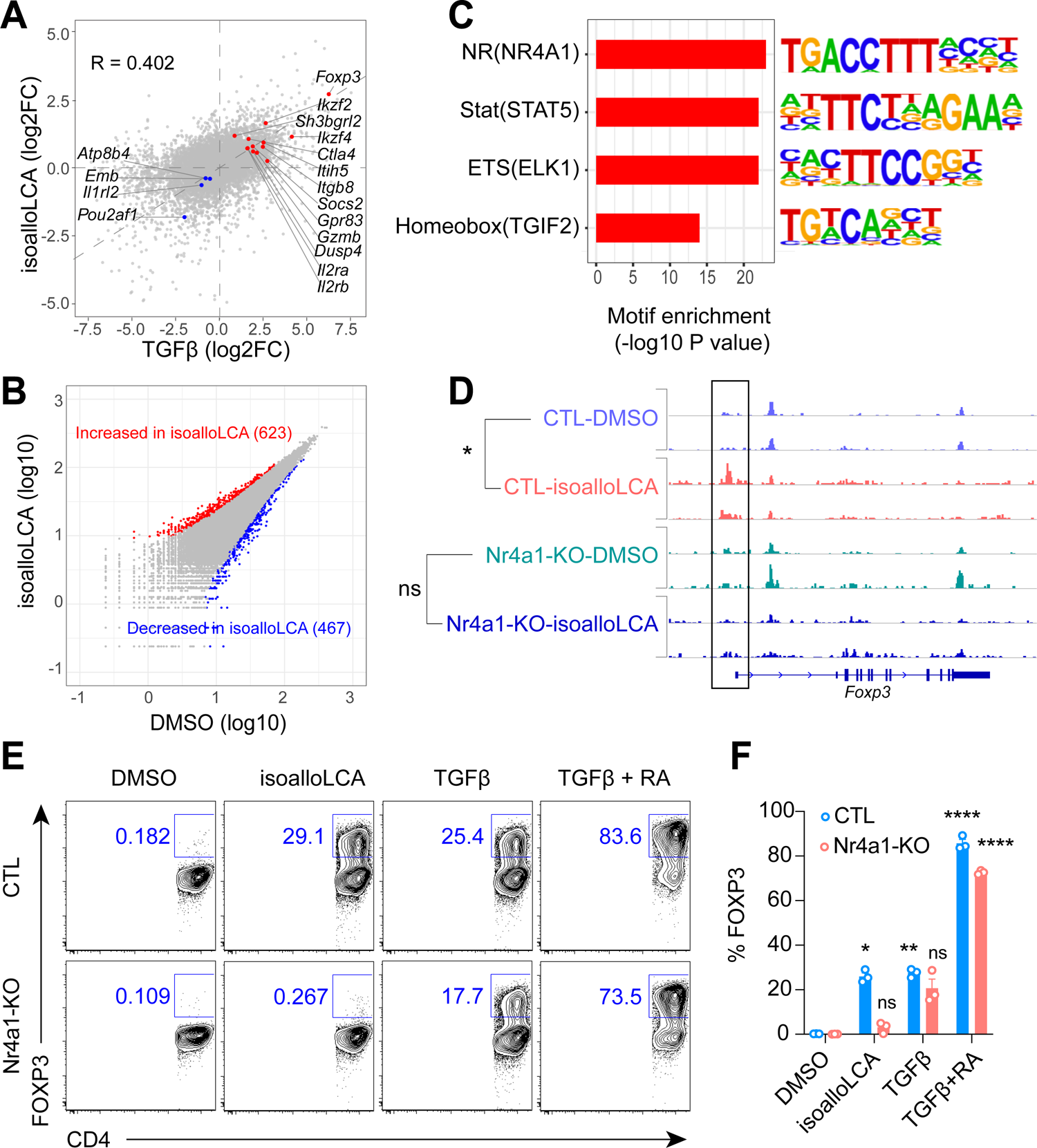
ATAC-Seq analysis identifies NR4A1 as required for Treg induction by isoalloLCA. (A) Two-way scatter plot comparing gene expression log2 scale fold changes under isoalloLCA or TGFβ treatment (all the expressed genes were used). Selected Treg signature genes are highlighted. R refers to the correlation coefficient. Naïve CD4^+^ T cells were purified from WT-B6 mice and culture under TH0 conditions (anti-CD3, anti-CD28, IL-2) in the presence of DMSO, isoalloLCA (20 µM), or TGFβ (0.25 ng/mL) for 48 hours. Cells were then collected for RNA-Seq analysis. Differentially expressed genes (DEGs) were determined by comparing DMSO- and isoalloLCA-treatments or DMSO-and TGFβ-treatments. (n = 3 biological replicates per group). (B) Scatter plot showing differentially enriched ATAC-Seq peaks between isoalloLCA and DMSO treatments. Naïve CD4^+^ T cells were purified from WT mice and culture under TH0 conditions (anti-CD3, anti-CD28, IL-2) in the presence of DMSO or isoalloLCA (20 µM) for 48 hours. Cells were then collected for the ATAC-Seq experiment (n = 2 biological replicates per group). (C) The top 4 transcription factor (TFs) binding motifs identified in differentially enriched ATAC-Seq peaks increased (Figure 3B, red), but not decreased (Figure 3B, blue), by isoalloLCA treatment. The specific TFs in the parenthese are the ones that have the smallest enrichment p values within the corresponding TF family. Data were analyzed using HOMER and the motifs were ranked by *P* value (see Method and Table S4). (D) Integrative Genomics Viewer (IGV) visualization of ATAC-Seq signals at the *Foxp3* gene locus. Duplicated samples for naïve CD4^+^ T cells were purified from WT-B6 (CTL) or NR4A1-deficient (KO) mice, cultured under TH0 condition (anti-CD3, anti-CD28, IL-2) in the presence of DMSO or isoalloLCA (20 µM) for 48 hours. Cells were then collected for the ATAC-Seq experiment. (n = 2 biological replicates per group, statistical tests were performed to compare the ATAC-Seq signals within the boxed region (unpaired t-test with 2-tailed *p*-value, ns, not significant, * *P* < 0.05). (E-F) Flow cytometry analysis and quantification of naïve CD4^+^ T cells purified from WT-B6 (CTL) or NR4A1-deficient (KO) mice, cultured under TH0 conditions (anti-CD3, anti-CD28, IL-2) in the presence of DMSO, isoalloLCA (20 µM), TGFβ (0.25 ng/mL), or TGFβ + retinoic acid (RA, 1 nM) for 72 hours are shown. Cells were stained with FOXP3 as a marker for Treg cells. (n = 3 biological replicates per group, data are shown as the mean ± SEM, two-way ANOVA with Tukey’s multiple comparisons test, each compared to the DMSO group, ns, not significant, * *P* < 0.05, ** *P* < 0.01, *** *P* < 0.001).

### NR4A1 Stimulates *Foxp3* Gene Expression and Enhances iTreg Differentiation

Increased chromatin accessibility and enrichment of the NR4A1 binding sites in the *Foxp3* locus suggest that NR4A1 may bind directly to the *Foxp3* promoter region upon isoalloLCA treatment. To test this hypothesis, we performed chromatin immunoprecipitation with an NR4A1 antibody. We found that, under non-polarizing TH0 conditions, isoalloLCA treatment led to an increased association of NR4A1 with the *Foxp3* locus 500 bp upstream of the transcription start site (Figure 4A). No increase in NR4A1 association was observed when cells were treated with DMSO control or when NR4A1-deficient cells were treated with isoalloLCA (Figure 4A). These results indicate that isoalloLCA induces recruitment of NR4A1 to the *Foxp3* gene promoter. To test if the association of NR4A1 directly activates the *Foxp3* promoter, we utilized a reporter construct driven by the *Foxp3* promoter (Zheng et al., 2010). NR4A1, but not its subfamily members NR4A2 or NR4A3, strongly activated the expression of a luciferase reporter gene placed under the control of the *Foxp3* promoter (Figures 4B and 4C). Deletion of putative NR4A1 binding sites in this region significantly reduced NR4A1-mediated activation of this reporter construct (Figures 4D and 4E). Moreover, naïve CD4^+^ T cells transduced with retrovirus carrying NR4A1, but not other NR4A proteins, exhibited enhanced Treg differentiation compared to those with vector control (Figures 4F and 4G). In contrast, NR4A1 over-expression in naïve CD4^+^ T cells with CNS3-deficiency resulted in reduced FOXP3 expression (Figures 4F and 4G). Together, these results demonstrate the direct involvement of NR4A1 in enhancing iTreg differentiation by stimulating *Foxp3* transcription in a CNS3-dependent manner (Figure S4E).

**Figure 4.**
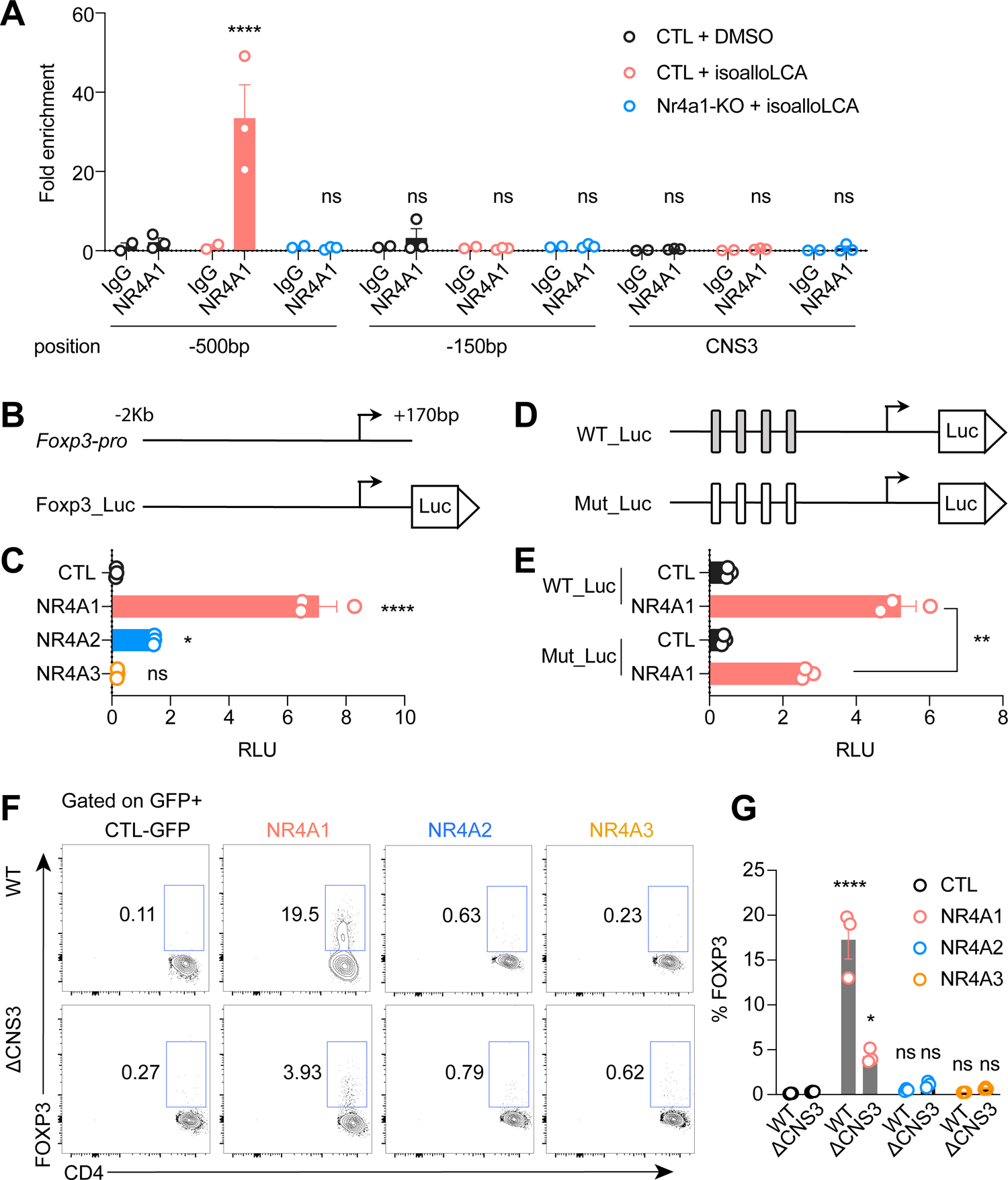
NR4A1 activates *Foxp3* gene transcription. (A) Chromatin immunoprecipitation analysis of NR4A1 on the *Foxp3* gene locus. Naïve CD4^+^ T cells purified from WT-B6 (CTL) or NR4A1-deficient (KO) mice and cultured under TH0 conditions (anti-CD3, anti-CD28, IL-2) in the presence of DMSO or isoalloLCA (20 µM) for 48 hours. The obtained chromatin was immunoprecipitated with mouse IgG or anti-NR4A1 antibody and analyzed by real-time PCR analyses. Primers targeting a region 500bp or 150bp upstream of the *FOXP3* transcription start site, or the CNS3 enhancer region, were used for qPCR quantification. Fold enrichment was presented relative to IgG (n = 2 for IgG group; n = 3 for NR4A1 group, biological replicates, data are shown as the mean ± SEM, one-way ANOVA with Dunnett’s multiple comparison test performed for NR4A1 group, compared to CTL-DMSO group, ns, not significant, **** *P* < 0.0001). (B) Schematics of a 2Kb (−2K to +170) *Foxp3* promoter region used in Foxp3-firefly luciferase reporter constructs (Foxp3_Luc). (C) Luciferase reporter assays with the construct shown in (B). Human embryonic kidney (HEK) 293 cells were transfected with a *Renilla* luciferase construct along with control (CTL), NR4A1-, NR4A2-, or NR4A3-expressing vectors. The ratio of firefly to *Renilla* luciferase activity, indicating Foxp3-Luc reporter transcriptional activity, is presented (n = 3 biological replicates per group, data are shown as the mean ± SEM, one-way ANOVA with Dunnett’s multiple comparison test, compared to CTL group, ns, not significant, * *P* < 0.05, **** *P* < 0.0001). (D) Schematics of 1Kb (−800 to +170) *Foxp3* promoter-firefly luciferase (WT_Luc) and the mutant constructs with deletions encompassing 4 putative NR4A1 binding sites mutant (Mut_Luc). (E) Luciferase reporter assays with constructs shown in (D) (n = 3 biological replicates per group), data are shown as the mean ± SEM by unpaired t-test with 2-tailed *p*-value, ** *P* < 0.01). (F-G) Flow cytometry analysis and quantification of naïve CD4^+^ T cells purified from WTmice (WT) or CNS3-deficient mice (ΔCNS3) cultured under TH0 conditions (anti-CD3, anti-CD28, IL-2) and transduced with retrovirus carrying GFP only (CTL), NR4A1, NR4A2, or NR4A3. Cells were stained with FOXP3 as a marker for Treg cells (n = 3 biological replicates per group, data are shown as the mean ± SEM, two-way ANOVA with Tukey’s multiple comparison test, compared to CTL group, ns, not significant, * *P* < 0.05, **** *P* < 0.0001).

### Gut Bacteria Produce IsoalloLCA In Vivo

To determine whether gut bacteria can produce isoalloLCA in vivo, we first monocolonized germ-free (GF) B6 mice with *B. dorei* DSM 17855, *P. merdae* ATCC 43184 or *B. vulgatus* ATCC 8482. We then fed the mice chow containing 3-oxoLCA (0.3% w/w) (Figures 5A). In contrast to the robust in vitro production of isoalloLCA by these strains (Figure S1B), monocolonization of GF mice with these bacteria resulted in detectable but low levels of isoalloLCA in cecal contents (Figure 5B). Because *P. merdae* ATCC 43184 does not possess a bile salt hydrolase (BSH), it is possible that enterohepatic recirculation of 3-oxoLCA followed by conjugation to taurine in the murine liver could result in reduced levels of unconjugated substrate 3-oxoLCA available in the gut (Yao et al., 2018). To test the hypothesis that bacterial BSH activity could increase isoalloLCA production, we co-colonized GF B6 mice with either *P. merdae* wild-type (WT) or the cluster-deficient mutant (*P. merdae* Δ04016-18) and *Bacteroides* sp. 3_1_19, a Bacteroidetes species that lacks a 5α-reductase and does not produce isoalloLCA in vitro (Figure 2H) but has strong BSH activity (Yao et al., 2018), and fed the mice 3-oxoLCA in chow (0.3% w/w) (Figure 5A). Cecal contents of mice co-colonized with *P. merdae* WT and *Bacteroides* sp. 3_1_19 contained on average 24 picomol isoalloLCA/mg wet mass, a concentration that was within an order of magnitude of the concentration that we previously showed was sufficient to enhance Treg differentiation in vivo (mean 47 picomol/mg wet mass) (Hang et al., 2019) (Figure 5C). Importantly, the cecal contents of mice co-colonized with *P. merdae* Δ04016-18 and *Bacteroides* sp. 3_1_19 did not contain detectable amounts of isoalloLCA, demonstrating that the three-gene cluster is required for isoalloLCA production by *P. merdae* in vivo (Figure 5C).

**Figure 5.**
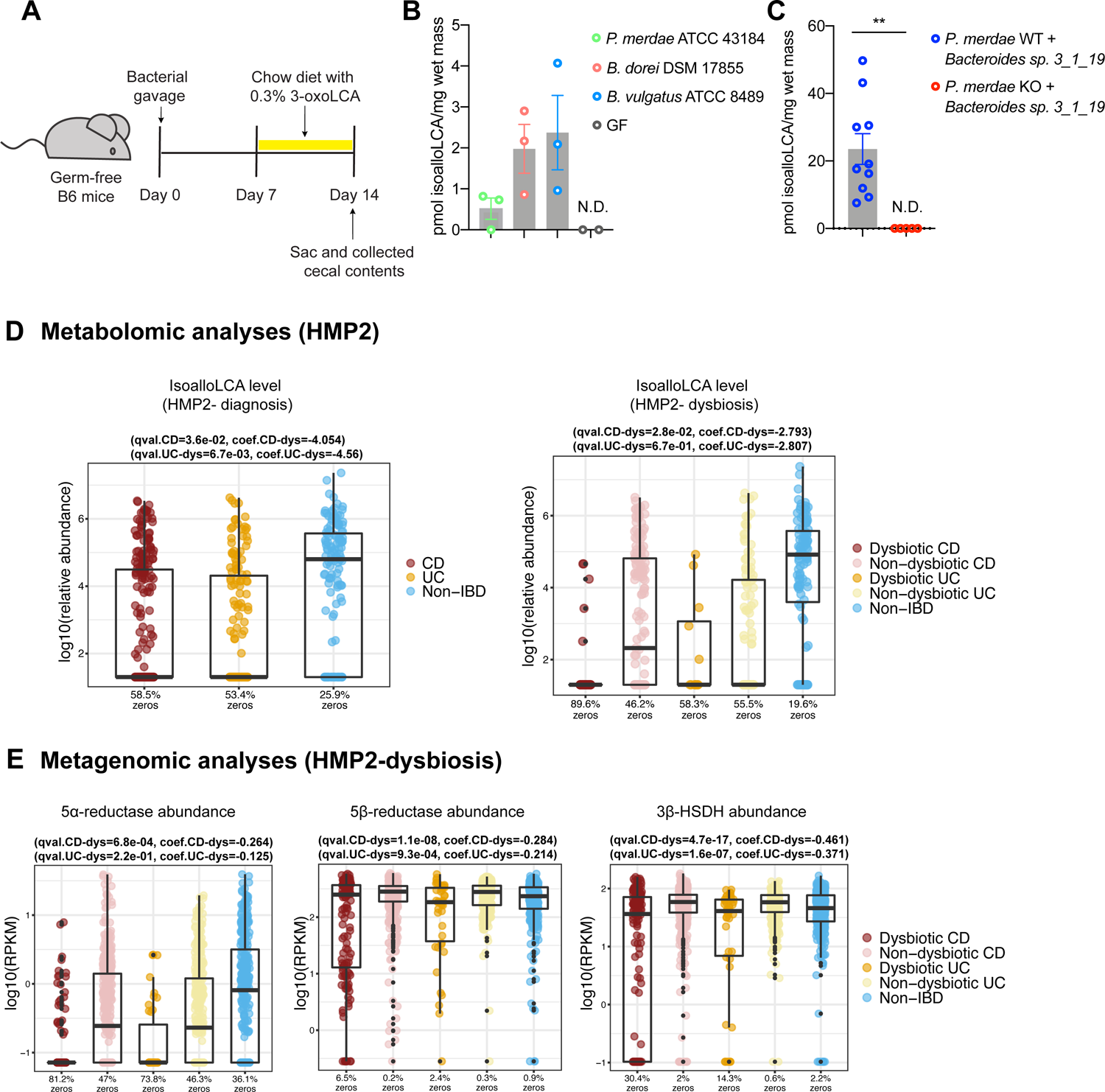
IsoalloLCA is produced by gut bacteria in vivo and is negatively correlated with IBD in humans. (A) Design of bacterial colonization experiment. Germ-free B6 mice were gavaged with bacteria on day zero. Starting on day seven, mono-colonized mice were fed a control diet or a diet containing 3-oxoLCA (0.3% w/w) for seven days. (B) Bile acid quantification of germ-free B6 mice fed with 0.3% 3-oxoLCA in chow (w/w) and mono-colonized with bacteria producing isoalloLCA, showing *in vivo* production of isoalloLCA in the cecum (*n* = 2, 3, 3, 3 respectively, biologically independent samples, data are shown as the mean ± SEM, N.D. = not detected). (C) Quantification of isoalloLCA in the cecal contents of germ-free B6 mice colonized with either *P. merdae* WT (Pm WT) or *P. merdae*Δ04016-18 (Pm KO) and *Bacteroides* sp. 3_1_19 (n = 10, 5 respectively, biological replicates pooled from two independent experiments, data are shown as the mean ± SEM by unpaired t-test with 2-tailed p-value, ** *P* < 0.01, N.D = not detected). (D) Left: isoalloLCA abundance was significantly reduced (FDR *q*-values < 0.05) in longitudinal fecal samples from 67 Crohn’s disease (CD) subjects (*n*=265 samples) and 38 ulcerative colitis (UC) subjects (*n*=146 samples) compared to 27 non-IBD controls (*n*=135 samples). Right: isoalloLCA abundance was also significantly depleted in dysbiotic CD samples (*n*=48) relative to non-dysbiotic baselines (*n*=169). Zero values are plotted toward the y-axis minima and enumerated per-group as x-axis tick labels (in percent). Boxplot “boxes” indicate the first, second (median), and third quartiles of the data; points outside boxplot whiskers are outliers. Statistical significance was determined from linear mixed effects models (relevant *q*-values and coefficients are duplicated in the panel titles; see Table S5 for full details). (E) The normalized abundances of the 5α-reductase, 5β-reductase, and 3β-HSDH homologs were significantly depleted (FDR *q*-values < 0.05) in the dysbiotic states of both CD and UC subjects from the HMP2 cohort compared to their individual baselines (based on 1,298 metagenomes from 50 CD subjects, 30 UC subjects, and 26 non-IBD controls). Boxplot definitions and statistical modeling mirror the descriptions from 5D (see Table S6 for full results).

### IsoalloLCA and its Biosynthetic Genes are Negatively Correlated with Inflammatory Bowel Diseases in Humans

We next investigated whether levels of the immunomodulatory metabolite isoalloLCA and the bacterial genes responsible for its production were altered in IBD patients. To do this, we focused on data from the HMP2 IBDMDB cohort (Lloyd-Price et al., 2019), which generated longitudinal metabolomic and metagenomic profiles of stool samples from subjects with Crohn’s disease (CD), ulcerative colitis (UC), and non-IBD controls. We found that isoalloLCA was significantly depleted in CD and UC subjects relative to the non-IBD controls (diagnosis) and further depleted within CD subjects under more severe (dysbiotic) perturbations of the gut microbiome (linear mixed effects models, FDR *q*-values < 0.05; Methods; Figure 5D). Moreover, the fold change in isoalloLCA in CD and UC subjects compared to controls was the largest among all identified bile acids in the metabolomics data from the HMP2 cohort (Table S5).

Using targeted functional profiling of HMP2 metagenomes, we found that the abundances of all three genes in the Bacteroidetes cluster were significantly depleted in the dysbiotic state of both CD and UC patients (FDR *q*-values < 0.05; Methods; Figure 5E and Table S6). The abundance of the 5α-reductase was also found to be significantly depleted in CD patients relative to controls in an independent IBD disease cohort, the Prospective Registry in IBD Study at MGH (PRISM) (Franzosa et al., 2019b) (linear model, FDR *q* = 0.0069; Methods; Figure S5A and Table S6). The 5α-reductase is a key gene in the biosynthesis of isoalloLCA because it is responsible for giving the molecule its unique 5α-, or allo-, configuration, whereas most bile acids have a 5β-configuration (Shiffka et al., 2017). Querying the NCBI nucleotide collection database (nt) revealed that most (303 out of 309) 5α-reductase homologs were found in Bacteroidetes strains, suggesting that this gene may play an important role in the human gut microbiome (Figure S5B and Table S3). Moreover, we found that in both metabolomics and metagenomics data sets, isoalloLCA levels and its biosynthetic genes were not only generally depleted but also entirely undetected in many samples in IBD (both CD and UC) compared to their controls (Figure 5D and 5E). Together, these data suggest that isoalloLCA levels and the bacterial genes responsible for its biosynthesis are correlated with the pathophysiology of IBD.

## DISCUSSION

Secondary bile acids are produced exclusively by gut bacteria and can be further transformed by the resident microbiota into a variety of metabolites with distinct physiological activities (Ridlon et al., 2006; Russell, 2003; Wahlstrom et al., 2016) (Paik, Yao et al., 2020, under review). In many cases, the bacteria and the biosynthetic pathways responsible for the production of these metabolites are unknown. Here, we used a high-throughput screen to identify isoalloLCA-producing bacteria from human stool isolates. Further, we identified a gene cluster consisting of a 5β-reductase, 5α-reductase, and 3β-HSDH that is responsible for isoalloLCA production from the gut metabolite 3-oxoLCA in the abundant gut bacterial phylum Bacteroidetes. This gene cluster is found in many prevalent Bacteroidetes strains, suggesting that this pathway may play an important role in the gut environment. Interestingly, isoalloLCA and other secondary bile acid metabolites, including deoxycholic acid (DCA), have been shown to inhibit the growth of non-Bacteroidetes species, affording a competitive advantage to the producing bacteria in a complex microbial community (Honda et al., 2020, under review) (Buffie et al., 2015). This study and our earlier work (Hang et al., 2019) suggest that isoalloLCA production by gut bacteria induces host immune tolerance through Treg induction, an immunoregulatory response that may benefit the host in the context of autoimmune and inflammatory diseases including IBD.

Recent studies have found that the immunomodulatory activities of bile acids are mediated by nuclear hormone receptors such as the farnesoid X receptor (FXR), the RAR-related orphan receptor gamma (RORγt), and the vitamin D receptor (VDR) (Campbell et al., 2020; Hang et al., 2019; Song et al., 2020). Interestingly, we found that the orphan nuclear receptor NR4A1 is responsible for the effect of isoalloLCA on Treg cells. NR4A family members have been shown to promote the development of thymic-derived Treg cells, and mice lacking all three members of this family in T cells show scurfy-like autoinflammatory phenotypes (Sekiya et al., 2013). However, the role of individual members of the NR4A family on peripherally induced Treg cells and the response of these nuclear receptors to environmental factors remains unknown. In contrast to a previous report showing that NR4A2 activates the *Foxp3* promoter (Sekiya et al., 2011), our results indicate that NR4A1, but not NR4A2 or NR4A3, promotes differentiation. Our findings suggest a model in which isoalloLCA induces an open chromatin region at the *Foxp3* gene promoter, a change that exposes the NR4A1 binding sites in this region, thereby leading to the activation of *Foxp3* transcription. In addition, NR4A1 is a downstream gene of T cell receptor (TCR) signaling (Jennings et al., 2020; Moran et al., 2011), a finding that is consistent with our previous observation that increased TCR signaling leads to enhanced isoalloLCA-dependent activity on Treg cell differentiation (Hang et al., 2019). While earlier work suggested that the NR4A family proteins including NR4A1 may not require ligand binding for their activities (Wang et al., 2003), it will be interesting to examine in the future if isoalloLCA can directly interact with NR4A1 and enhance its transcriptional activity.

The causes of IBD are still unclear, but genetic, immunological and environmental factors are thought to contribute to the risk of disease onset and progression (Knights et al., 2014; Manichanh et al., 2012). Recent multi-omics studies revealed that altered intestinal bile acid pools are associated with IBD (Lloyd-Price et al., 2019). Bacterial metabolites of host-derived bile acids have been shown to modulate Treg cell activity and contribute to disease progression in animal models of IBD (Campbell et al., 2020; Song et al., 2020). Here, we identified bacteria and bacterial enzymes that generate a Treg-enhancing bile acid, suggesting that specific commensal bacteria likely mediate host immune tolerance through the chemical modification of host-derived compounds. Furthermore, we observed that IBD in human patients is associated with decreased bacterial production of the immunomodulatory compound isoalloLCA and reduced representation of responsible bacterial genes in the gut microbiome. These results suggest that microbiota-inspired therapies targeting the adaptive immune system could be developed as treatments for inflammatory diseases affecting the GI tract.

## ACKNOWLEDGMENTS

We thank members of the Devlin, Huh, and Clardy labs (Harvard Medical School-HMS) for helpful discussions. We thank the BPF Next-Gen Sequencing Core Facility and the ICCB-Longwood Screening Facility at Harvard Medical School for their expertise and instrument support, Julian Avila-Pacheco and Clary B. Clish for metabolomics support, Arijit A. Adhikari for technical support and advice. We thank Randy Longman for human stool samples, Laurie E. Comstock for the pLGB13 plasmid (Addgene plasmid #126618) and Leonor García-Bayona for her technical support. This work was supported by National Institutes of Health grants R01 DK110559 (J.R.H.), and MIRA R35 GM128618 (A.S.D.), a Harvard Medical School Dean’s Innovation Grant in the Basic and Social Sciences (A.S.D. and J.R.H.), a John and Virginia Kaneb Fellowship (A.S.D.), a Harvard Medical School Christopher Walsh Fellowship (L.Y.) and a Wellington Postdoctoral Fellowship (L.Y.). The computations in this paper were run in part on the FASRC Cannon cluster supported by the FAS Division of Science Research Computing Group at Harvard University.

## AUTHOR CONTRIBUTIONS

W.L., S.H., L.Y., A.S.D., and J.R.H. conceived the project and designed the experiments. W.L. performed the bacterial culture experiments, enzyme characterization, bile acid synthesis, and bile acid profiling by UPLC-MS. S.H. performed the mouse experiments and in vitro T cell and reporter assays. L.Y. performed the human isolate screen. S.B. and Y.Z. performed the bioinformatics analyses. M.D.M. performed BLASTP searches to identify the putative biosynthetic gene cluster. M.B. helped with chemical characterization of synthetic Δ4-isoLCA.

E.A.F. and C.H. supervised the computational analyses. Y.F. performed RNA-Seq and ATAC-Seq analyses. G.W. prepared ATAC-Seq library. W.L., S.H., Y.F., S.B., Y.Z., L.Y., A.S.D, and J.R.H. wrote the manuscript, with contributions from all authors.

## DECLARATION OF INTERESTS

A.S.D. is a consultant for Takeda Pharmaceuticals and HP Hood. J.R.H. is a consultant for CJ Research Center, LLC. C.H. is on the scientific advisory boards of Seres Therapeutics, Empress Therapeutics, and ZOE Nutrition.

## METHODS

### RESOURCE AVAILABILITY

#### Lead Contact

Further information and requests for resources and reagents should be directed to and will be fulfilled by the Lead Contact, Jun R. Huh.

#### Materials Availability

All unique/stable reagents generated in this study are available from the Lead Contact with a completed Materials Transfer Agreement.

#### Data and Code Availability

ATAC-Seq and RNA-Seq data are available at the Gene Expression Omnibus (GEO), with the accession code GSE163349 and GSE163350. Metabolomics profiles from the PRISM cohort were taken from the associated publication’s supporting information (Franzosa et al., 2019a). The metagenomic sequencing reads from 155 PRISM samples were downloaded from SRA BioProject PRJNA400072 in April 2019. The metagenomic sequencing reads for HMP2 metagenomes and metabolomes profiles were downloaded from http://ibdmdb.org in July 2020. All other data generated during this study are included in this article and its Supplemental Information files. The software packages used in this study are free and open source. Source code for Sickle is available from https://github.com/najoshi/sickle. Cutadapt is available from http://bioconda.github.io/recipes/cutadapt/README.html. Bowtie2 is available from http://bioconda.github.io/recipes/bowtie2/README.html. SAMtools is available from https://github.com/samtools/. deepTools is available from https://github.com/deeptools/deepTools. IGV is available from http://software.broadinstitute.org/software/igv/. MACS2 is available from https://pypi.org/project/MACS2/. R package DESeq2 is available from https://bioconductor.org/packages/release/bioc/html/DESeq2.html. HOMER is available from http://homer.ucsd.edu/homer/index.html. TopHat2 is available from https://ccb.jhu.edu/software/tophat/index.shtml. HTSeq is available from https://htseq.readthedocs.io/en/master/. kallisto is available from https://pachterlab.github.io/kallisto/download.html. R package clusterProfiler is available from https://bioconductor.org/packages/release/bioc/html/clusterProfiler.html. MaAsLin2 is available via http://huttenhower.sph.harvard.edu/maaslin as source code and installable packages.

ShortBRED is available via https://huttenhower.sph.harvard.edu/shortbred/. The R package limma is available from https://www.bioconductor.org/packages/release/bioc/html/limma.html. Analysis scripts employing these packages are available from the authors upon request.

## KEY RESOURCES TABLE

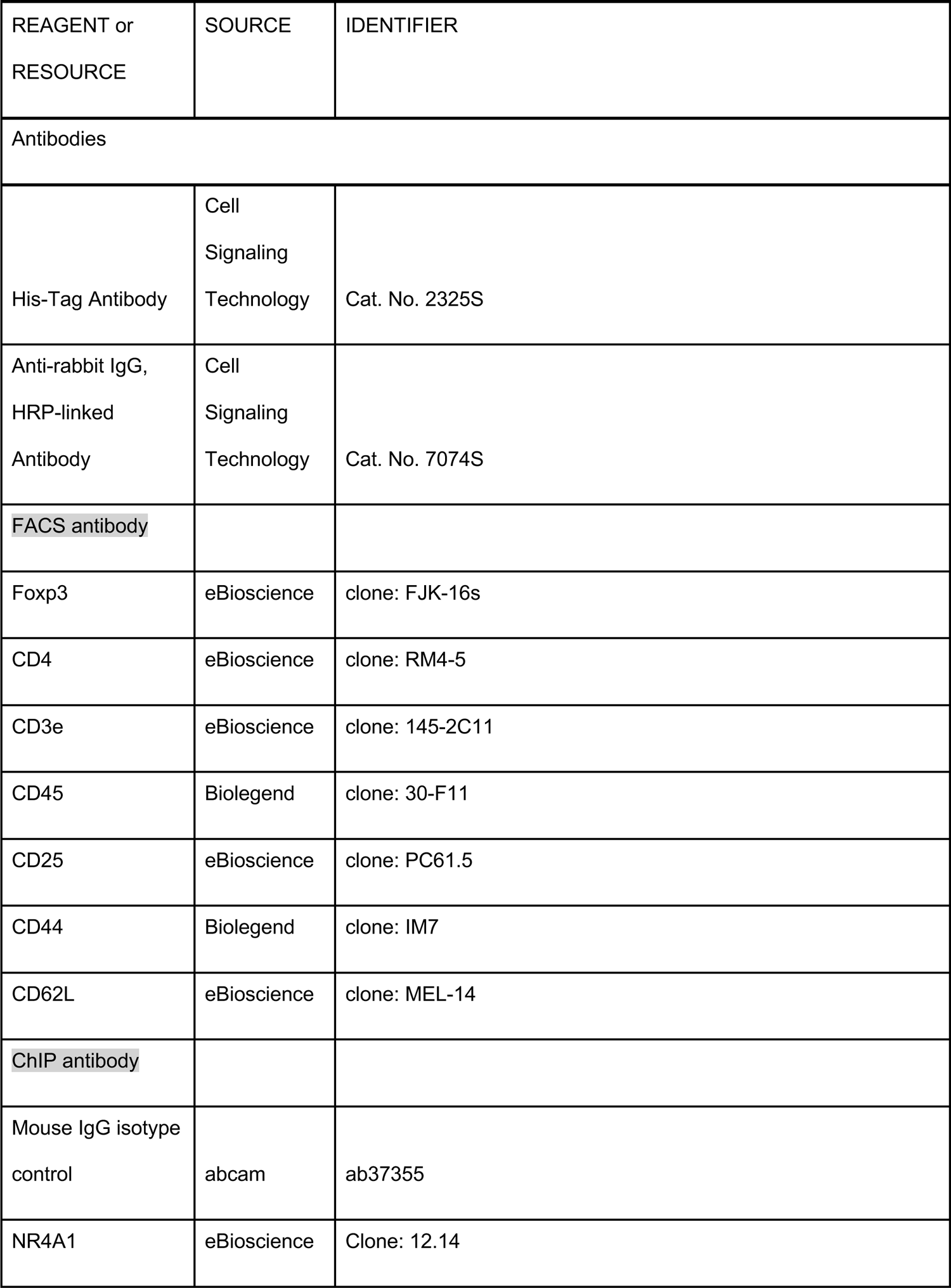
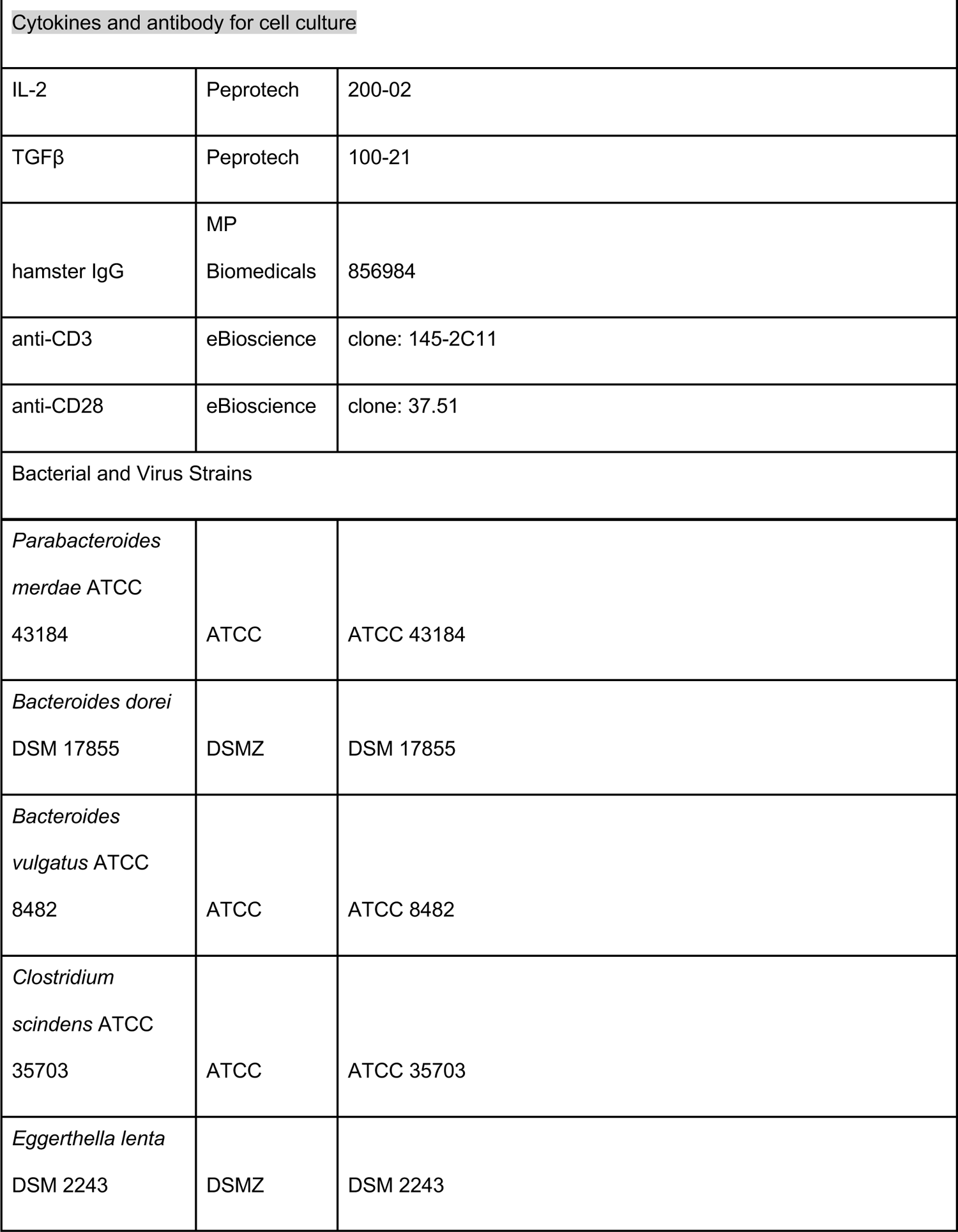
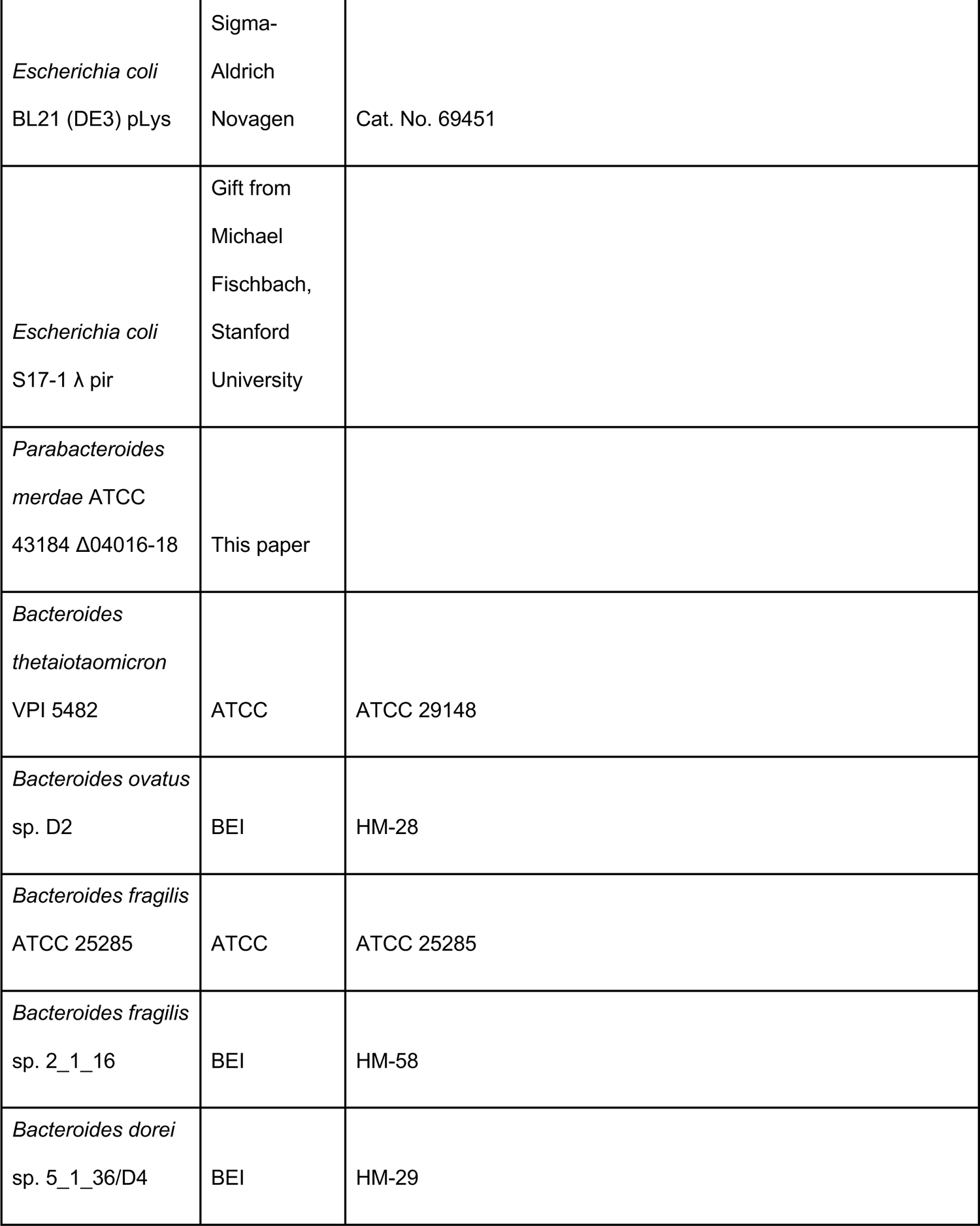
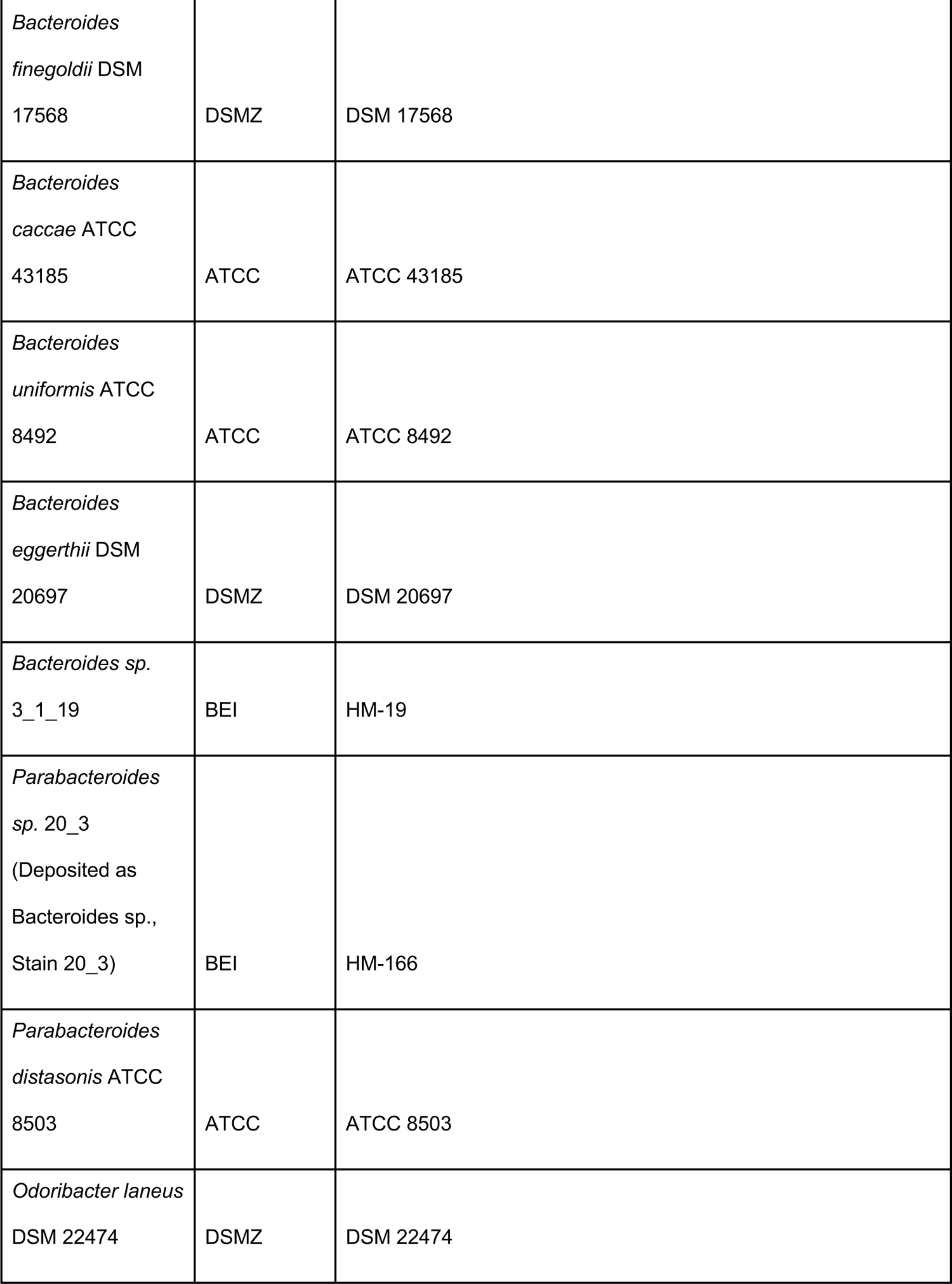
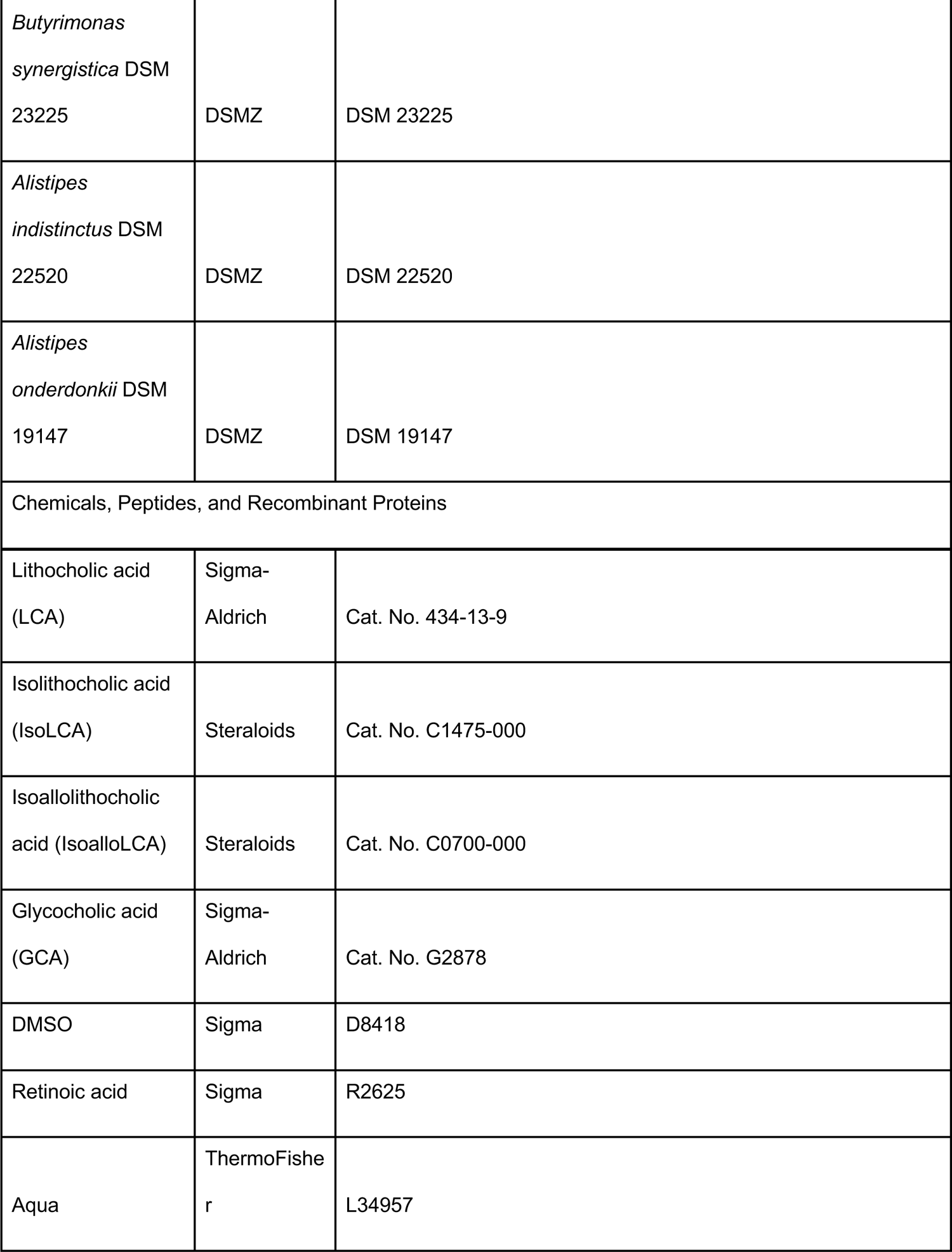
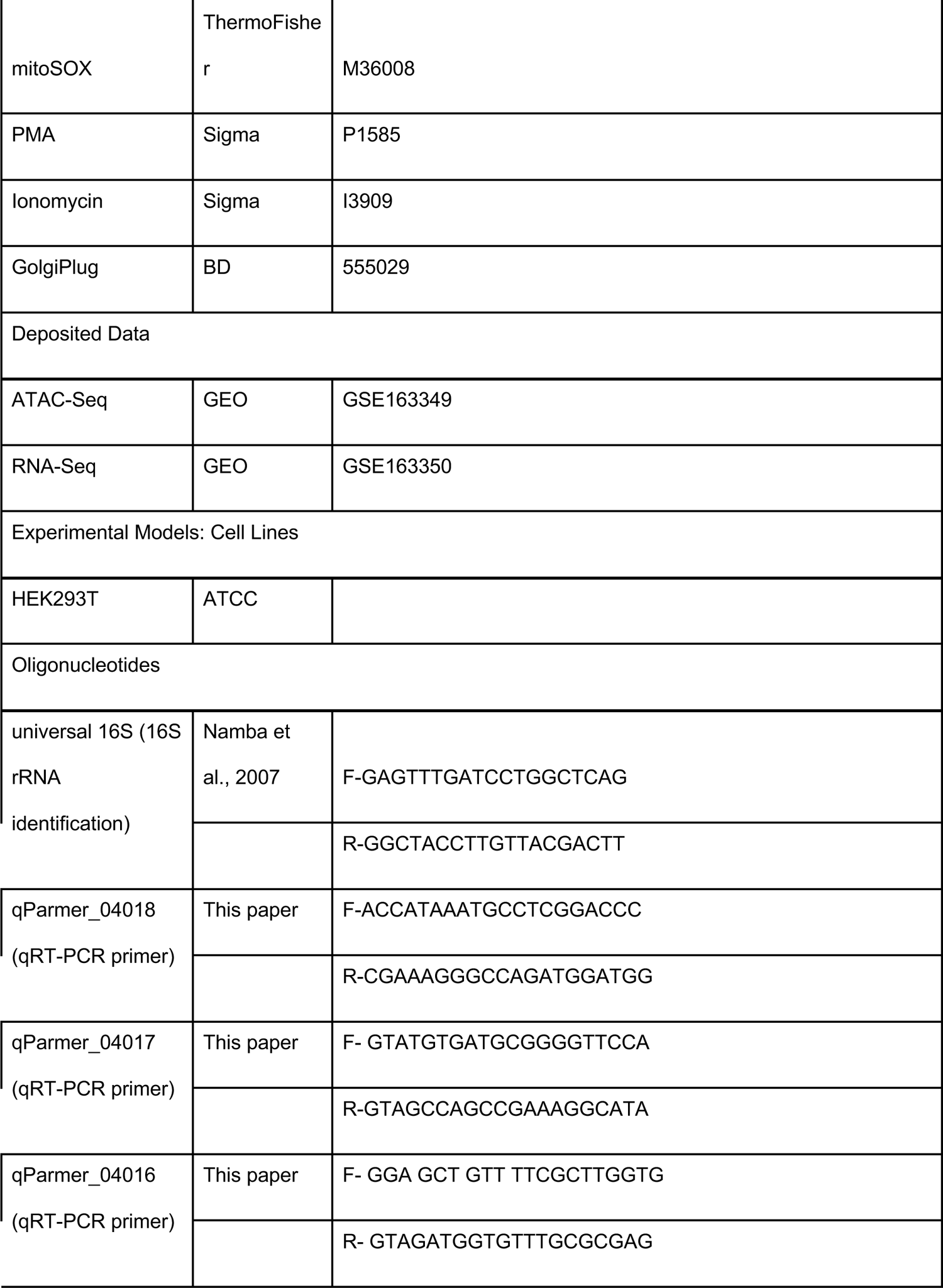
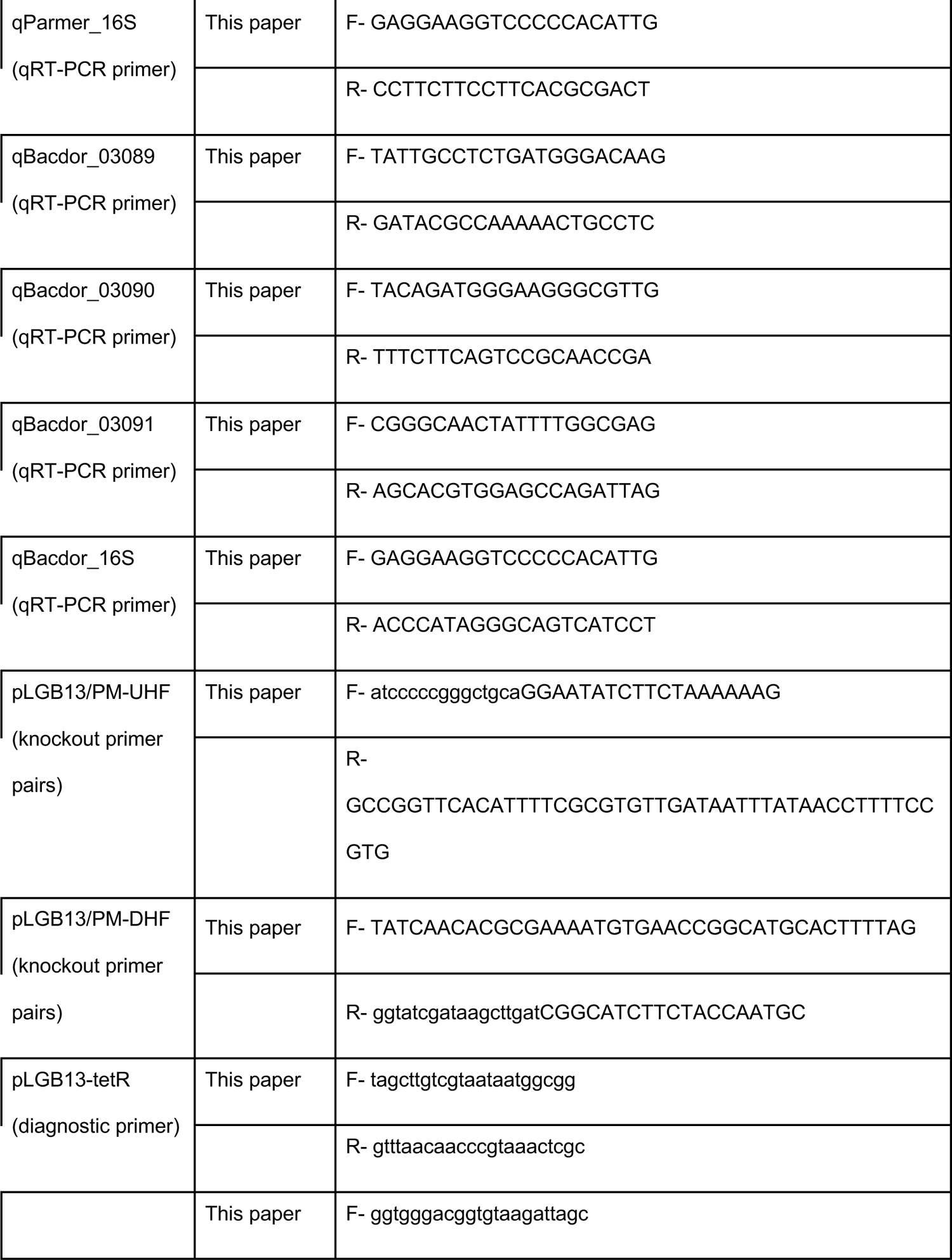
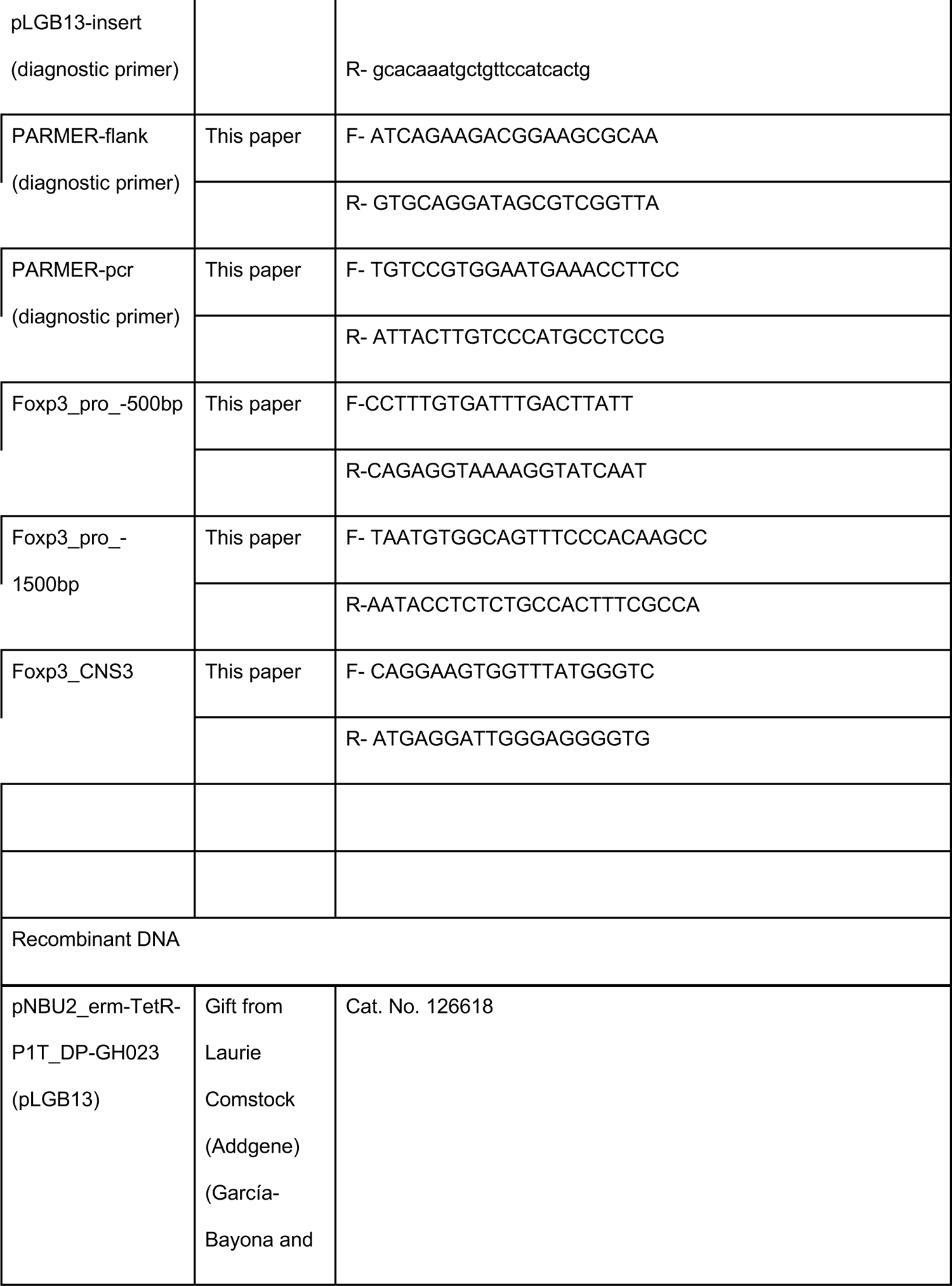
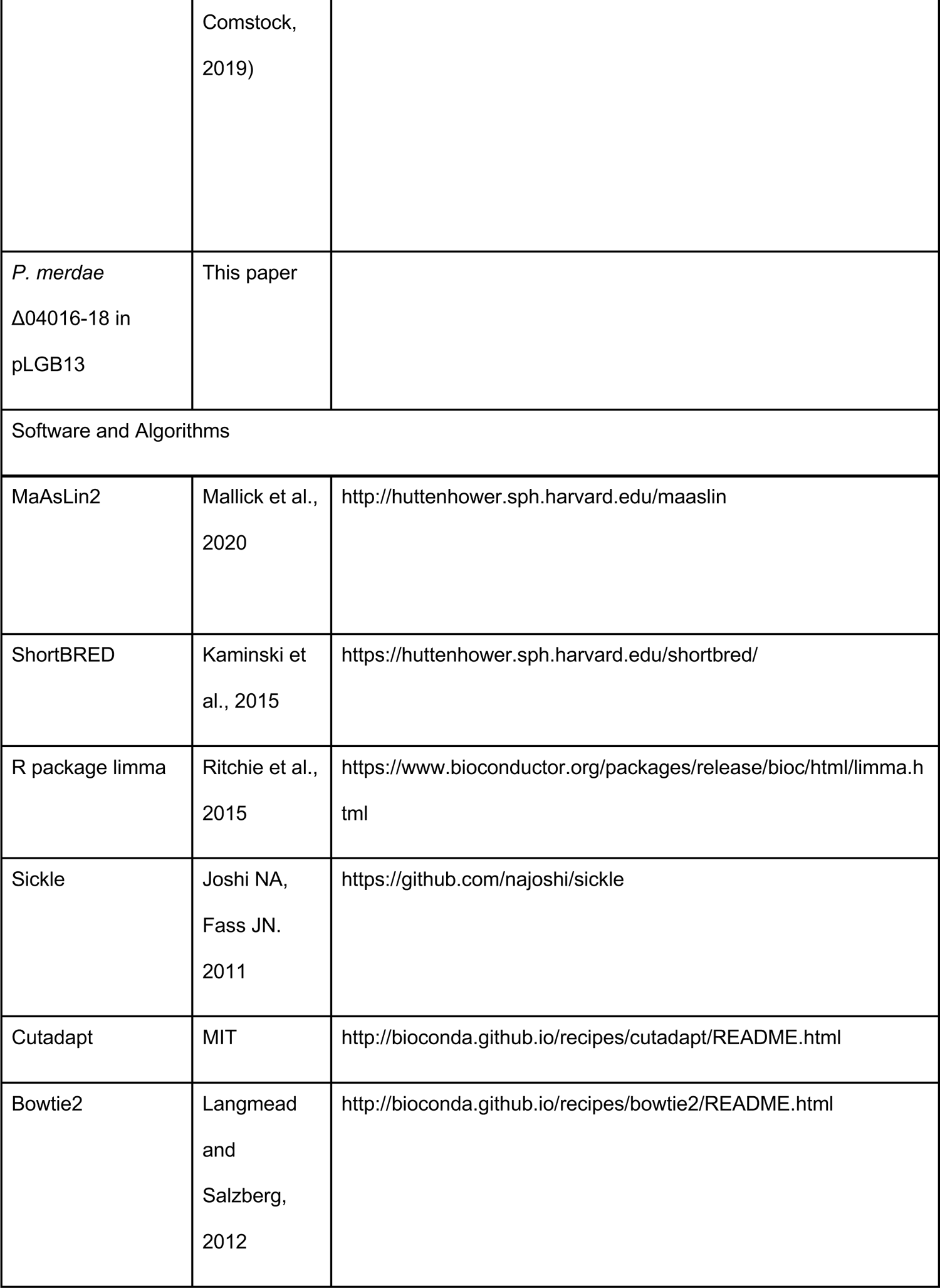
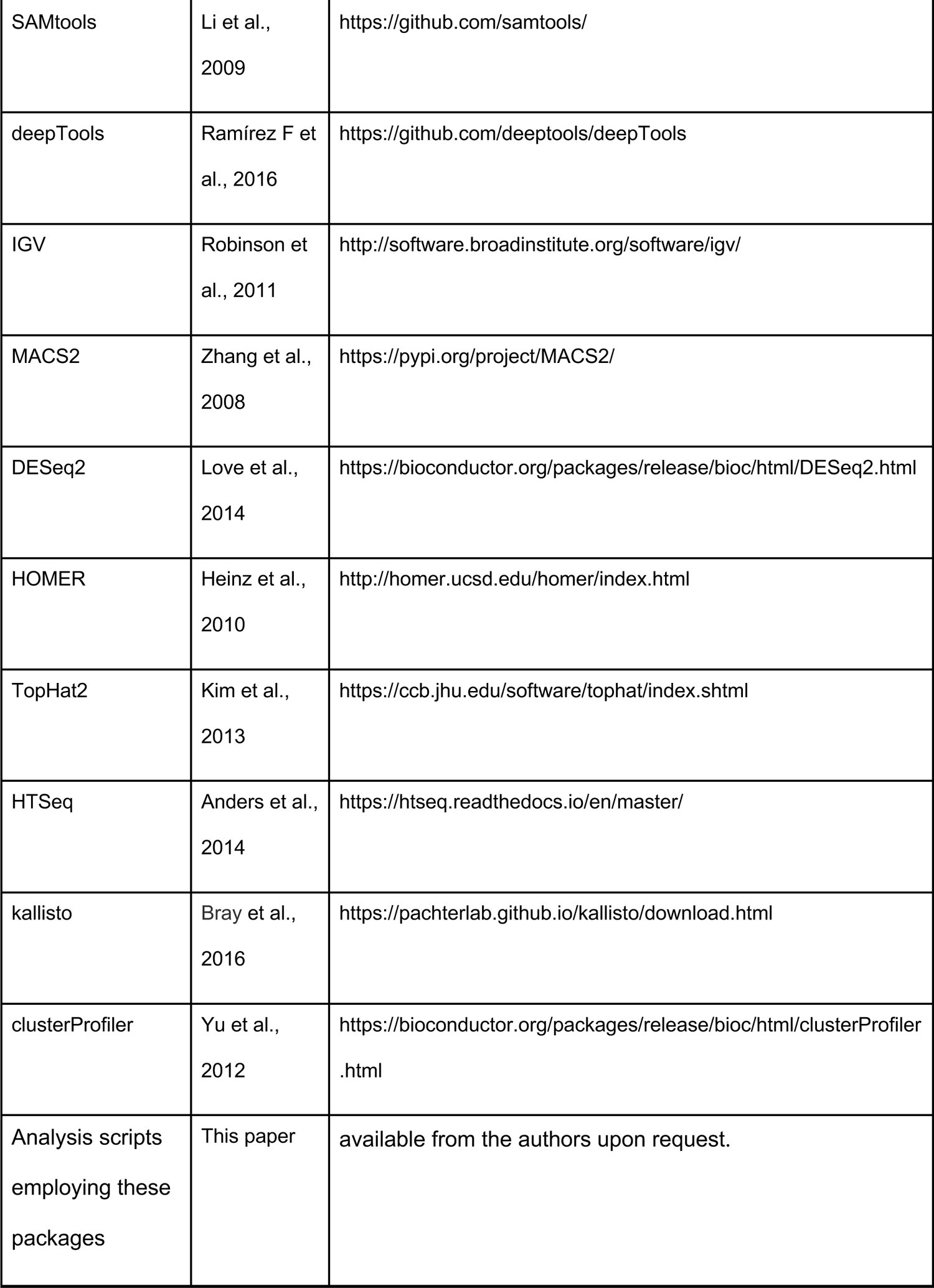

### EXPERIMENTAL MODEL AND SUBJECT DETAILS

#### Gnotobiotic Mice Experiments

Animals were maintained in specific pathogen-free conditions or GF conditions as appropriate. C57BL/6, Nr4a1-KO mice were purchased from Jackson Laboratory. All mouse studies were performed on mice ages 5-8 weeks old. For gnotobiotic experiments, sex-matched GF mice were orally gavaged with bacterial cultures and maintained in an Isocage system (Tecniplast). Mice were colonized with designated bacteria by oral gavage with bacterial culture in the biosafety station and kept in Isocages for the duration of the experiment. Control powder meal (Teklad Global 19% protein extruded diet, #2019) or a chow evenly mixed with 0.3% 3-oxoLCA (w/w) were autoclaved and provided to mice during the experiment. Bile acid substrates were provided as 0.3% in powder chow (w/w). Feces and cecal contents were collected at 6 days post-colonization for bile acid analyses. All animal procedures were approved by the Institutional Animal Care and Use Committee at Harvard Medical School.

### METHODS DETAILS

#### Chemical Synthesis

Detailed synthetic methods and characterization data for 3-oxo-Δ4-LCA, Δ4-isoLCA, and 3-oxoalloLCA are included in the Supplemental Information.

### Bacterial Culturing

All human gut bacteria were cultured in an anaerobic chamber (Coy Laboratory Products) with a gas mixture of 5% hydrogen and 20% carbon dioxide (balance nitrogen) unless otherwise stated. Bacteroidetes strains were cultured at 37 °C in brain heart infusion (BHI) (Bacto BHI, BD) media supplemented with 5 mg/L hemin (Sigma-Aldrich, H3039), 2.5 μL/L Vitamin K1 (Sigma-Aldrich, V3501), and 500 mg/L cysteine HCl (Sigma-Aldrich, C6852) (BHI+).

*Escherichia coli* strains were grown aerobically at 37 °C in Luria-Bertani (LB) or Terrific Broth (TB) media supplemented with ampicillin or kanamycin to select for the pLGB13 or pET28a(+) plasmid, respectively.

### Human Stool Isolate Screen

To screen human isolates for isoalloLCA production, isolates were retrieved from stock plates from our human fecal screen library (Paik, Yao et al., 2020, under review) and cultured in 600 μL Cullen-Haiser Gut (CHG) media (Hall et al., 2017), which consists of BHI supplemented with 1 % BBL vitamin K1-hemin solution (BD Biosciences, 212354), 1% trace minerals solution (ATCC, MD-TMS), 1 % trace vitamins solution (ATCC, MD-VS), 5% fetal heat-inactivated bovine serum (FBS) (Genesee Scientific, 25-514), 1 g/L cellobiose (Sigma-Aldrich, C7252), 1 g/L maltose (Sigma-Aldrich, M5895) and 1 g/L fructose (Sigma-Aldrich, F0127), containing 0.5% (w/v) arginine (Sigma-Aldrich, A5006) for 48 hours at 37°C in 96-well plates. Each isolate, as well as the negative controls, was then diluted 1:10 in new media containing 100 μM LCA (Sigma-Aldrich) or 100 μM 3-oxoLCA (Steraloids) for an additional 48 hours. 0.2 mL cultures were harvested and extracted for bile acid analyses (see below). This experiment was conducted once per substrate for all isolates from the original eleven library plates. Following bile acid analysis, we performed 16S rRNA sequencing on isoalloLCA-producing strains; subsequently, their function was verified in culture tubes in triplicate.

### *In Vitro* Mono-culture Assays for IsoalloLCA Production

Individual strains were plated from glycerol stocks onto BHI+ agar and grown for 3 days. Colonies were then inoculated into 3 mL of BHI+ media in Falcon™ Round-Bottom polystyrene tubes and grown overnight. The following day, these cultures were diluted to pre-log phase (OD600 = 0.1) in fresh BHI+ to a final volume of 3 mL. 3-oxoLCA was then added to each culture to obtain a final concentration of 100 μM substrate. Cultures were then incubated at 37 °C for 48 hours or 96 hours, and a 1 mL aliquot was removed for bile acid analyses (see below). The experiments were performed in triplicate and repeated twice unless otherwise stated.

### *In Vitro* Co-culture Assays for IsoalloLCA Production

Starting from single colonies, *Clostridium scindens* ATCC 35703, *Eggerthella lenta* DSM 2243 and *Parabacteroides merdae* ATCC 43184 were grown anaerobically for 24 hours in CHG + 0.5% arginine media at 37 °C. These starter cultures were diluted to pre-log phase (OD600 = 0.1) by addition of fresh CHG + 0.5% arginine media. Cultures were grown until mid-log phase (5 hours growth for *C. scindens* ATCC 35703, 4 hours growth for *E. lenta* DSM 2243, and 2 hours growth for *P. merdae* ATCC 43184) at 37 °C. Equal volumes of each culture were then combined and CDCA was added to the resultant cultures to obtain a final concentration of 100 μM of substrate. At the 96-hour time point, a 1 mL aliquot was removed for bile acid analyses (see below).

### Bioinformatics Searches for 5β-reductase, 5α-reductase and 3β-HSDH Candidate Genes

BLASTP searches were performed against the isoalloLCA producers *Bacteroides dorei* DSM 17855, *P. merdae* ATCC 43184, and *B. vulgatus* ATCC 8489 on Integrated Microbial Genomes, the US Department of Energy’s Joint Genome Institute (IMG JGI), using a 5β-reductase, baiCD, from *Clostridium scindens* (NCBI Protein accession code EDS05767) (Kang et al., 2008; Ridlon et al., 2010) (cutoff E-value of 10^-5^), a 5α-reductase, SRD5A1, from *Homo sapiens* (NCBI Protein accession code NP_001038) (Eminovic et al., 2001) (cutoff E-valoue 10^-5^), and a 3β-HSDH, Rumgna_00694, from *Ruminococcus gnavus* (NCBI Protein accession code EDN78833) (Devlin and Fischbach, 2015) (cutoff E-value of 10^-2^) as query sequences.

### Gene Expression Analysis

Total RNA was purified using AllPrep Bacterial DNA/RNA/Protein Kit (QIAGEN) from cell pellets of triplicate cultures of *P. merdae* ATCC 43184 or *B. dorei* DSM 17855 grown in ΒΗΙ+ with or without 3-oxoLCA for 1 hour. RNA was DNase treated, and cDNA was prepared using the High-Capacity cDNA Reverse Transcription Kit (Applied Biosystems). Transcripts of interest were quantified by real-time PCR carried out using LightCycler 480 SYBR Green I Master (Roche Life Science). All qPCRs were normalized to 16S rRNA gene expression. Primers used are listed in the Key resource table.

### Protein Expression and Lysate Experiments of 5β-reductase and 5α-reductase Genes

Candidate 5β-reductase and 5α-reductase genes from *B. dorei* DSM 17855 (Bacdor_03090 and Bacdor_03091) were cloned by Twist Biosciences into pET28a(+) vectors with an N-terminal His6 tag. The obtained constructs were transformed into BL21 (DE3) pLysS *E. coli* (Novagen), grown to saturation in LB medium supplemented with kanamycin (50 μg/mL) at 37 °C and diluted 1:100 into TB medium supplemented with kanamycin (50 μg/mL). The expression of the N-terminal His6 fusion proteins was induced at an OD600 of 0.5-0.6 with 400 μM or 50 μM isopropyl β-D-1-thiogalactopyranoside (IPTG) (Sigma-Aldrich, I5502-1G) for Bacdor_03090 and Bacdor_03091 respectively, and the induced cells were incubated at 18 °C for 20 hours. Cells from 200 mL of culture were pelleted by centrifugation (15 min at 4,200 rpm at 4 °C), resuspended in 20 mL of ice-cold phosphate-buffered saline (PBS) (Genesee Scientific, 25-507) containing 5% glycerol and 0.25 mM tris(2-carboxyethyl)phosphine hydrochloride (TCEP) (Sigma-Aldrich, C4706) and lysed by sonication. Cell debris was removed by centrifugation (20 min at 12,000g at 4 °C). Protein expression was confirmed in a qualitative western blot assay.

Bile acid substrate (3 μL of a 100 mM solution of 3-oxoLCA or 3-oxo-Δ4-LCA in DMSO) was added to 3 mL of clarified supernatant. After incubation at 37 °C for 24 hours, a 1 mL aliquot was removed for bile acid analyses (see below).

### Western Blot Analysis

Western blot of N-terminal His6 fusion Bacdor_03090 and Bacdor_03091 expression samples was performed using an 8% SDS-PAGE and transferred to a nitrocellulose membrane. The membrane was immunoblotted with a His-tag antibody (Cell Signaling Technology, 2325S) using a horseradish peroxidase (HRP)-conjugated secondary antibody (Cell Signaling Technology, 7074S). Total protein staining was performed using Amido Black 10B (Bio-Rad Laboratories) as a loading control.

### *In Vitro* Reconstitution of 3β-HSDH Activity

The candidate 3β-HSDH gene from *B. dorei* DSM 17855 (Bacdor_03089) was cloned by Twist Biosciences into the pET28a(+) vector with an N-terminal His6 tag. The obtained construct was transformed into BL21 (DE3) pLysS *E. coli* cells, grown to saturation in LB medium supplemented with kanamycin (50 μg/mL) at 37 °C and diluted 1:100 into TB medium supplemented with kanamycin (50 μg/mL). The expression of the N-terminal His6 fusion proteins was induced at an OD600 of 0.5-0.6 with 400 μM IPTG, and the induced cells were incubated at 18 °C for 20 hours. Cells from 3 L of culture were pelleted by centrifugation (12 min at 7,000g at 4 °C), resuspended in 40 mL of ice-cold PBS buffer containing 10% glycerol, 0.25 mM TCEP and 10 mM imidazole (Sigma-Aldrich, 56750), and lysed by sonication. Cell debris was removed by centrifugation (20 min at 16,000g at 4 °C). The supernatant was then mixed with preformed Ni-NTA for 1.5 hours at 4 °C. The nickel-bound protein was eluted with gradually increasing concentration of imidazole in PBS with 0.25 mM TCEP and 5% glycerol. Collected fractions were tested for purity by SDS-PAGE. The fractions containing Bacdor_03089 were combined and concentrated followed by dialysis using the storage buffer (PBS at pH 7.4 with 0.25 mM TCEP and 10% glycerol). 20 μM of 3-oxoalloLCA and 100 μM of β-nicotinamide adenine dinucleotide (NADH) (Sigma-Aldrich, N8129) was added to 1 mL of 1.0 μM protein. After incubation at 37 °C for 48 hours, a 1 mL aliquot was removed for bile acid analyses (see below).

### Bile Acid Analyses

Bile acid analyses were performed using a previously reported method (Yao et al., 2018). Stock solutions of all bile acids were prepared by dissolving compounds in molecular biology-grade DMSO (Sigma-Aldrich, D8418). These solutions were used to establish standard curves.

Glycocholic acid (GCA) (Sigma-Aldrich) was used as the internal standard. HPLC-grade solvents were used for preparing and running UPLC-MS samples. All data were analyzed using Agilent ChemStation.

### Sample Preparation for Bacterial Culture or Protein Expression Assay

Bacterial cultures or buffered solutions were quenched by acidification to pH = 1 using 6N HCl (Sigma-Aldrich, 258148). The resultant solutions were then extracted twice using 2 mL (for 1 mL aliquots) or 400 μL (for 200 μL aliquots) of ethyl acetate (Sigma-Aldrich, 319902). In case of an emulsion, the biphasic solution was centrifuged at 4,200 rpm for 10 min to obtain a clear separation. The combined organic extracts were then air dried and reconstituted in 50% methanol (EMD Millipore, MX0475) in dH2O for UPLC-MS analysis as we recently reported (Paik, Yao et al., 2020, under review).

### Sample Preparation for Cecal Contents

Bile acids were extracted from mouse cecal and faecal samples and quantified by UPLC-MS as previously reported (Yao et al., 2018). Briefly, tissue samples (approximately 250 mg) were pre-weighed in homogenizing tubes (Precellys lysing kit tough micro-organism lysing VK05 tubes) with ceramic beads. 400 μL methanol containing 10 μM internal standard (GCA) was added and thereafter homogenized (5,000 rpm for 90 s × 2, 6,500 rpm for 60 s, sample kept on ice between each run) and spun down for 20 min at 15,000 g. Of the supernatant, 200 μL was then transferred to a tube containing 200 μL of 50% methanol in dH2O followed by centrifugation for an additional 5 min at 15,000g. Of the supernatant, 50 μL was used for UPLC-MS analysis as previously described (Yao et al., 2018).

### Construction of *P. merdae* Knockout Mutants

The *P. merdae* Δ04016-18 mutant strain was constructed using a reported method with slight modifications (Garcia-Bayona and Comstock, 2019). Briefly, the 1 kb region upstream and downstream of Parmer_04016-18 was PCR-amplified, cloned into the pLGB13, and transformed into *E. coli* S17-1 λ pir chemical competent cells (Cullen et al., 2015; Degnan et al., 2014). *E. coli* S17 λ pir cells containing the desired plasmid were cultured aerobically in 3 mL of LB media at 3°C, and the recipient strain (*P. merdae* ATCC 43184) was cultured anaerobically in 3 mL BHI+ media at 37 °C. The *E. coli* S17 donor strain and *P. merdae* recipient strains were then subcultured in 25 mL or 5 mL of fresh media, respectively. At mid-to-late log growth, the cultures were spun down and the S17-1 λ donor strain was resuspended with the recipient strain culture in 100 μL BHI+ media. The resuspension was spotted directly onto a BHI+ plate and incubated aerobically at 37 °C agar-side down. After 16–24 hours, the bacterial biomass from the conjugation plates was scraped and resuspended in 1 mL PBS buffer, and then spread onto a BHI+ plate containing 200 μg/mL gentamicin and 10 μg/mL erythromycin in the anaerobic chamber. After confirmation of cointegrates via PCR, each strain was grown overnight in 3 mL BHI+ media, diluted 1:100 in fresh BHI+ media, and incubated for 6 to 8 hours. Aliquots of 50, 5, 1, and 0.5 μL were plated onto BHI+ plates containing 100 ng/mL anhydrotetracycline (aTC). After 2 to 4 days, single colonies were restreaked onto fresh BHI+ plates containing 100 ng/mL aTC. Knockout strain colonies were confirmed via PCR and sequencing. Loss of function of the knockout strain was confirmed via UPLC-MS with 100 μM 3-oxoLCA as the substrate. Primers used are listed in the Key resource table.

### *In Vitro* CD4+ T Cell Culture

Naive CD4+ T cells were purified from dissociated spleens and lymph nodes of C56BL/6N mice by fluorescence-associated cell sorting (FACS) (CD4+CD62L^hi^CD44^lo^). Purified cells were cultured under TH0 condition stimulated with anti-CD3e (5 μg/mL, clone 145-2C11, eBioscience), anti-CD28 (5 μg/mL, clone 37.51, eBioscience), and human IL-2 (100 U/mL, Peprotech). Additional TGFβ (0.25 ng/mL, Peprotech) was added for Treg polarization conditions. Bile acids, retinoic acid (1 nM, Sigma), or mitoParaquat (5 μM, Sigma) were added on day 0. IsoalloLCA dissolved in DMSO was resuspended in culture media and sonicated before being added to the culture. Cells were harvested either at 48 hours for RNA-Seq, ATAC-Seq, ChIP analysis, or at 72 hours followed by staining with anti-CD4 (RM4-5, eBioscience) or anti-Foxp3 (FJK-16s, eBioscience) antibodies, and analyzed with LSR II flow cytometer (BD). For mitoROS detection, cells cultured for 48 hours were incubated with 5 μM mitoSOX (ThermoFisher) for 30 min and assayed by flow cytometry. Flow cytometry data were acquired on an LSR II flow cytometer or Symphony flow cytometer (both BD) and data were analyzed with FlowJo software (TreeStar).

### ATAC-Seq and RNA-Seq Data Analysis

ATAC-Seq and RNA-Seq library construction and high-throughput sequencing were performed on approximately 20,000–50,000 sorted live cells, cultured for 48 hrs in vitro in the presence of DMSO, isoalloLCA (20 µM), or TGFβ (0.25 ng/mL).

ATAC-Seq reads were first filtered based on quality using sickle (default settings for either single-end or paired-end reads based on the library type; v1.33). Then the adapter sequences were trimmed using Cutadapt (-e 0.1, -m 20; v1.14). Reads were aligned to the mouse reference genome (mm10) using Bowtie 2 (v2.3.4.3). Mapped reads that had multiple alignments were removed using SAMtools (v1.3.1). PCR duplicates were removed by using Picard (v2.8.0; https://broadinstitute.github.io/picard/). The final output bam file was converted to bigwig format using deepTools (bamCoverage --normalized Using RPKM; v3.0.2) for visualization on the Integrative Genomics Viewer (IGV). To identify ATAC-Seq peaks for individual replicates, MACS2 (2.1.1.20160309) was used. Reproducible peaks among replicates were retained (IDR<0.1). DESeq2(v1.22.2) was then used for differential analysis of ATAC-Seq peaks to identify differential peaks in samples with various drug treatments and genetic backgrounds. Enriched transcription factor-binding motifs for different ATAC-Seq peak sets were identified using HOMER (findMotifsGenome.pl -size given -mask). PCA of ATAC-Seq data, differential peak heatmaps and statistical tests were performed using R (v3.5.2).

For RNA-Seq data analysis, low-quality reads were first removed by using sickle as described above. Reads were then mapped to the mm10 reference genome using the TopHat2 (v2.1.1) splice-junction mapper. Duplicated reads were removed using Picard. The HTSeq (htseq-count; v0.12.4) was used to count the number of reads overlapping genes. Transcripts per million (TPM) values of genes were calculated using kallisto (v0.45.1). DESeq2 was then used to identify differentially expressed genes (DEGs). To identify enriched gene sets and pathways associated with DEGs, we used clusterProfiler (v3.10.1).

### Chromatin Immunoprecipitation Analysis

*In vitro* cultured T cells treated with DMSO or isoalloLCA for 48 hours were harvested and fixed for 10 min with 1% formaldehyde. ChIP assays were performed according to standard protocol with Magna CHIP A/G kit (MilliporeSigma). Chromatin was immunoprecipitated with 3 µg of mouse IgG isotype control (Abcam) or NR4A1 antibody (12.14, eBioscience). The relative abundance of precipitated DNA fragments was analyzed by qPCR using SYBR Green Supermix (Bio-Rad). Primers are listed in the Key resources table.

### Luciferase Reporter Assay

HEK293T cells were transfected with a DNA mixture containing Foxp3-promoter-firefly luciferase reporter plasmid (WT-Luc), control *Renilla* luciferase (Promega pRL-CMV), and mouse NR4A1, NR4A2, and NR4A3 (Origene). Transfections were performed using TransIT-293 (Mirus) according to the manufacturer’s instructions. Luciferase activity was measured 24 hours later using the dual-luciferase reporter kit (Promega). The *Foxp3* promoter region was cloned into the pGL4.10 vector (Promega) as indicated in the figure legends. NR4A1 binding sites were predicted with JASPAR online server. The Mut_Luc construct harbors deletions in four predicted NR4A1 binding sites: GTAAAGGGCAAA, CAAAATTTCAAA, AGAAAGGCTACA and AAAAGGTATC.

### Statistical Analysis of IsoalloLCA Levels and Biosynthetic Genes in Human IBD Cohorts

#### Microbial Data Overview

We used two publicly available IBD metabolomics and metagenomics datasets for determining the differential abundance (DA) of isoalloLCA in disease/dysbiotic conditions, specifically 1) the Prospective Registry in IBD Study at MGH (PRISM) (Franzosa et al., 2019a) and 2) the IBDMDB study within the integrated Human Microbiome Project (HMP2 or iHMP) (Lloyd-Price et al., 2019).

PRISM is a cross-sectional cohort incorporating subjects diagnosed with Crohn’s disease (CD; n = 68); ulcerative colitis (UC, n = 53); and non-IBD controls (n = 34). PRISM stool samples were subjected to metabolomic profiling using a combination of four LC-MS methods, and the corresponding metabolome profiles were taken from the published paper (Franzosa et al., 2019a). Metagenomic sequencing reads from the paired 155 PRISM metagenomic samples were downloaded from SRA BioProject PRJNA400072 in April 2019. These reads had been previously processed by the KneadData workflow (http://huttenhower.sph.harvard.edu/kneaddata) which first trims the low-quality reads and bases, removes adapters, and finally, removes any human contaminant reads by mapping to the human genome (Franzosa et al., 2019a).

HMP2 is a longitudinal cohort containing 132 participants with CD (*n* = 67), UC (n = 38), and non-IBD controls (n = 27) followed for up to one year. We downloaded the HMP2 metagenomic sequencing reads and metabolomic profiles from the Inflammatory Bowel Disease Multi’omics Databases (http://ibdmdb.org) as of July 2020. These reads included 1,298 metagenomes samples and 546 metabolomes samples from 106 subjects (CD, n = 50; UC, n = 30; and non-IBD, n = 26). The metagenomic sequencing reads had been previously quality controlled by the KneadData workflow (as described above) Microbiome-associated IBD activity was previously defined in this cohort by comparing gut microbial composition of IBD patients to non-IBD controls (Lloyd-Price et al., 2019).

### Identifying Differentially Abundant Metabolites

We identified isoalloLCA in the HMP2 metabolomics dataset by running a pure standard of isoalloLCA in negative mode under the same LC-MS conditions used to collect HMP2 data and matching the *m/z* and chromatographic retention time of this known compound to a compound in HMP2: C18n_QI13593, m/z = 375.2905 at 11.04 min (Paik, Yao et al., 2020, under review). Differential abundance of isoalloLCA with respect to IBD diagnosis and dysbiosis state in the longitudinally sampled HMP2 cohort were determined using linear-mixed model models following our recent work (Paik, Yao et al., 2020, under review).

### Identifying Differentially Abundant Microbial Genes

We used ShortBRED (Kaminski et al., 2015) to accurately profile the abundance of genes involved in the isoalloLCA biosynthetic pathway in metagenomes from the PRISM (Franzosa et al., 2019a) and HMP2 (Lloyd-Price et al., 2019) datasets. In order to construct ShortBRED markers for these gene families, we first gathered homologs of 3β-HSDH, 5β-reductase, and 5α-reductase by performing BLASTP (Camacho et al., 2009) searches against the NCBI nr protein database (O’Leary et al., 2016) using 3β-HSDH, 5β-reductase, and 5α-reductase sequences from *B. dorei* DSM 17855, *P. merdae* ATCC 43184, and *B. vulgatus* ATCC 8489 as queries. Database sequences were selected for marker construction if they exhibited >80% query coverage, >50% alignment identity, and E-value < 10^-10^. These sequences were combined with UniRef90 (June 2020 version) as a comprehensive background for marker identification using ShortBRED-identify under default settings. We then applied the resulting marker sequences to profile the abundances of 3β-HSDH, 5β-reductase, and 5α-reductase from HMP2 and PRISM metagenomes using ShortBRED-quantify (also under default settings). Abundances are reported in RPKM units (read mapped per kilobase of coding sequence per million sample reads).

We tested the abundances of 3β-HSDH, 5β-reductase and 5α-reductase homologs for differential abundance over diagnosis and dysbiosis states following a very similar approach to the one introduced above in the context of HMP2 metabolomics. For the PRISM cohort, differential abundance over diagnosis was determined by evaluating the following linear model for each gene:

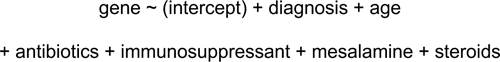

Diagnosis was coded as a categorical variable (CD, UC, non-IBD control) with non-IBD control as the reference state. Age was coded as a continuous covariate and four medication exposures (use of antibiotics, immunosuppressants, mesalamine, and steroids) were coded as binary covariates (with non-use as the reference state). Gene abundance values were zero-smoothed and log-transformed prior to linear model fitting within the MaAsLin 2 package (http://huttenhower.sph.harvard.edu/maaslin). The same random effects model formulation applied to HMP2 metabolomics was applied to HMP2 metagenomics within MaAsLin. Nominal *p*-values were adjusted for multiple hypothesis testing using the Benjamini-Hochberg method.

## SUPPLEMENTAL FIGURE LEGENDS

**Figure S1.**
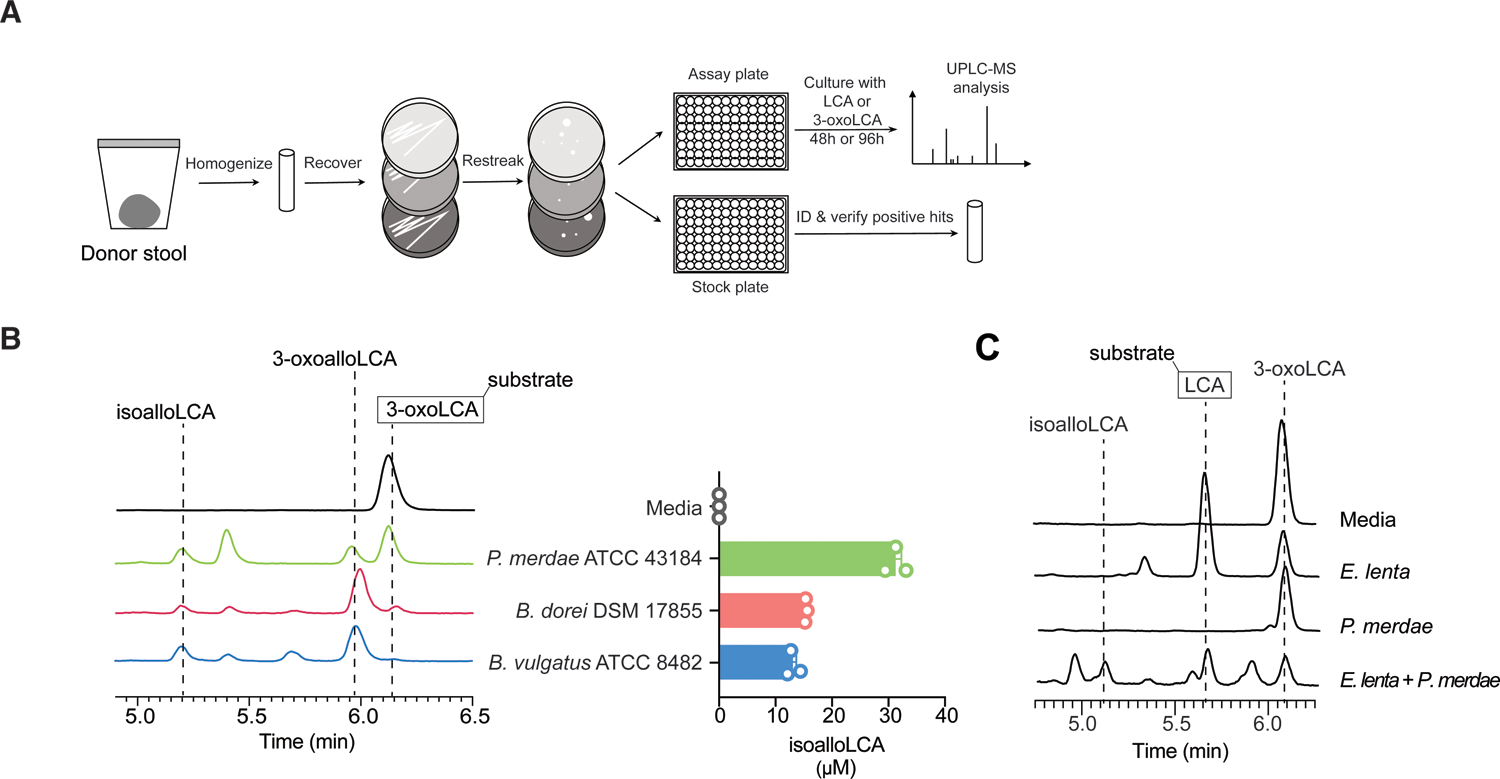
Type strains of gut bacteria produce isoalloLCA from the gut bacterial metabolite 3-oxoLCA. (A) Schematic representation of the screen of gut human bacterial isolates for isoalloLCA production. Bacterial strains were isolated from human stool as described previously (Paik, Yao et al., 2020, under review). The 990 bacterial strains isolated were then incubated with either 100 μM LCA or 3-oxoLCA. IsoalloLCA-producing strains were identified by UPLC-MS analysis, and 16S rRNA sequencing was performed on these positive hits. (B) Representative UPLC-MS traces (left) and quantification of isoalloLCA production (right) showing that type strain *P. merdae* ATCC 43184, *B. dorei* DSM 17855 and *B. vulgatus* ATCC 8482 metabolize 3-oxoLCA to isoalloLCA. Aliquots were removed for UPLC-MS analysis following 96 h of incubation. (n = 3 biological replicates per group, data are shown as the mean ± SEM, N.D. = not detected). (C) Representative UPLC-MS traces showing that a co-culture of *Eggerthella lenta* DSM 2243 and *Parabacteroides merdae* ATCC 43184 converted LCA to isoalloLCA while monocultures of these bacteria did not produce isoalloLCA.

**Figure S2.**
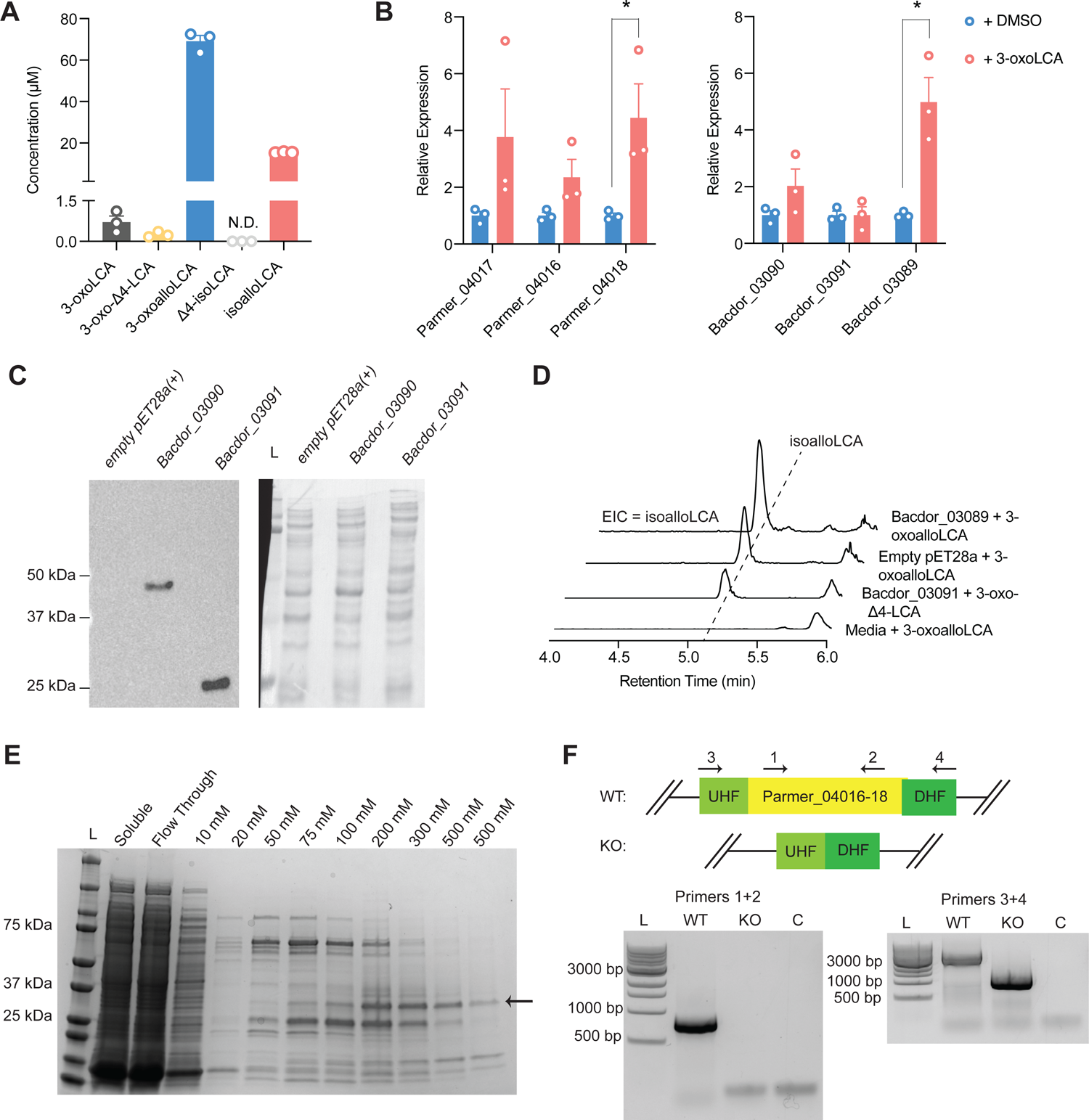
Identification of an inducible gene cluster in Bacteroidetes species that converts 3-oxoLCA to isoalloLCA. (A) The bile acid metabolites 3-oxo-Δ4-LCA, 3-oxoalloLCA, and isoalloLCA were detected by UPLC-MS in cultures of *B. dorei* DSM 17855 incubated with 100 μM 3-oxoLCA for 96 hrs, while Δ4-isoLCA was not detected (n = 3 biological replicates per group, data are shown as the mean ± SEM, N.D. = not detected). (B) As measured by RT-qPCR, the expression of genes in the Bacteroidetes was higher in cultures treated with 100 μM 3-oxoLCA for 1 hr compared to DMSO-treated cultures (n = 3 biologically independent samples per group, data are shown as the mean ± SEM by Welch’s t-test with 2-tailed *p*-value, * *p* < 0.05). (C) Western blot against anti-His tag antibodies and then stained with Amido black showing induction of N-terminal His6-Bacdor_03090 and N-His6-Bacdor_03091 in *E. coli* BL21. N-His6-Bacdor_03090 has a molecular weight of ∼45.3 kDa, and N-His6-Bacdor_03091 has a molecular weight of ∼28.2 kDa. L = Ladder using Precision Plus Protein Dual Color Prestained Protein Standard. (D) Representative UPLC-MS traces showing the formation of isoalloLCA from 3-oxoalloLCA in *E. coli* BL21 cell lysate. EICs for isoalloLCA (m/z 375) are shown. (E) SDS-PAGE showing purification of N-terminal His6-Bacdor_03089. N-His6-Bacdor_03089 has a molecular weight of ∼31.7 kDa. L = Ladder using Precision Plus Protein Dual Color Prestained Protein Standard. Soluble = clarified, soluble supernatant fraction from whole cell lysate; Flow Through = resulting unbound flow-through fraction after application to Ni-NTA column; X mM = washes/elutions of increasing imidazole concentrations. Arrow indicates predicted N-His6-Bacdor_03089 protein. (F) PCR verification of the counter-selected clones for deletion of the 5.23 kb Parmer_04016-04018 region. WT, *P. merdae* ATCC 43184 wild type; KO, *P. merdae* ATCC 43184 ΔParmer_04016-04018; C, no-DNA control; L = Ladder using NEB 1 kb DNA ladder. (Left) PCR using primers 1 and 2, expect 887 b for WT, 0 b for KO; (Right) PCR using primers 3 and 4, expect 4.13 kb for WT, 1.27 kb for KO.

**Figure S3.**
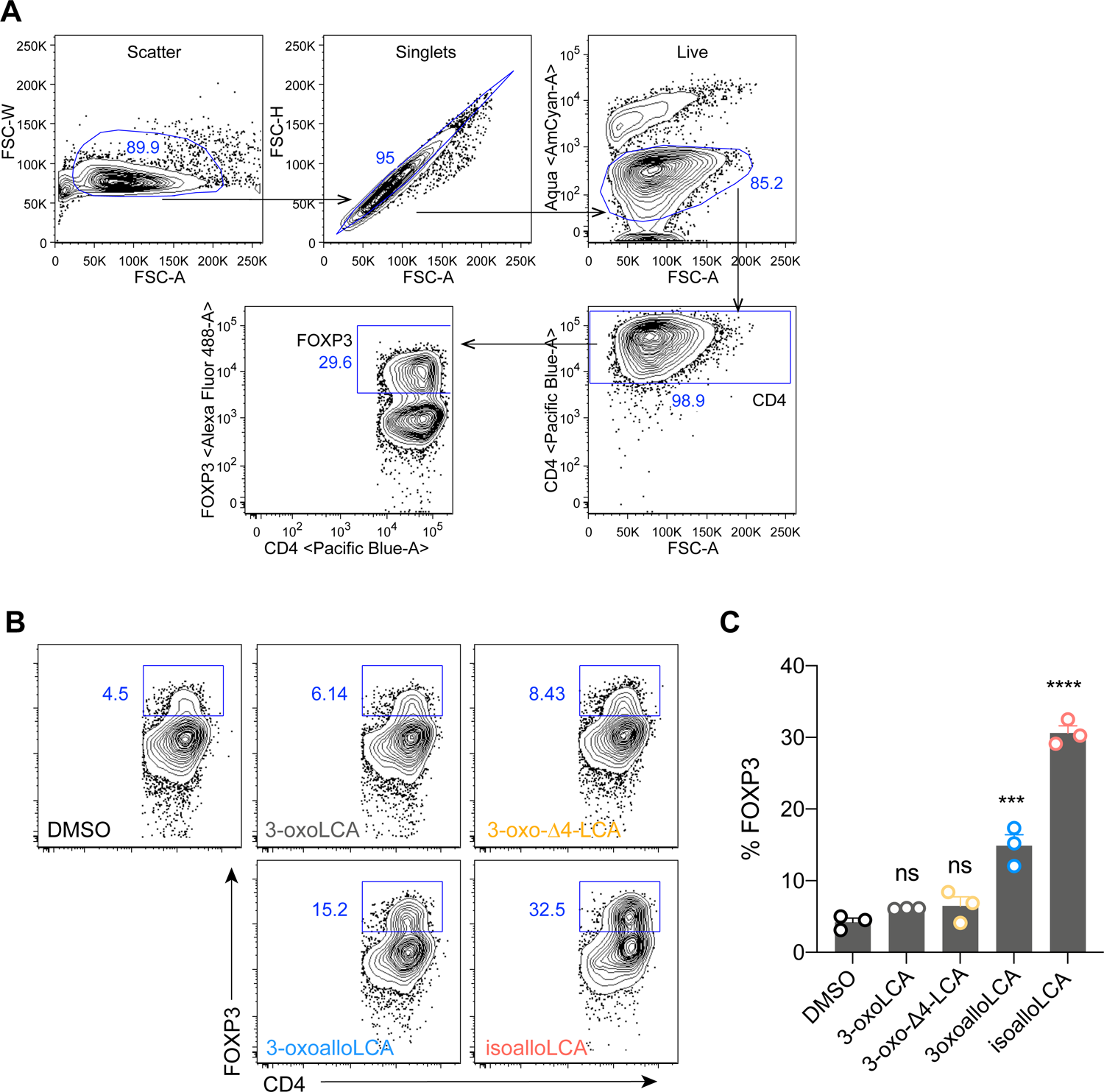
Effects of the bile acid intermediates of isoalloLCA biosynthetic pathway on Treg differentiation *in vitro*. (A) Gating strategy of the flow cytometric analysis of *in vitro* cultured T cells. (B-C) Flow cytometry analysis and quantification of naïve CD4^+^ T cells purified from WT-B6 mice, culture under Th0 conditions (anti-CD3, anti-CD28, IL-2) in the presence of DMSO, 3-oxoLCA (20 µM), 3-oxo-Δ4-LCA (20 µM), 3-oxoalloLCA (20 µM), or isoalloLCA (20 µM) for 72 hours. Wells were stained with FOXP3 as a marker for Treg cells (n = 3 biological replicates per group, data are shown as the mean ± SEM, one-way ANOVA with Dunnett’s multiple comparison test, compared to the DMSO group, ns, not significant, *** *P* < 0.001, **** *P* < 0.0001).

**Figure S4.**
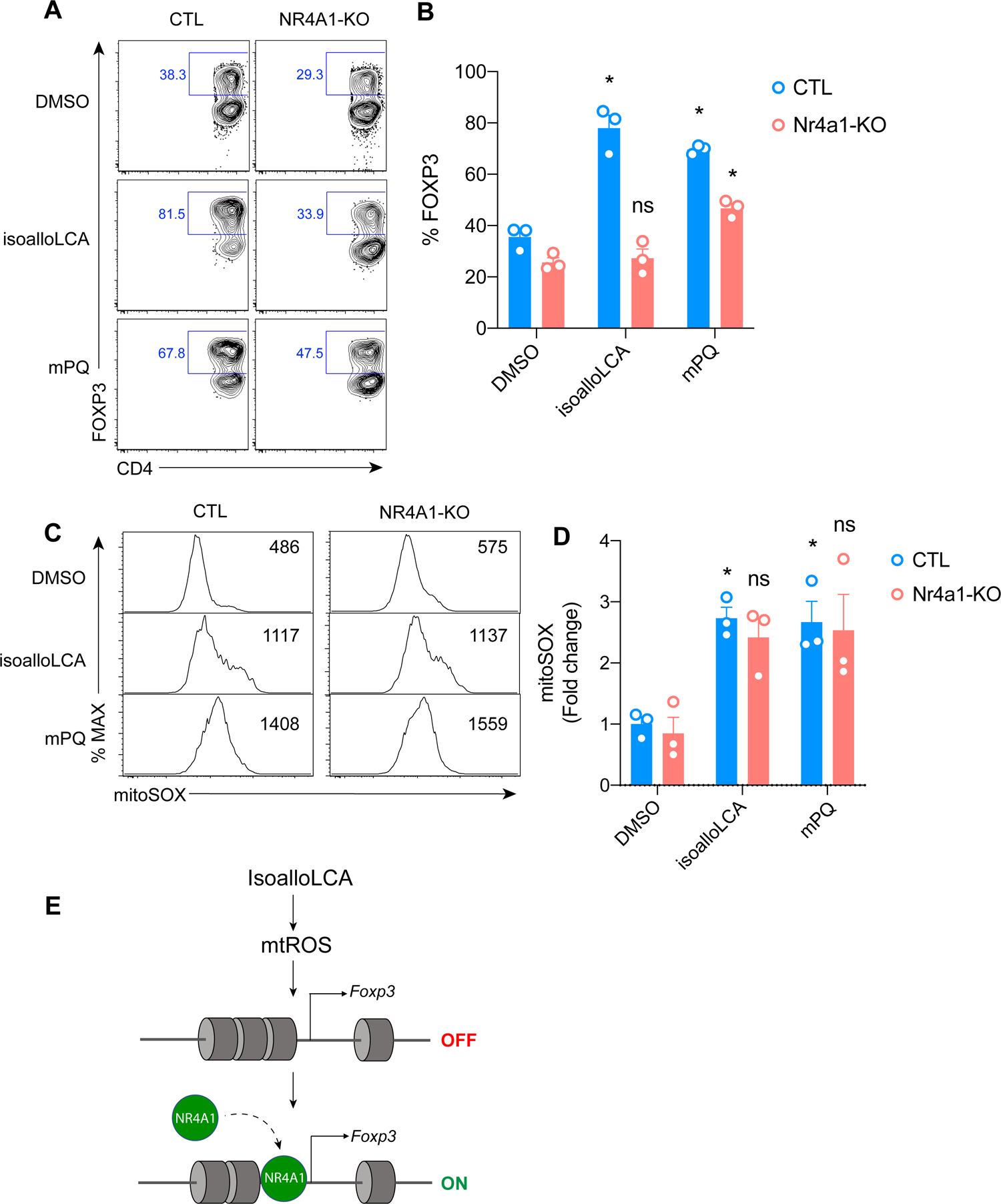
NR4A1 deficiency does not affect isoalloLCA-induced mitochondrial ROS production. (A-B) Flow cytometry and quantification of CD4^+^ T cells stained intracellularly for FOXP3. Naive CD4^+^ T cells isolated from WT (CTL) or *Nr4a1*-KO mice were cultured under Treg polarizing conditions (anti-CD3, anti-CD28, IL-2 and TGFβ) in the presence of DMSO, isoalloLCA (20 µM), or mPQ (5 µM) for 72 hours (n = 3 biological replicates per group, data are shown as the mean ± SEM, two-way ANOVA with Tukey’s multiple comparison test, each group was compared to the DMSO group, ns, not significant, * *P* < 0.05). (C-D) Flow cytometry and quantification of mitochondrial ROS production measured by mitoSOX with cells cultured under Treg polarizing conditions (anti-CD3, anti-CD28, IL-2 and TGFβ) in the presence of DMSO, isoalloLCA (20 µM), or mPQ (5 µM) for 48h (n = 3 biological replicates per group, data are shown as the mean ± SEM, two-way ANOVA with Tukey’s multiple comparison test, each group was compared to the DMSO group, ns, not significant, * *P* < 0.05). (E) Model depicting the mechanism by which isoalloLCA and NR4A1 enhance Treg differentiation. IsoalloLCA-induced mtROS production leads to chromatin modification, including acetylation of H3K27. Increased chromatin accessibility at the *Foxp3* promoter exposes the NR4A1 binding sites. Thus, increased association of NR4A1 results in activation of *Foxp3* gene transcription.

**Figure S5.**
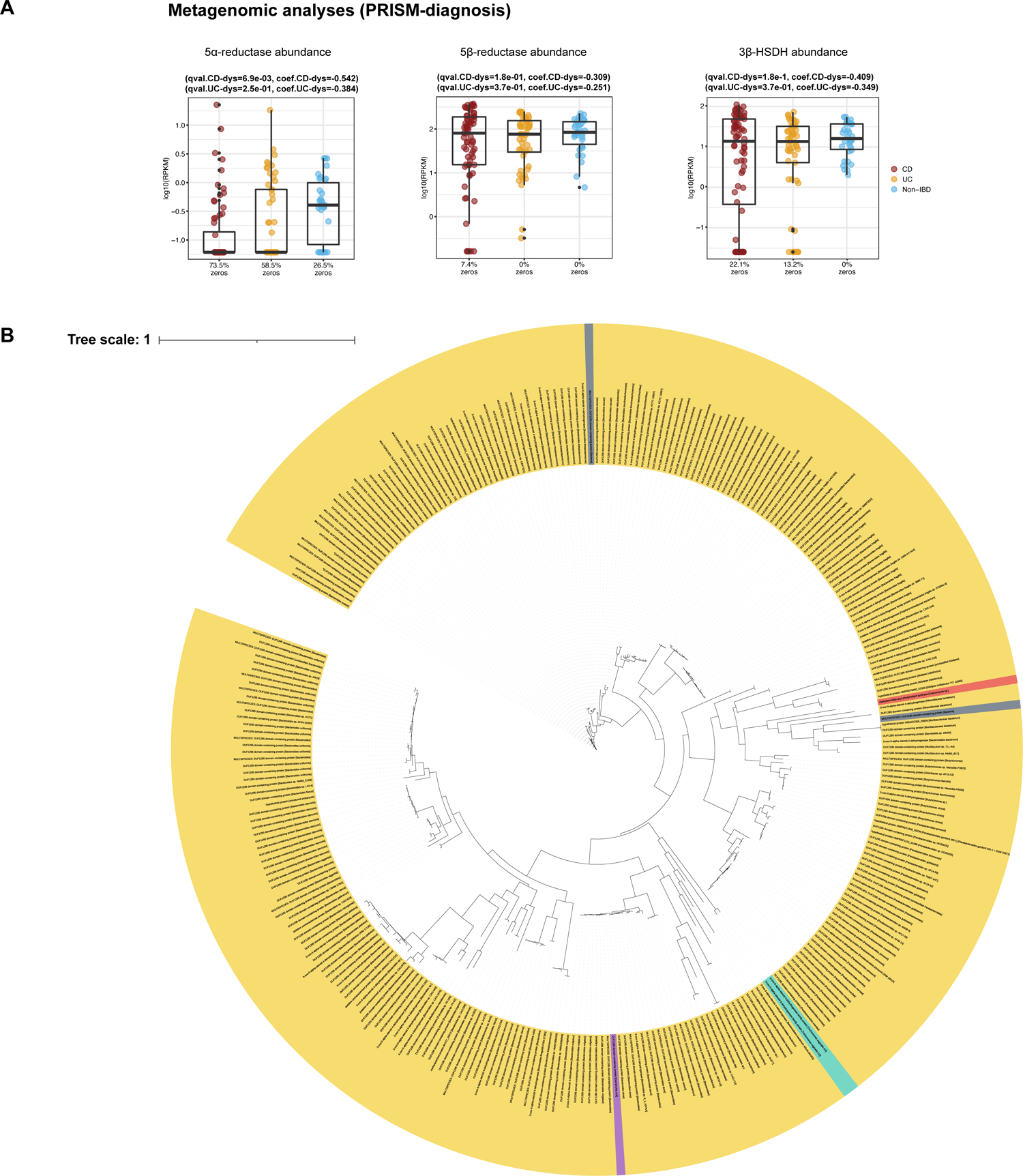
Effects of isoalloLCA-producing bacteria in mice and associations in humans. (A) The abundance of 5α-reductase homolog was significantly depleted in CD patients compared to nonIBD controls in PRISM, while no significant differential abundance was identified for 5β-reductase and 3β-HSDH homologs (*n* = 155 samples from 68 participants with CD, 53 with UC and 34 nonIBD controls; linear model, FDR-adjusted two-tailed *p*-value < 0.05). Boxplots show median and lower/upper quartiles with outliers outside of boxplot ‘whiskers’ (indicating the inner fences of the data). The percentage of zeros in each condition are added as x-axis tick labels. (See Table S6 for full results.) (B) Maximum likelihood phylogenetic tree for 5α-reductase homologs identified by querying the NCBI nucleotide collection for Parmer_04016 and Bacdor_03091. Homologs are largely found in the Bacteroidetes phylum. (Yellow: Bacteroidetes, Purple: Proteobacteria, Grey: Unclassified bacteria, Green: Metamonada in Eukaryota, Red: Neocallimastigomycota in Fungi) Sequences were aligned using MUSCLE (standard settings), and a maximum likelihood tree was created using FastTree (standard settings, 20 rate categories of sites). The tree files were uploaded to the Interactive Tree of Life web server (https://itol.embl.de/) to annotate the trees (Letunic and Bork, 2016).

## Supplementary Information

### Synthetic Procedures

#### General

All reagents were obtained commercially unless otherwise noted. All anhydrous reactions were run under an atmosphere of argon or nitrogen. Anhydrous tetrahydrofuran (THF) was purchased from Sigma Aldrich. Silica gel column chromatography was performed using 60 Å silica gel (230−400 mesh). Thin layer chromatography was performed on EMD Millipore TLC silica gel 60 F254 plates (250 μm). Visualization of the developed chromatogram was accomplished by fluorescence quenching and by staining with aqueous p-anisaldehyde. Nuclear magnetic resonance (NMR) spectra were acquired on a Varian MR spectrometer operating at 400 and 100 MHz for ^1^H and ^13^C respectively, and Bruker Advance II spectrometer operating at 600 and 150 MHz for ^1^H and ^13^C respectively, and are referenced internally according to residual solvent signals. Data for ^1^H NMR are recorded as follows: chemical shift (δ, ppm), multiplicity (s, singlet; d, doublet; t, triplet; q, quartet; m, multiplet; br, broad), integration, coupling constant (Hz). Data for ^13^C NMR are reported in terms of chemical shift (δ, ppm). High resolution mass spectra were obtained using an Agilent 6530 Quadrupole Time of Flight (Q-TOF) mass spectrometer.

Experimental protocols and characterization data:

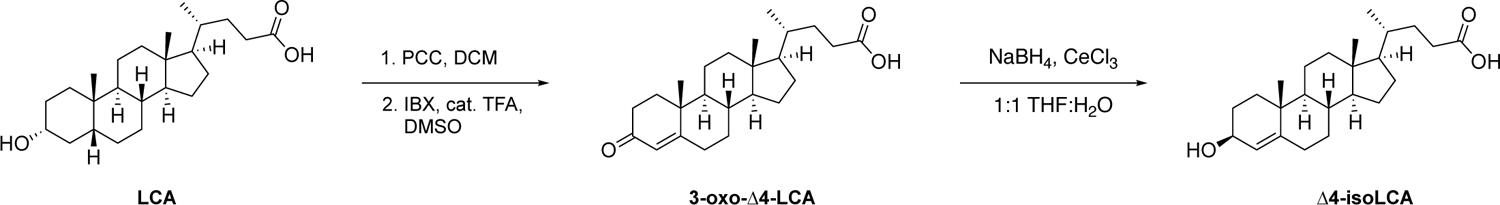

**Scheme 1.**
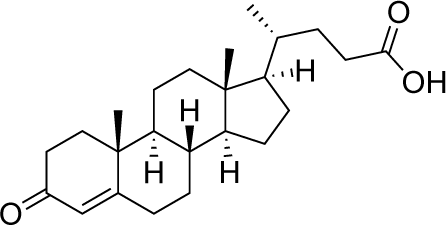
Synthesis of 3-oxo-4Δ-LCA and 4Δ-isoLCA.

## 3-oxo-4Δ-LCA

<u>Step 1.</u> A protocol reported in the literature was followed for this step of the synthesis – *Nat. Chem. Biol.*, **2020**, *16*, 318-326.

To a suspension of pyridinium chlorochromate (PCC) (TCI Chemicals, P0930) (8.73 g, 40.48 mmol) and silica gel (57.25 g) in 200 mL DCM at room temperature (rt) was added lithocholic acid (5.0 g, 13.28 mmol) slowly. The resulting solution was stirred at rt for 3 h. The reaction mixture was then filtered through a bed of Celite and the residue was concentrated under reduced pressure to provide the crude 3-oxolithocholic acid. The crude compound was then purified by silica gel chromatography (20% ethyl acetate/90%hexanes to 60% ethyl acetate/40% hexanes) to provide pure 3-oxolithocholic acid (1.97 g, 40%) as a white powder.

<u>Step 2.</u> A protocol reported in the literature was adapted for this step of the synthesis – *Tetrahedron Lett.*, **2011**, *52 (32)*, 4137-4139.

To a suspension of 3-oxolithocholic acid (449.7 mg, 1.21 mmol) and 2-iodoxybenzoic acid^1^ (790.0 mg, 2.82 mmol) in 6.5 mL DMSO at rt was added trifluoroacetic acid (27.7 µL, 0.35 mmol) dropwise. The resulting solution was stirred at 40 °C for 4 h. The mixture was then cooled to rt and diluted with 1 mL ethyl acetate and 1 mL saturated sodium bicarbonate solution. The organic layer was separated and the aqueous layer was extracted with ethyl acetate (3 x 10 mL). The combined organic layers were then dried over sodium sulfate, filtered and concentrated under reduced pressure. The crude compound was then purified by silica gel chromatography (20% ethyl acetate/80% hexanes to 80% ethyl acetate/20% hexanes) to provide the target compound 3-oxo-Δ4-LCA (82.1 mg, 18%) as a white powder.

### 3-oxo-4Δ-LCA

TLC R*f* = 0.38 (Benzene; 1,4-Dioxane; Acetic Acid, 90:9:1 v/v); ^1^H NMR (400 MHz, CDCl3): δ 5.73 (s, 1H), 2.45-2.21 (m, 6H), 2.03-1.99 (m, 2H), 1.90-1.77 (m, 3H), 1.71-1.23 (m, 9H), 1.16-0.97 (m, 8H), 0.91-0.88 (m, 4H), 0.70 (s, 3H); ^13^C NMR (100 MHz, CDCl3): δ 199.95, 179.80, 171.99, 123.66, 55.80, 55.72, 53.72, 42.41, 39.55, 38.57, 35.59, 35.56, 35.23, 33.84, 32.90, 31.97, 31.02, 30.70, 27.98, 24.10, 20.98, 18.19, 17.33, 11.94; HRMS (*m/z*): [M - H]^-^ calcd for C24H35O3, 371.2591, found 371.2605.

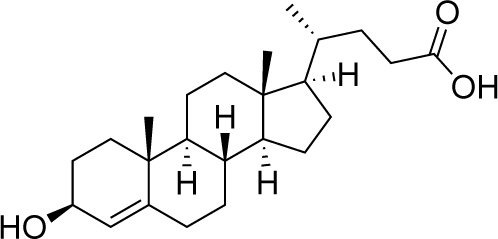

### 4Δ-isoLCA

To a solution of cerium chloride (58.3 mg, 0.16 mmol) and 3-oxo-Δ4-lithocholic acid (22.3 mg, 0.06 mmol) in 4 mL 1:1 THF:water at rt was added sodium borohydride (21.8 mg, 0.58 mmol) slowly. The resulting solution was stirred under nitrogen at rt for 30 min. The reaction mixture was then quenched with 4 mL water and 4 mL ethyl acetate. The organic layer was separated and the aqueous layer was extracted with ethyl acetate (3 x 5 mL). The combined organic layers were then dried over sodium sulfate, filtered and concentrated under reduced pressure. The crude compound was then purified by injecting onto a Luna^®^ 5 µm, C18(2) 100 Å, 250 x 4.6 mm LC column at room temperature and was eluted using a 55 min method of 55% A to 45% B (A = acetonitrile + 0.1% formic acid; B = water + 0.1% formic acid). Fractions were collected and concentrated to provide the target compound Δ4-isoLCA (2.6 mg) as a white powder.

### Δ4-isoLCA

TLC R*f* = 0.31 (Ethyl acetate:dichloromethane, 50:50 v/v); ^1^H NMR (600 MHz, CD3OD): δ; 5.24 (s, 1H), 4.07-4.04 (t, 1H, 12 Hz), 2.24-2.19 (m, 2H), 2.07-1.99 (m, 3H), 1.93-1.85 (m, 2H), 1.82-1.70 (m, 3H), 1.62-1.57 (m, 1H), 1.50-1.26 (m, 10H), 1.16-1.08 (m, 3H), 1.07 (s, 3H), 1.01-0.97 (m, 1H), 0.95-0.94 (m, 3H), 0.90-0.83 (m, 1H), 0.75-0.73 (m, 1H), 0.72 (s, 3H); ^13^C NMR (150 MHz, CD3OD): δ 168.86, 146.55, 123.31, 67.07, 56.16, 54.68, 42.24, 39.82, 36.96, 35.92, 35.71, 35.35, 34.22, 32.99, 32.37, 31.91, 28.49, 27.73, 23.82, 20.71, 17.93, 17.47, 11.00; HRMS (*m/z*): [M - H2O + H]^+^ calcd for C24H38O3, 357.2793, found 357.2787.

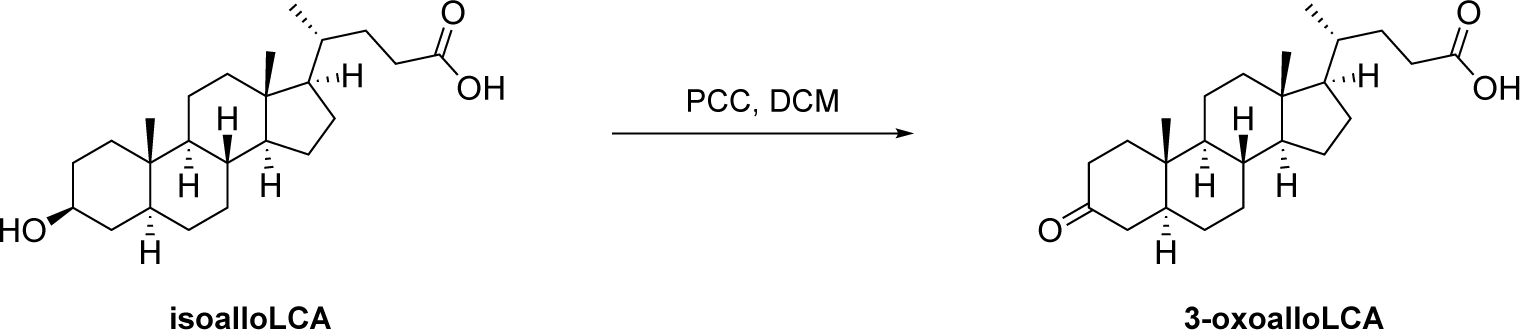

**Scheme 2.**
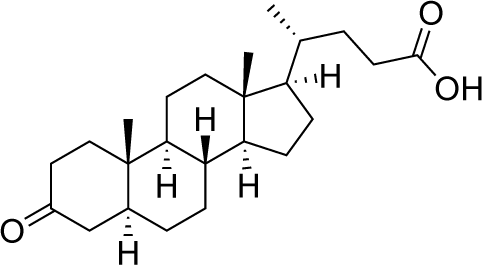
Synthesis of 3-oxoalloLCA.

### 3-oxoalloLCA

A protocol reported in the literature was followed for this step of the synthesis – *Nat. Chem. Biol.*, **2020**, *16*, 318-326.

To a suspension of PCC (33.9 mg, 0.16 mmol) and silica gel (218 mg) in 400 µL DCM at rt was added isoallolithocholic acid (20.0 mg g, 0.05 mmol) slowly. The resulting solution was then stirred at rt for 23 h. The reaction mixture was then filtered through a bed of Celite and the residue was concentrated under reduced pressure to provide the crude 3-oxolithocholic acid. The crude compound was then purified by silica gel chromatography (10% ethyl acetate/90%hexanes to 80% ethyl acetate/20% hexanes) to provide pure 3-oxolithocholic acid (9.4 mg, 47%) as a white powder.

### 3-oxoalloLCA

TLC R*f* = 0.84 (Ethyl acetate:Hexanes, 50:50 v/v); ^1^H NMR (400 MHz, CDCl3): δ 2.43-2.22 (m, 5H), 2.10-1.95 (m, 3H), 1.90-1.76 (m, 2H), 1.71-1.67 (m, 1H), 1.62-1.25 (m, 12H), 1.18-1.00 (m, 7H), 0.93-0.92 (m, 2H), 0.89-0.83 (m, 1H), 0.76-0.72 (m, 1H), 0.68 (s, 3H); ^13^C NMR (100 MHz, CDCl3) δ 212.18, 179.38, 56.23, 55.87, 53.75, 46.67, 44.68, 42.64, 39.85, 38.54, 38.15, 35.62, 35.38, 35.28, 31.67, 30.85, 30.73, 28.93, 28.06, 24.16, 21.42, 18.22, 12.06, 11.44.; HRMS (*m/z*): [M - H]^-^ calcd for C24H37O3, 373.2748, found 373.2738.

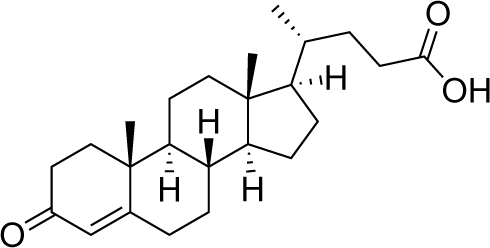

## 3-oxo-4Δ-LCA

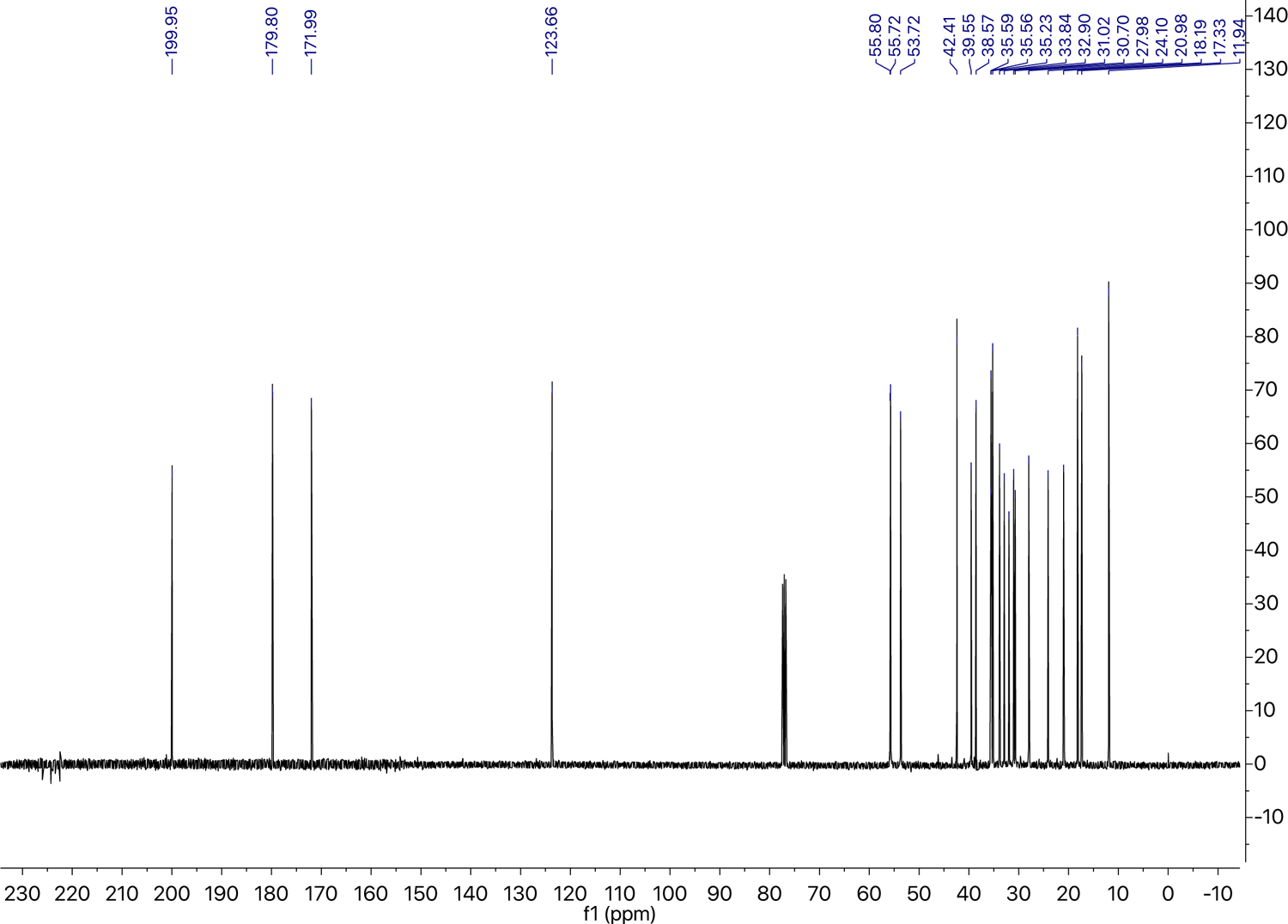
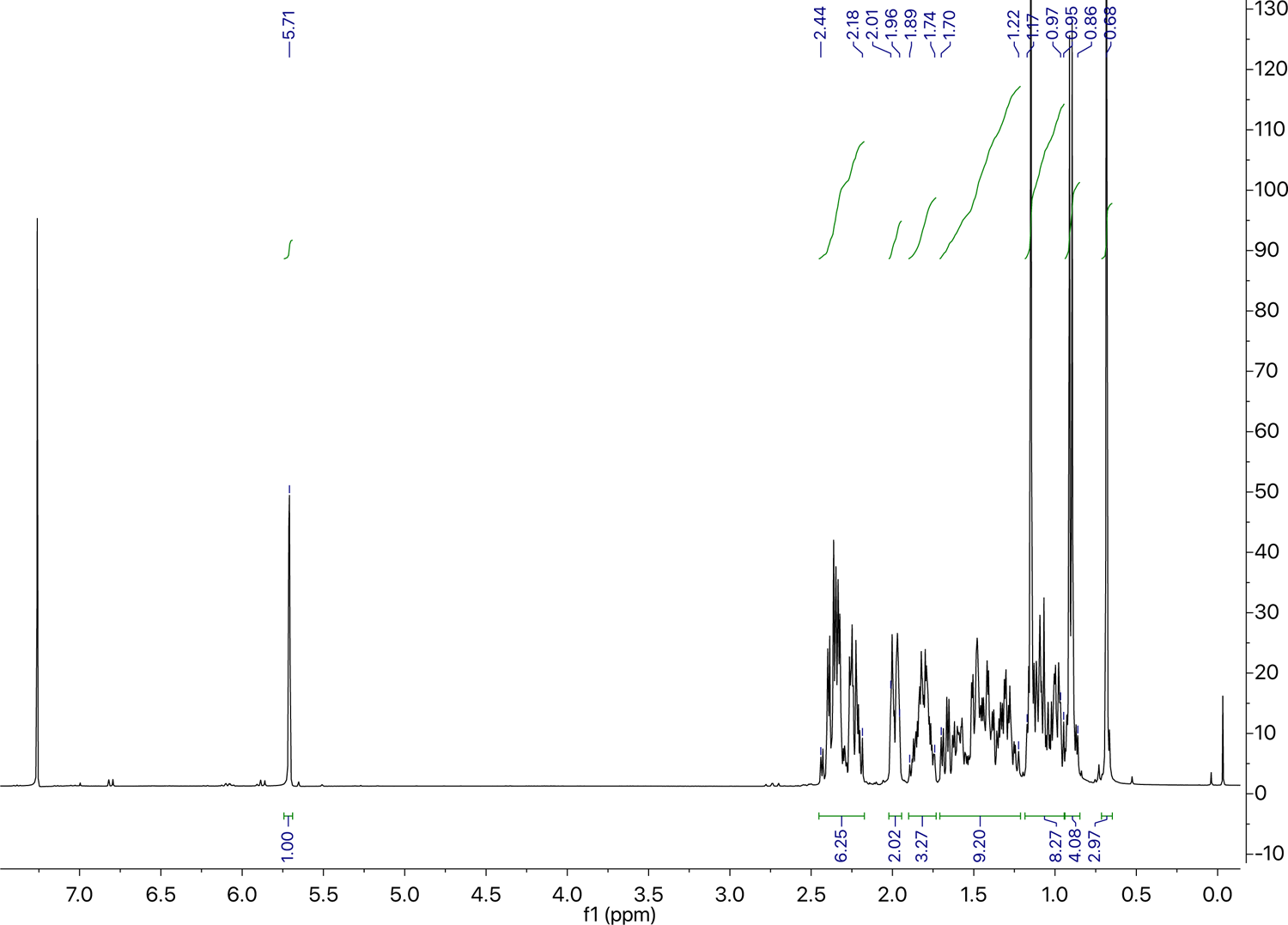
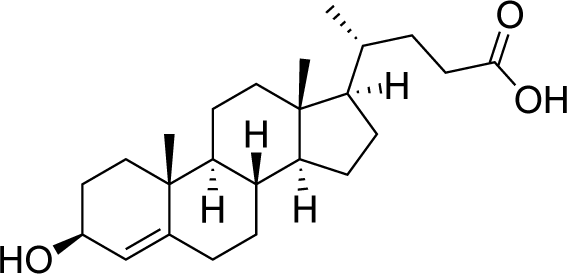

### 4Δ-isoLCA

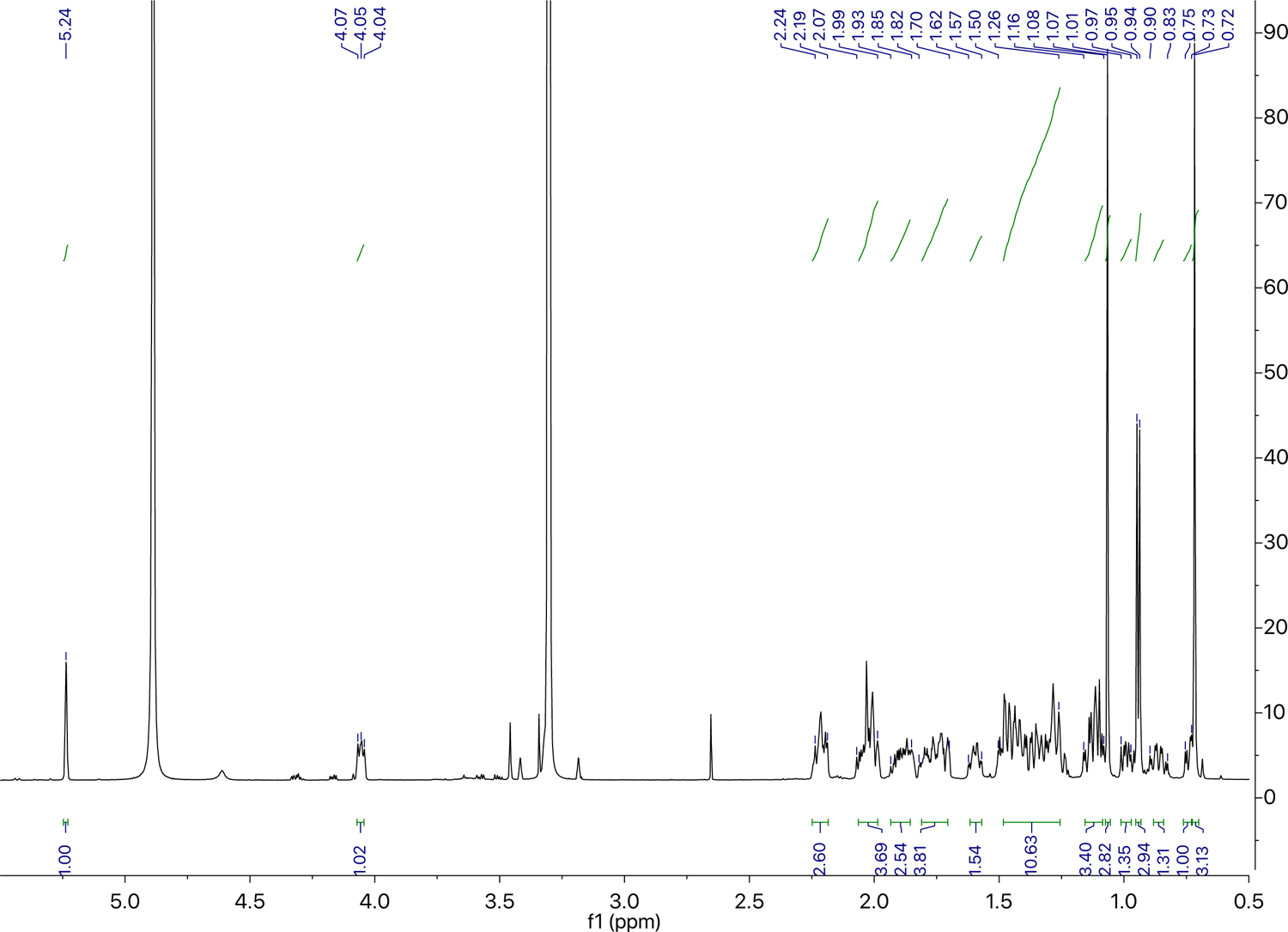
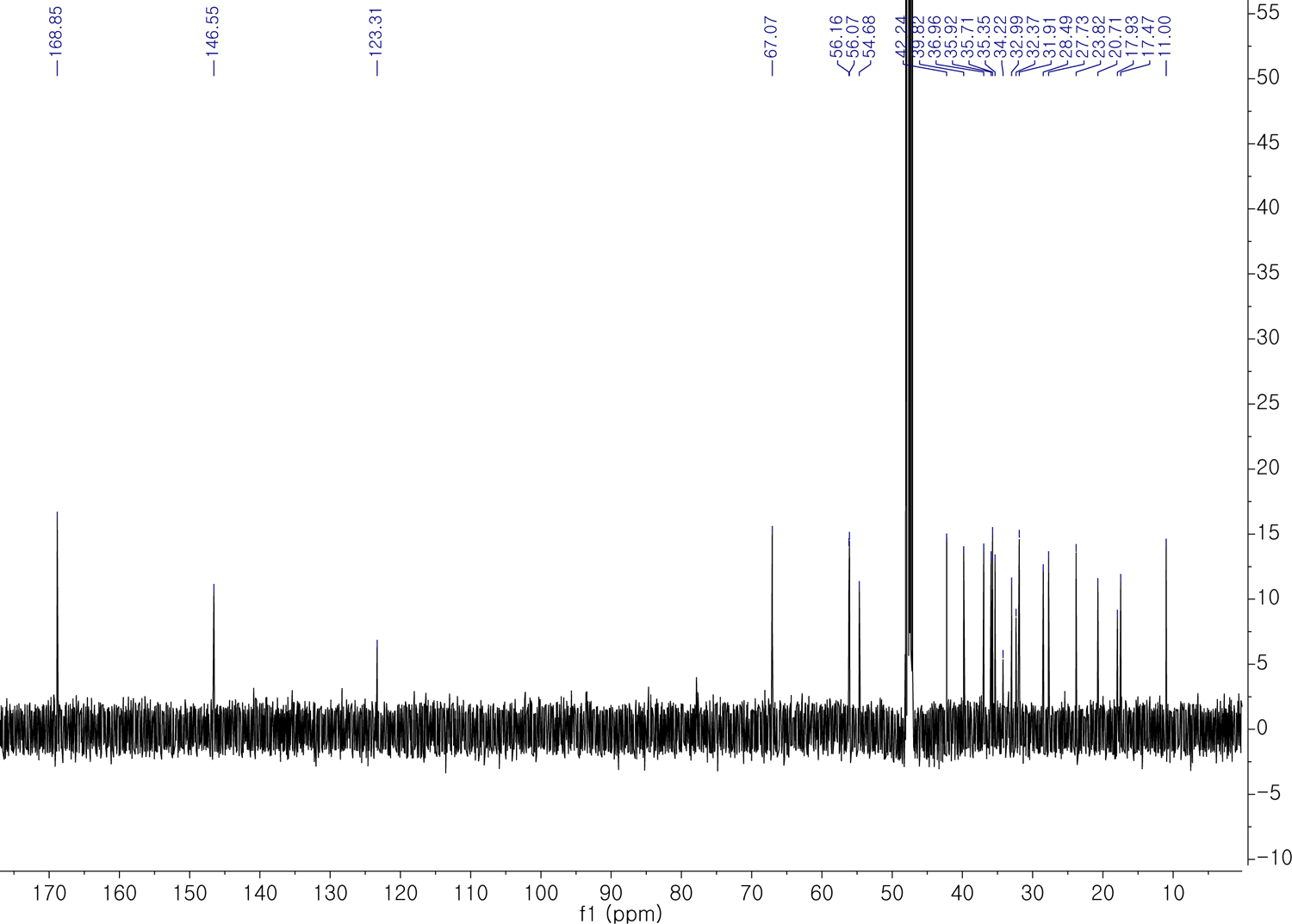
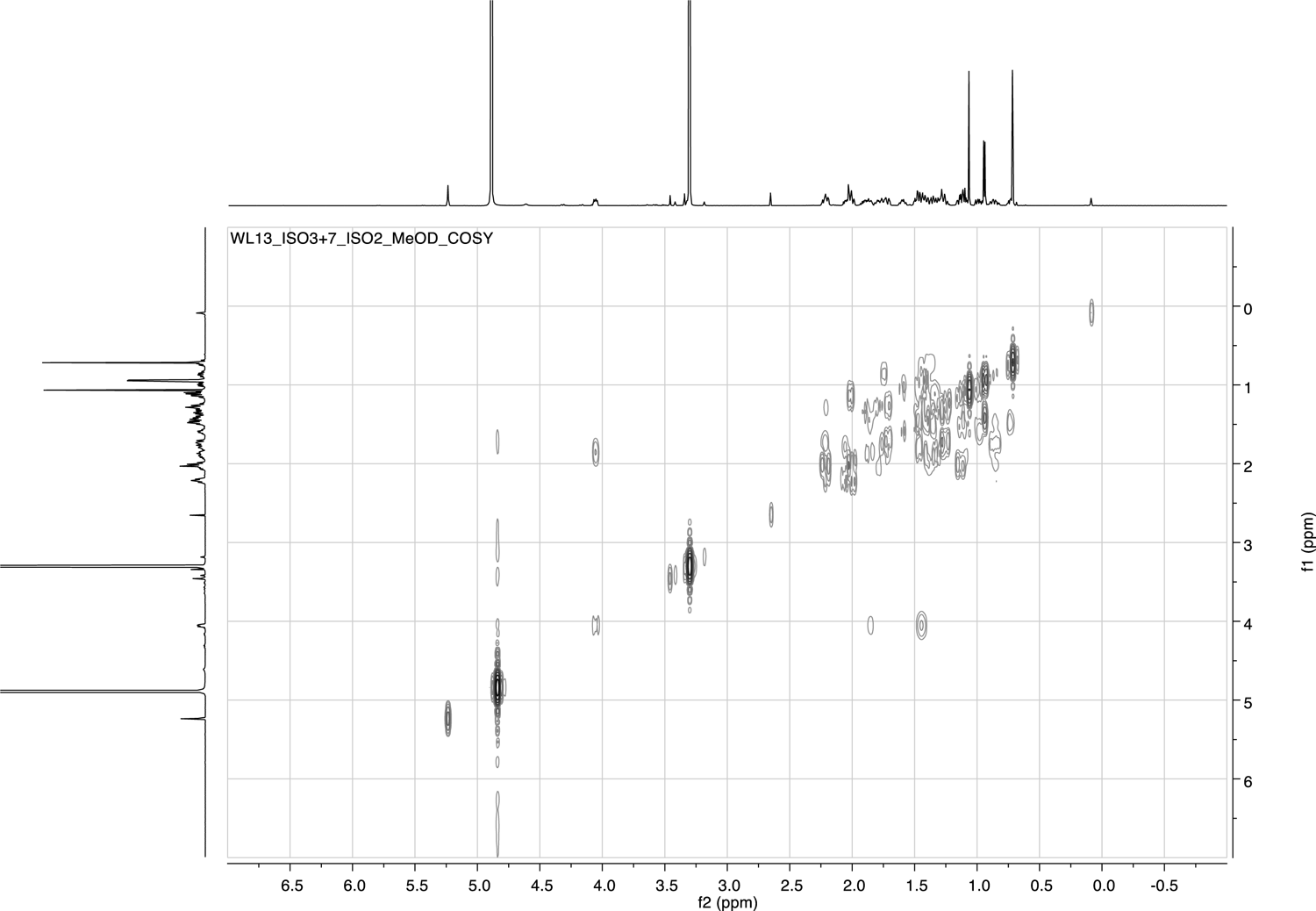
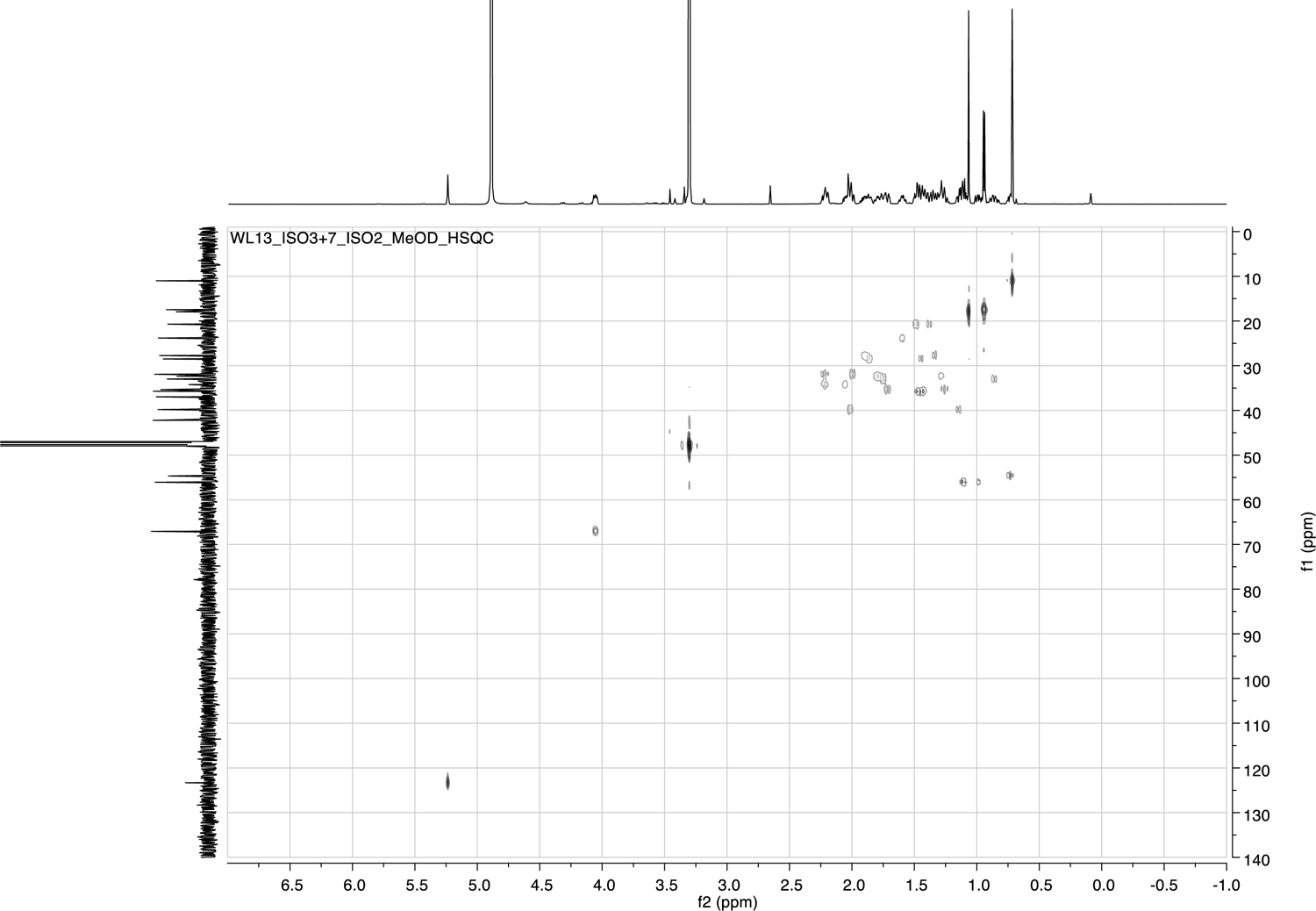
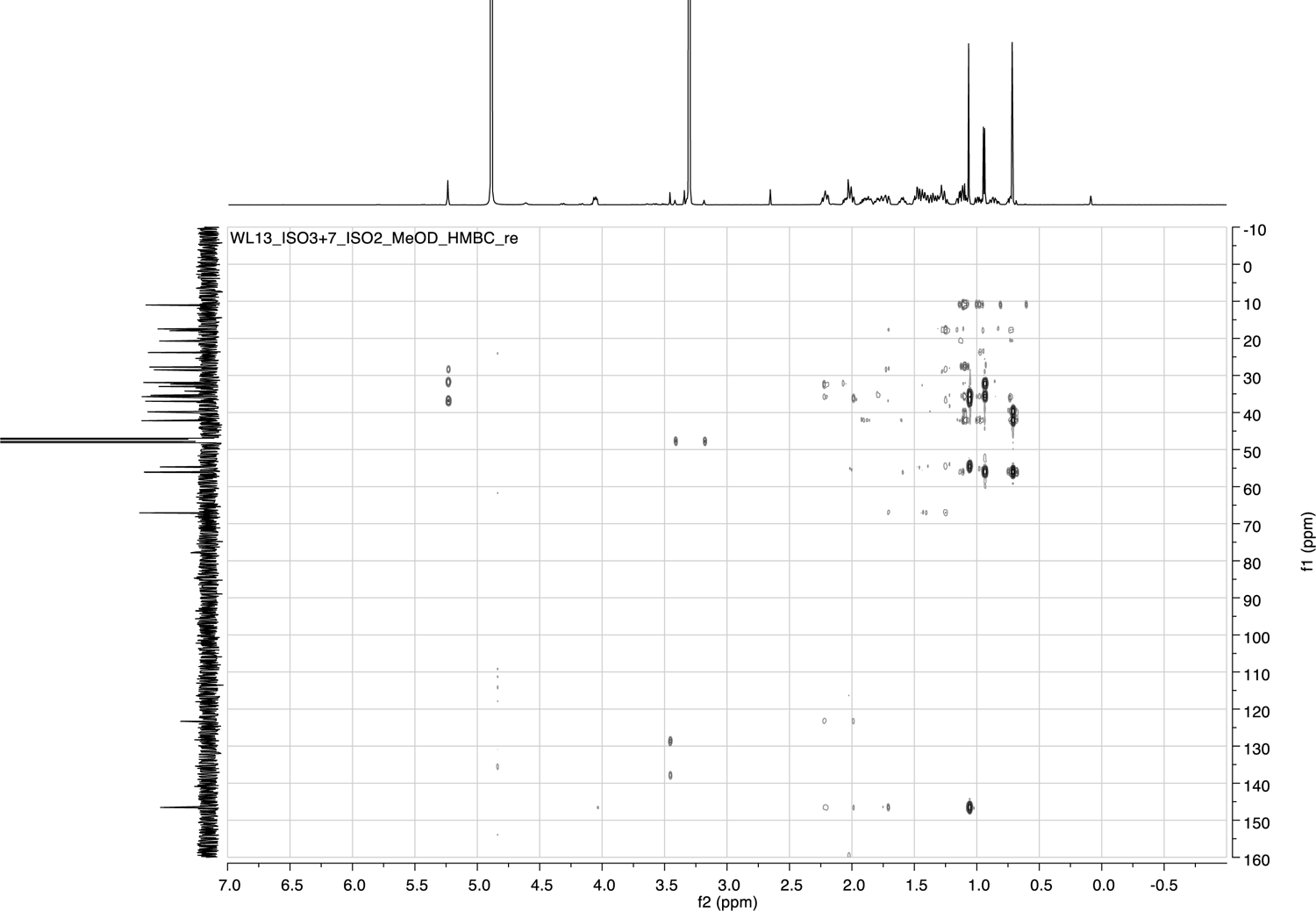
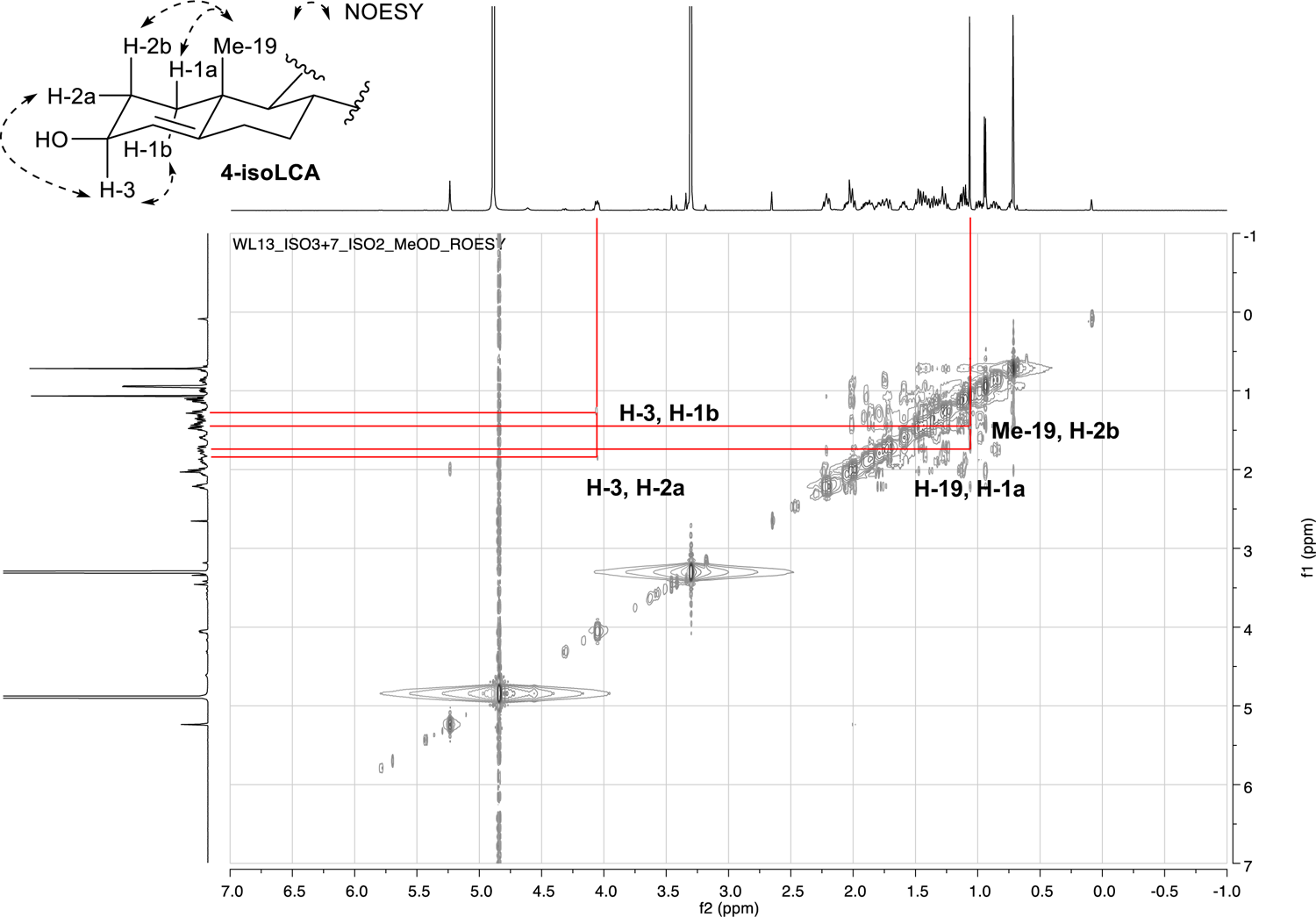
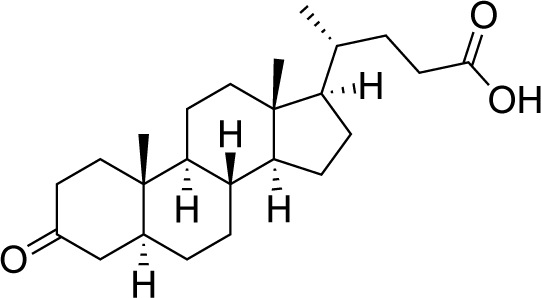

### 3-oxoalloLCA

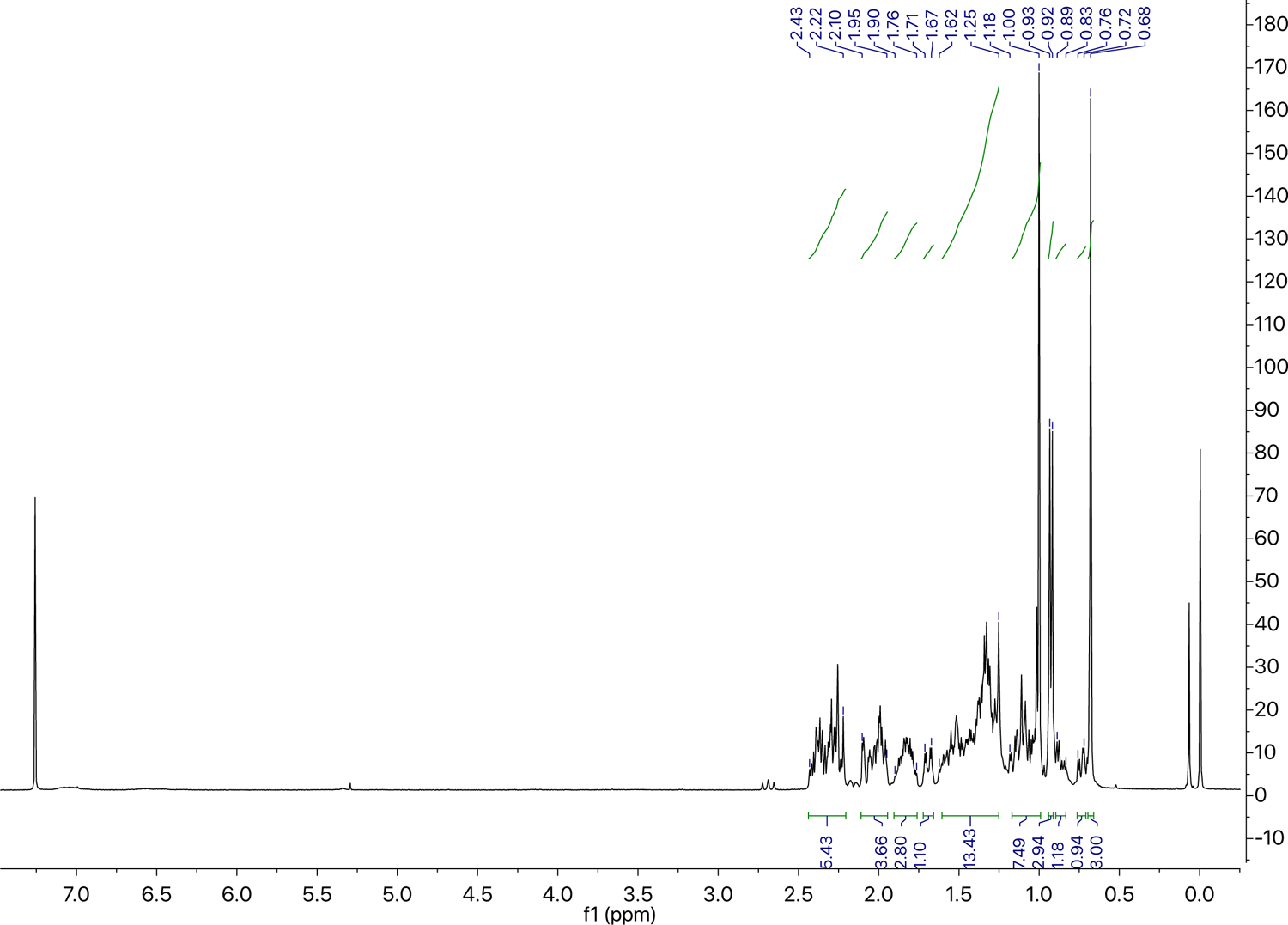
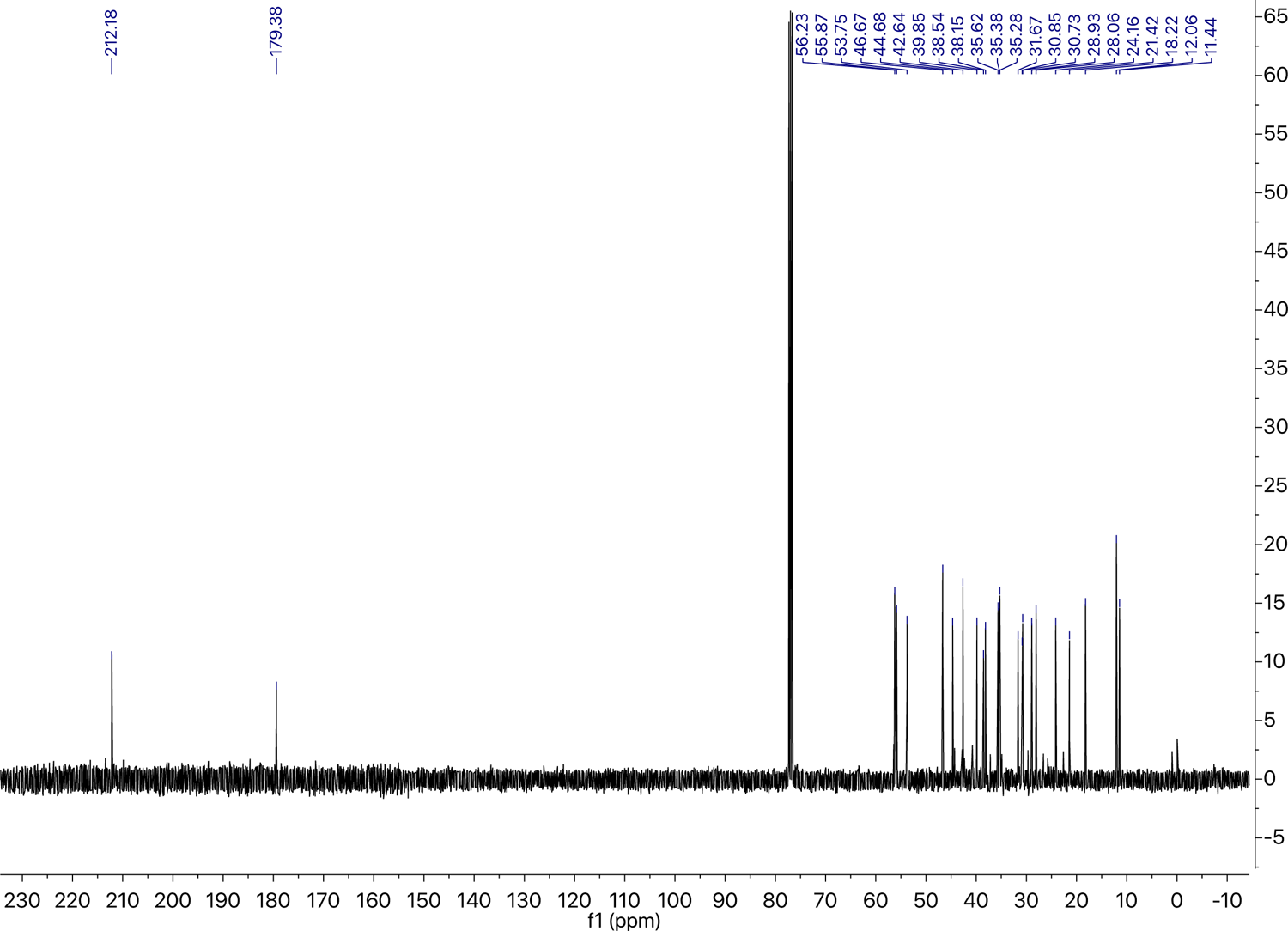

---

**Supplementary Information Table 1.**
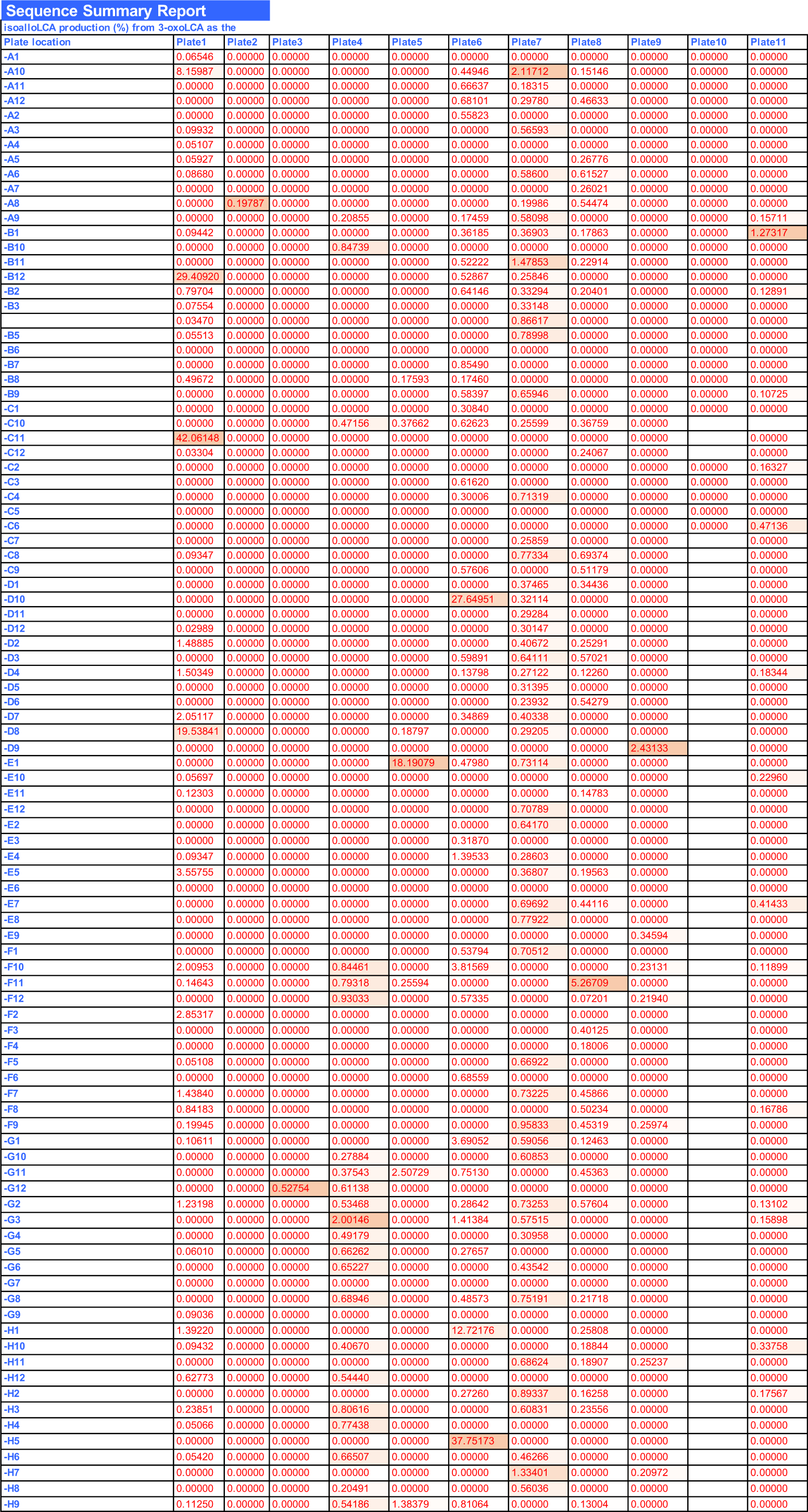
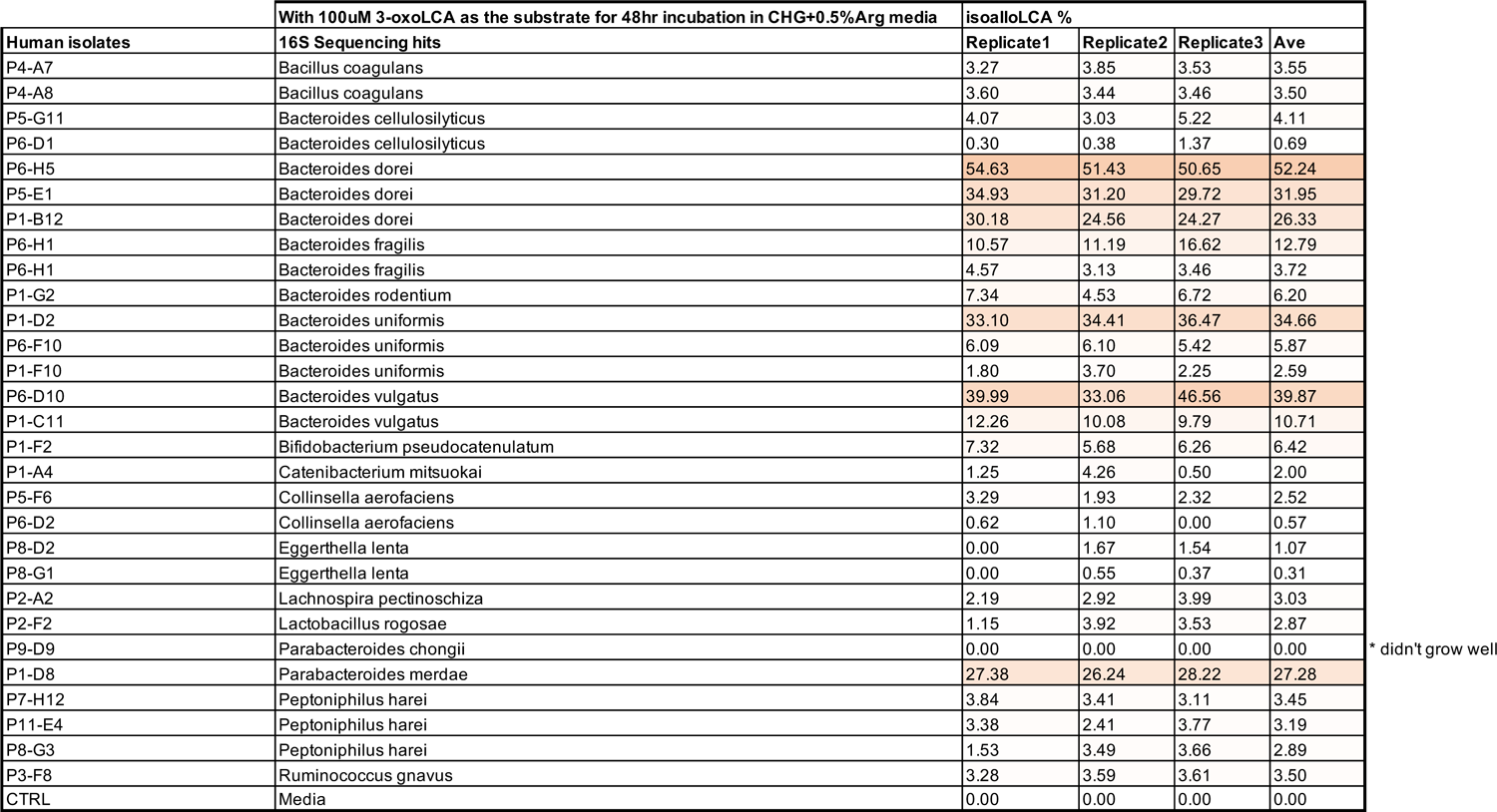
Human screen for isoalloLCA producer strains. Table fields in the first sheet named RAW indicate eleven 96-well plates containing 990 colonies from two patients’ stool samples (see Methods). Assays were performed in plates using 100 μM 3-oxoLCA as a substrate (no isoalloLCA was detected using LCA as the substrate). IsoalloLCA production in percentage conversion for each well is shown. Table fields on the next page include positive bacterial metabolizers that were verified in bacterial culture tubes in biological triplicate and their conversion rates.

**Supplementary Information Table 2.**
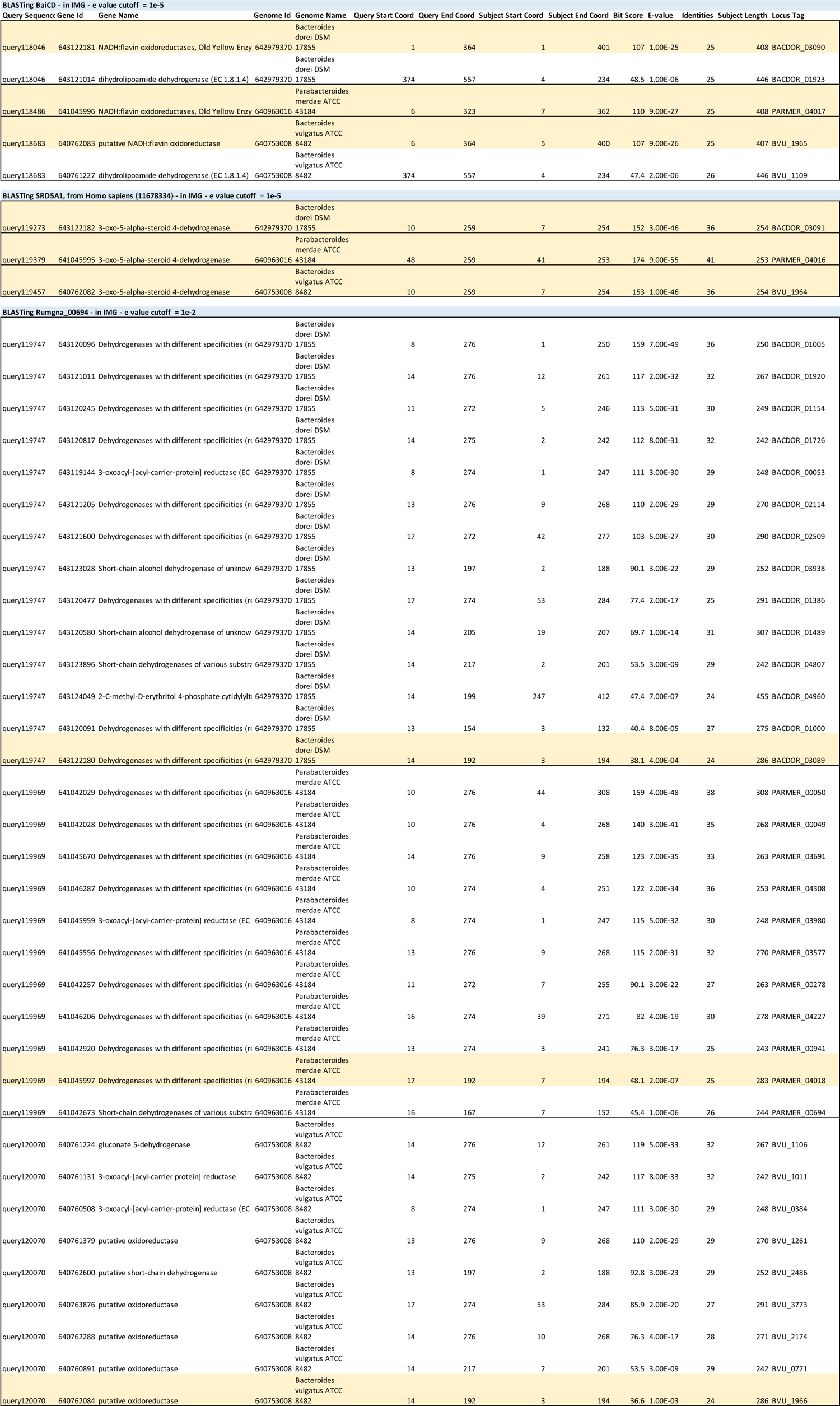
BLASTP searches revealed a three-gene cluster in the isoalloLCA producers *Bacteroides dorei* DSM 17855, *P. merdae* ATCC 43184, and *B. vulgatus* ATCC 8489. BLASTP alignment of 3 Bacteroidetes genomes against known 5β-reductase (baiCD), 5α-reductase (SRD5A1) and 3β-HSDH (Rumgna_00694) sequences.

**Supplementary Information Table 3.**
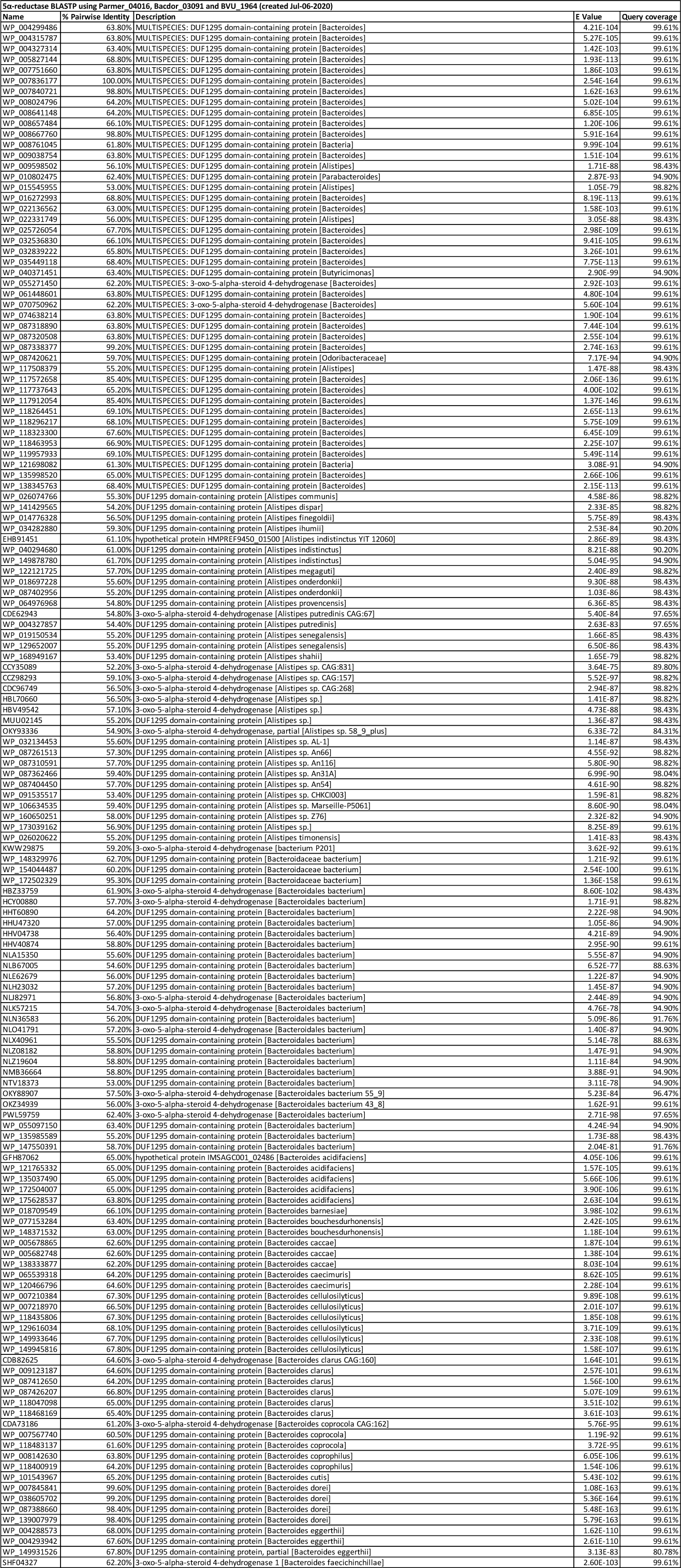
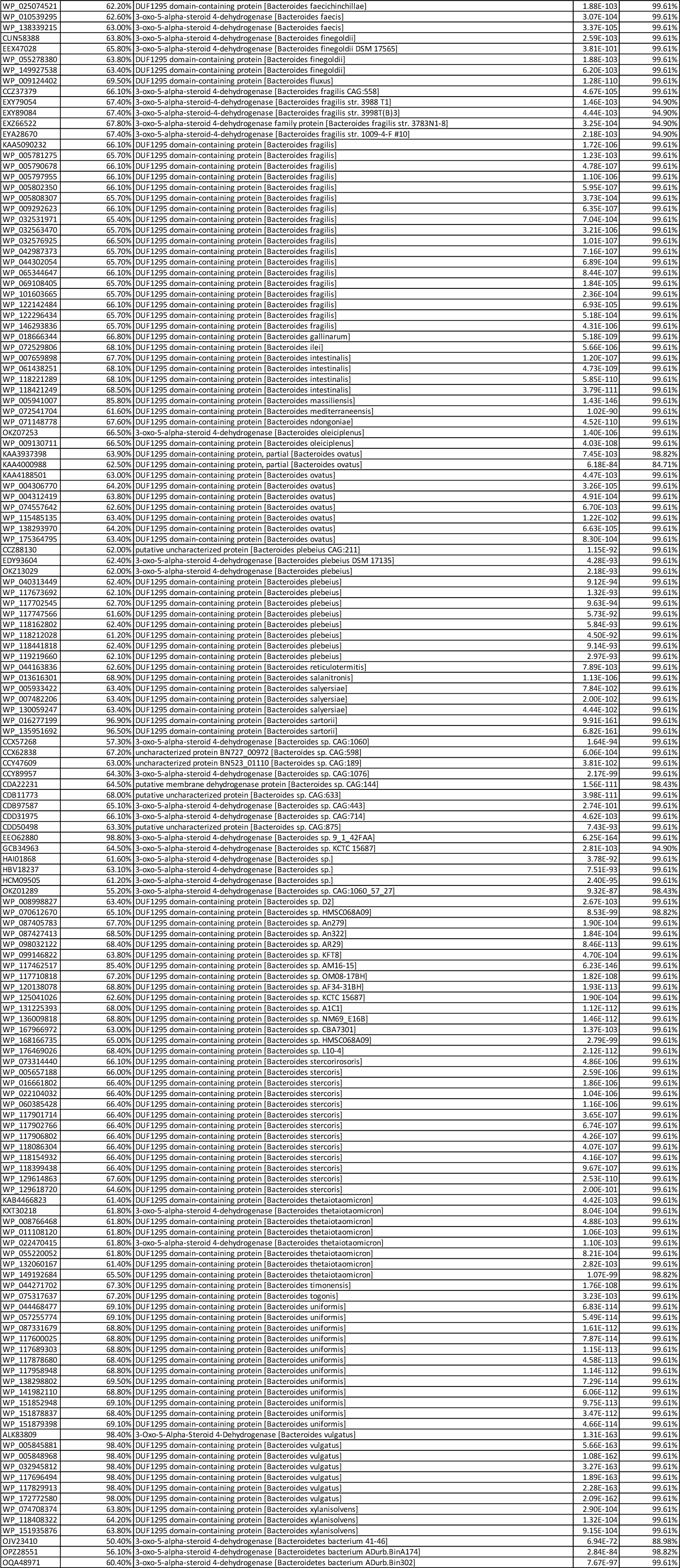
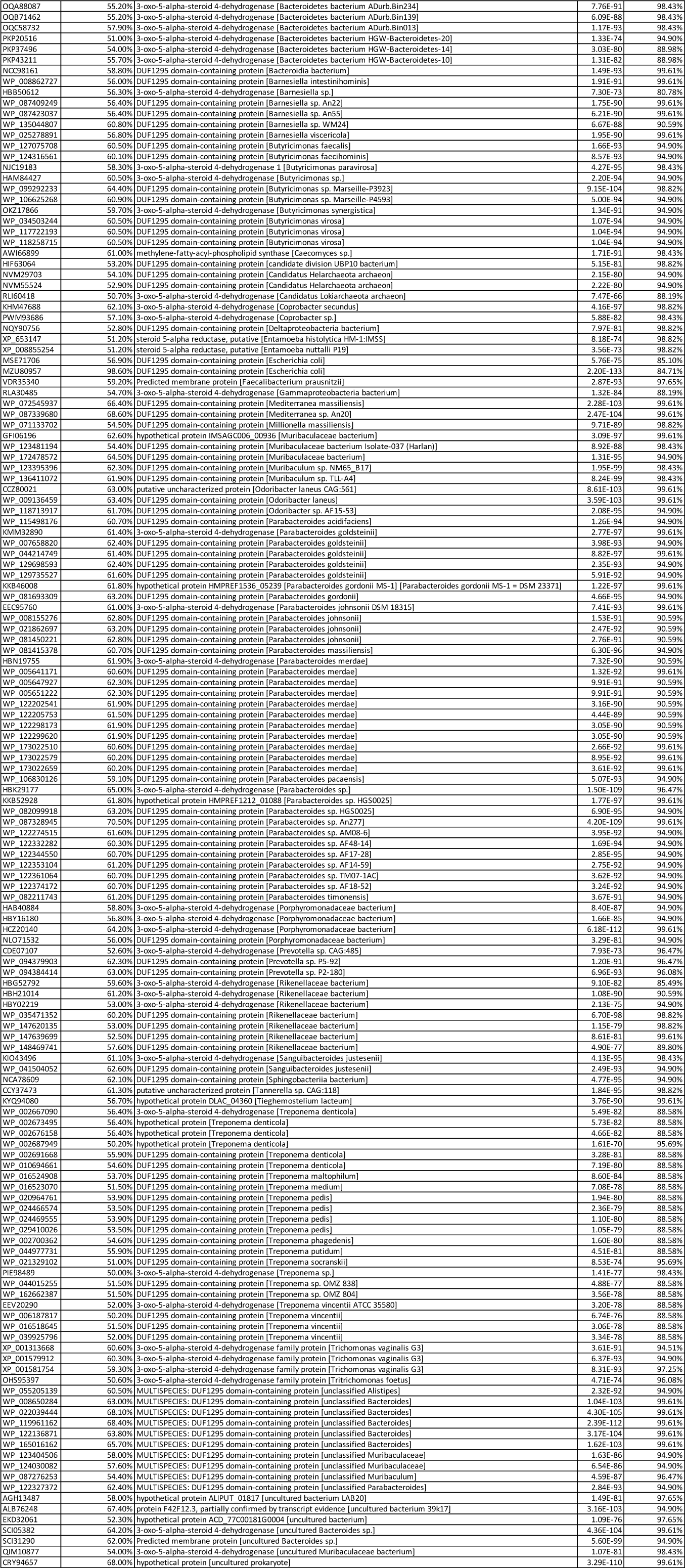
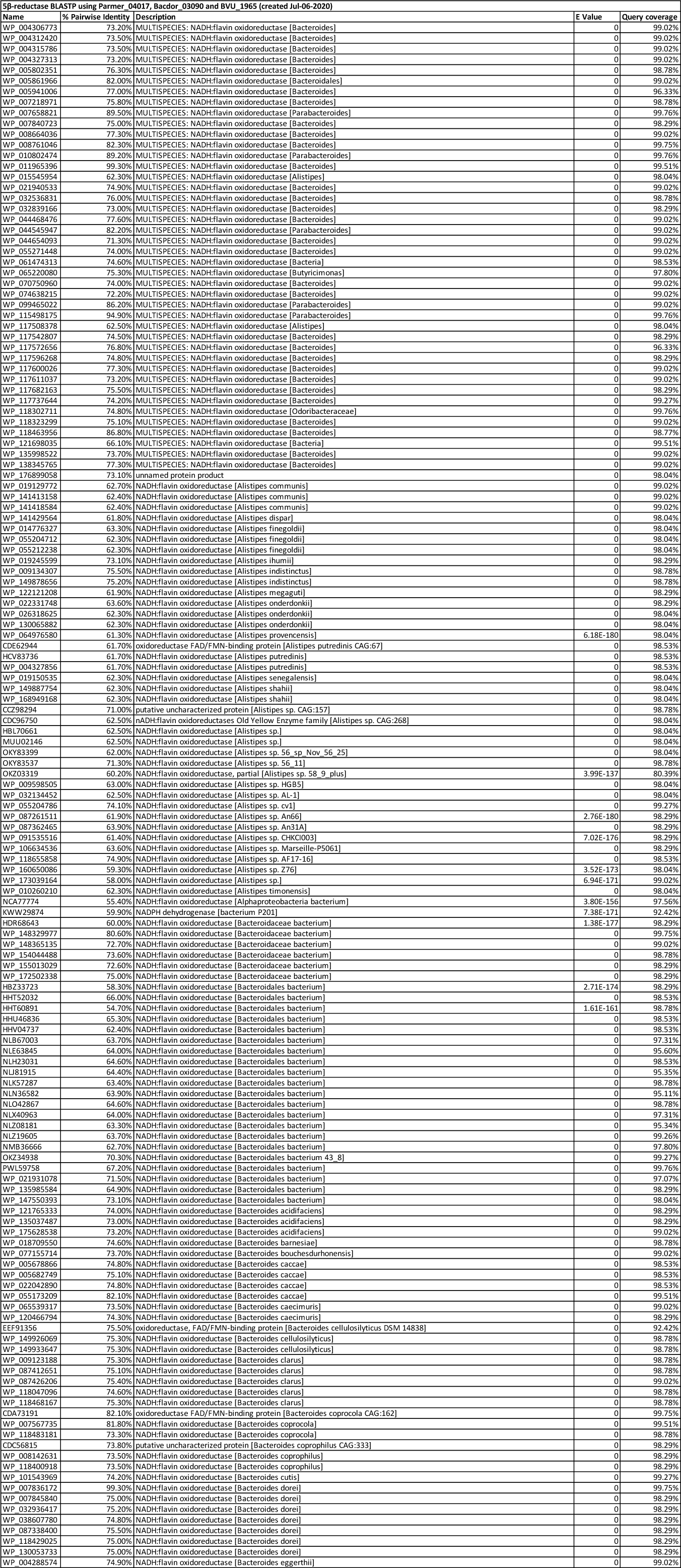
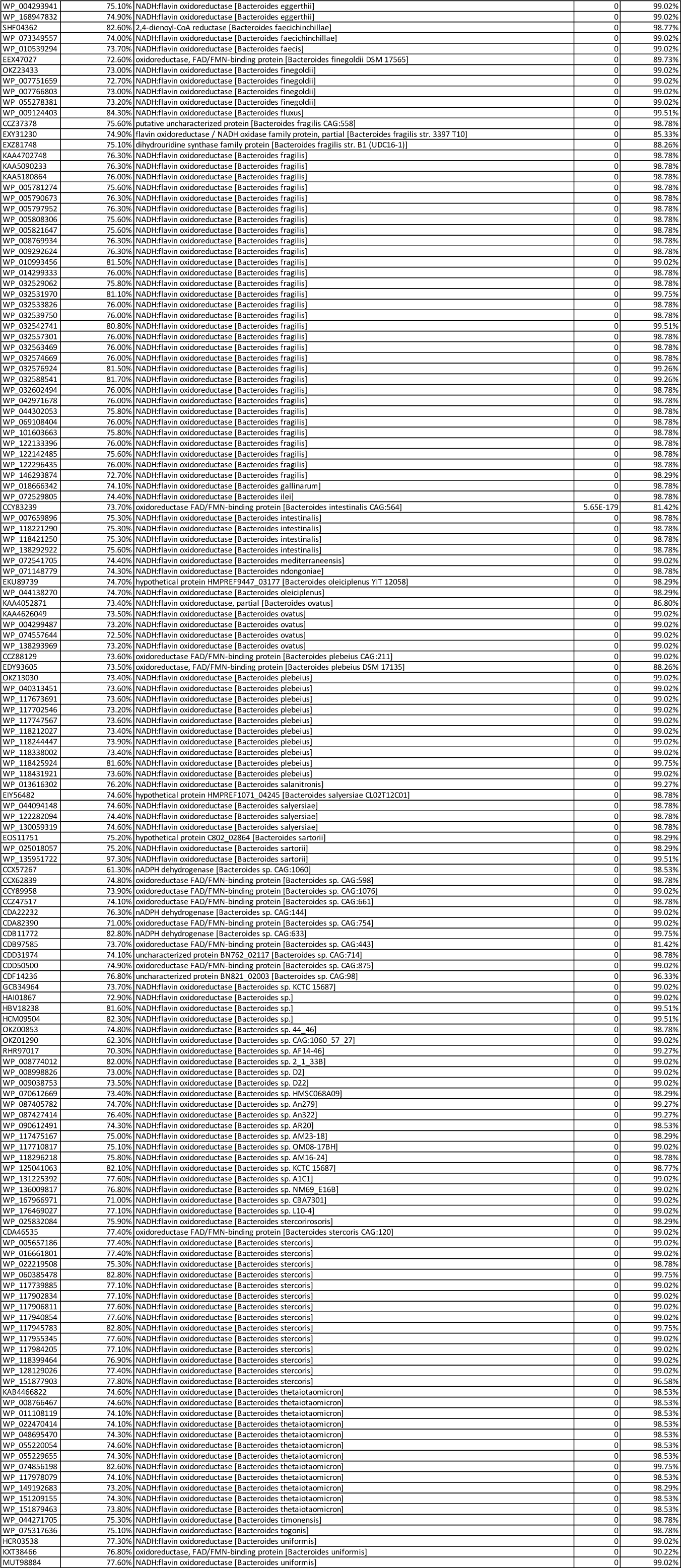
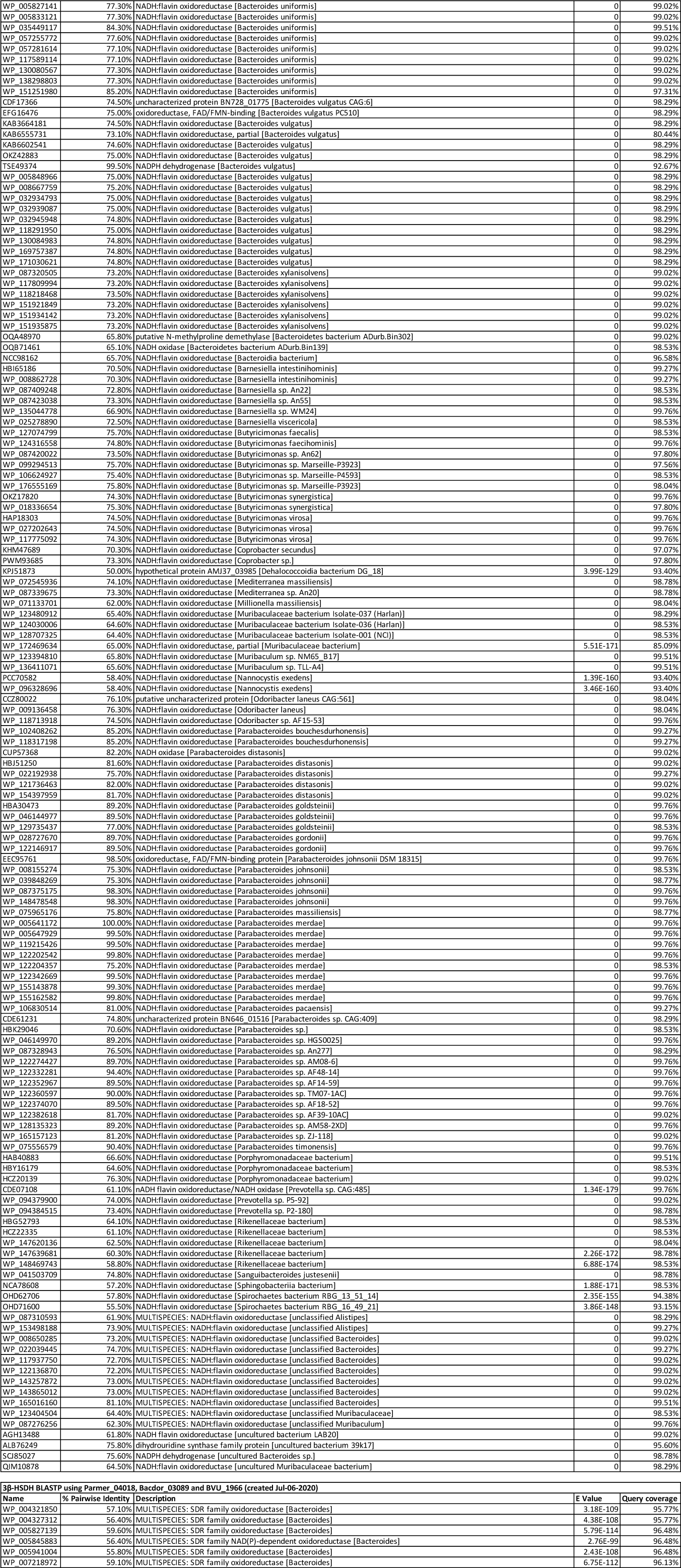
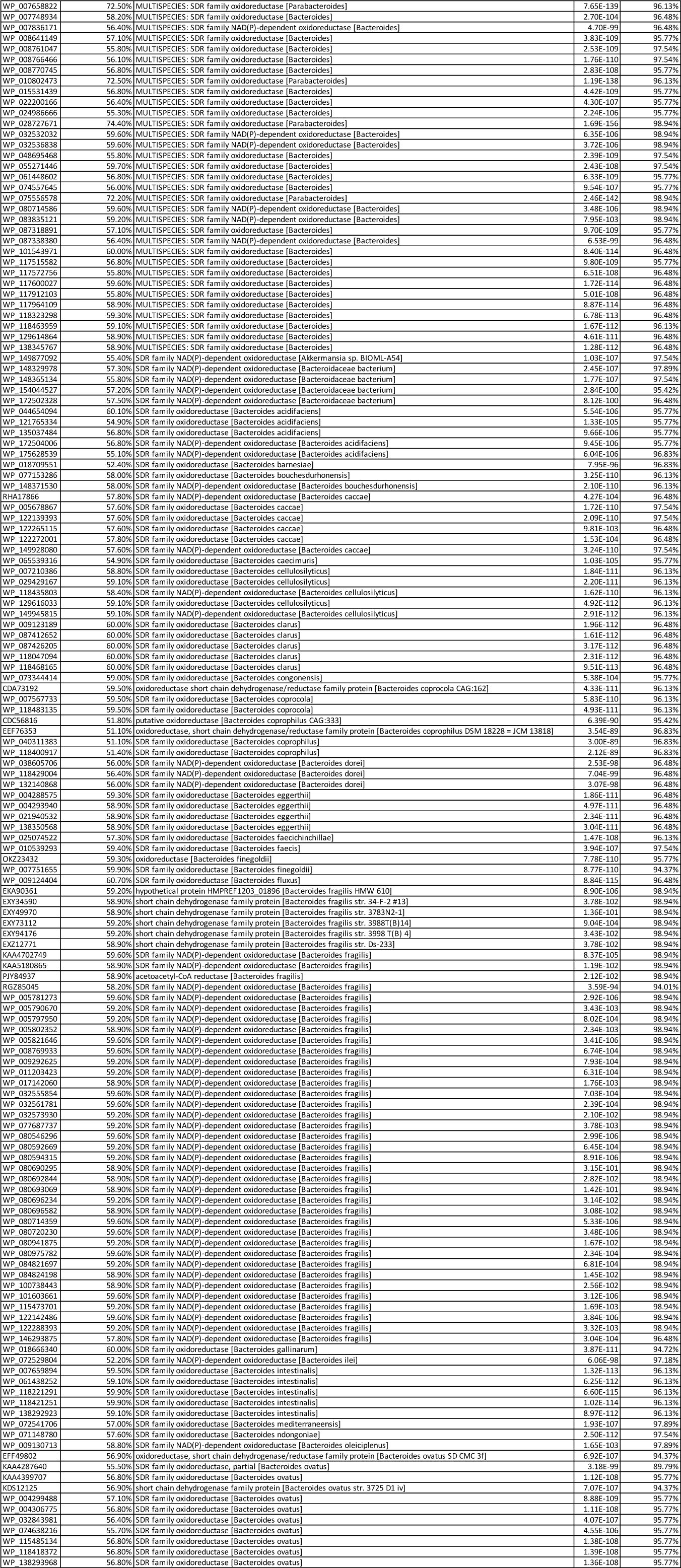
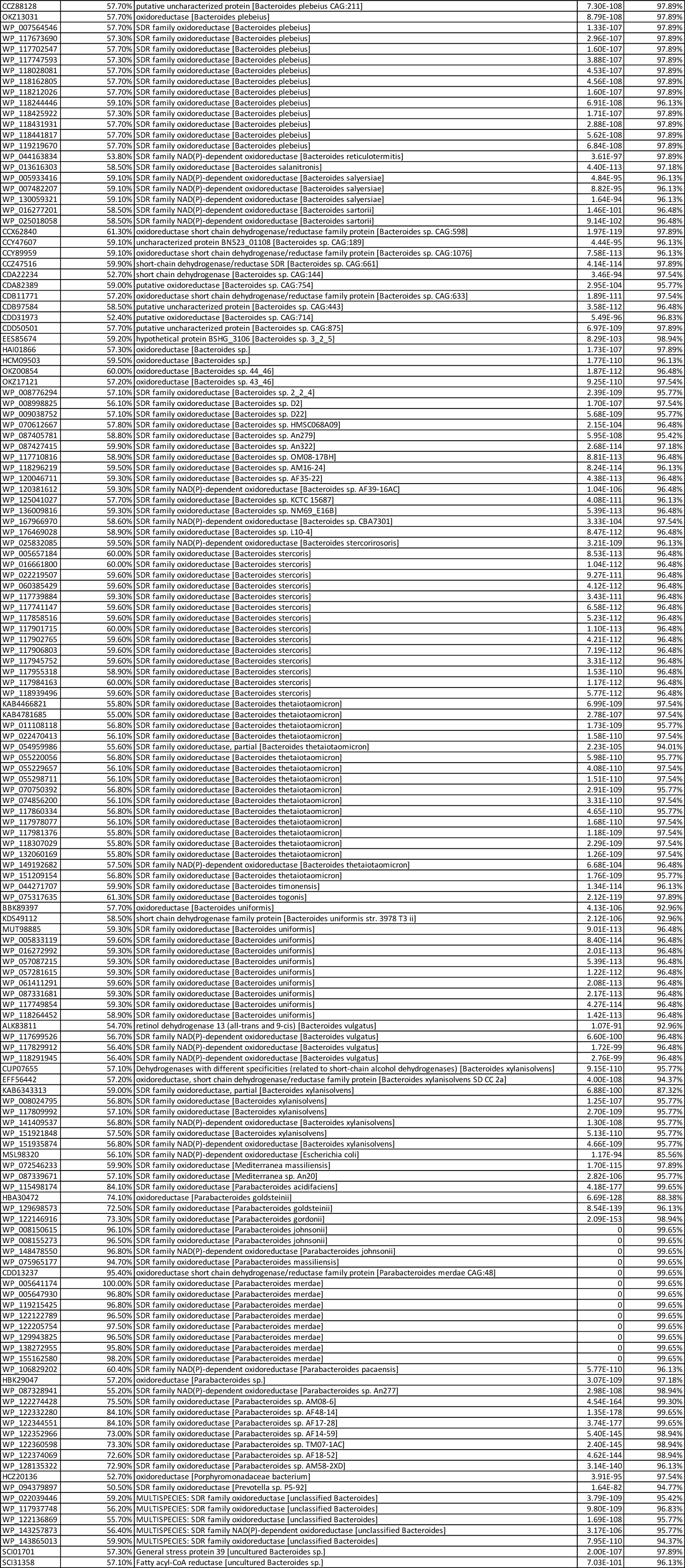
BLASTP searches suggested a variety of bacteria contain the cluster genes. BLASTP alignment of NCBI nucleotide collection database (nr) against confirmed genes in *Bacteroides dorei* DSM 17855, *P. merdae* ATCC 43184, and *B. vulgatus* ATCC 8489 (Figure 2C).

**Supplementary Information Table 4.** Full list of transcription factor binding motifs exclusively enriched in ATAC-seq peaks that are more accessible in isoalloLCA treated cells. The first column in the table contains the names of the TF families whose motifs are exclusively enriched in ATAC-seq peaks that are more accessible in isoalloLCA treated cells. The second column contains the names of candidate TFs within each corresponding TF family. The third column shows the corresponding p values of the TFs in the second column (same order as the TFs). The candidate TFs shown were limited to the TFs that were within 2 log10(p values) of the smallest p value among all candidate TFs of a specific TF family. The enriched motifs were identified using HOMER with a threshold of p value < 0.01. Note that the binding motifs of NR4A2 and NR4A3 were not in the HOMER database used in the analysis. They were added to the candidate list based on the fact that NR4A2 and NR4A3 share the same core binding motif as NR4A1.

| Full List of ATAC-Seq motifs exclusively enriched in isoalloLCA treated cells |  |  |
| --- | --- | --- |
| TF_family | TFs | p_value |
| NR | NR4A1,NR4A2,NR4A3 | 1e-23 |
| ETS | Elk1,EHF,ELF3 | 1e-22,1e-20,1e-20 |
| Stat | STAT5 | 1e-22 |
| Homeobox | Tgif2,Tgif1 | 1e-14,1e-12 |
| HMG | Tcf3 | 1e-09 |
| Zf | BORIS | 1e-09 |
| MAD | Smad2,Smad4 | 1e-08,1e-06 |
| bZIP | c-Jun-CRE,Atf7,Atf1,Atf4,Atf2 | 1e-07,1e-06,1e-06,1e-05,1e-05 |
| bHLH | Myf5,Tcf12,Ap4,MITF,Tcf21 | 1e-05,1e-04,1e-04,1e-04,0.001 |
| T-box | Tbx5,Tbet | 1e-04,0.01 |
| POU,Homeobox | Brn1,Oct6 | 1e-04,0.01 |
| HTH | AMYB,BMYB,Rfx6 | 0.001,0.001,0.001 |
| IRF | T1ISRE | 0.001 |
| MADS | Mef2d,Mef2c | 0.001,0.01 |
| Forkhead | Foxo1 | 0.001 |
| CTF | NF1-halfsite | 0.01 |
| E2F | E2F,E2F3 | 0.01,0.01 |

**Supplementary Information Table 5.**
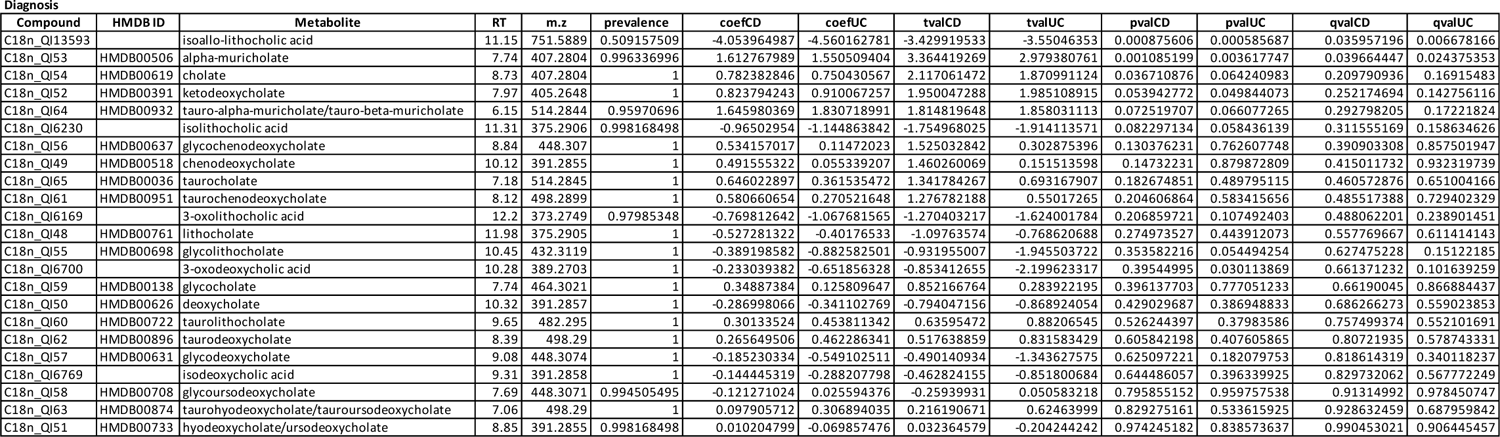
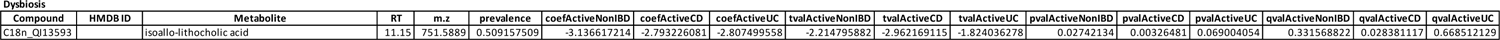
Differentiation abundances of identified bile acid metabolites from the HMP2 metabolomic database. List of bile acids metabolites and their differential abundances in different disease phenotypes (CD/UC/non-IBD) within HMP2 fecal metabolomes, using multivariate association testing with a linear mixed effects model (see Methods). For each bile acid metabolite, the prevalence of metabolites across the HMP2 cohort, coefficient estimates, test statistics, associated two-tailed p-values and FDR-adjusted p-value between CD vs. non-IBD and UC vs. non-IBD are reported. Differential abundance of isoalloLCA with dysbiosis status was also tested in the same linear mixed effects model. Prevalence of isoalloLCA across the HMP2 cohort, coefficient estimates, and test statistics (CD dysbiosis vs. CD non-dysbiosis, UC dysbiosis vs. UC non-dysbiosis) and the associated two-tailed p-value and FDR-adjusted p-value are also reported.

**Supplementary Information Table 6.** Differential abundance modeling of 5α-reductase, 5β-reductase and 3β-HSDH homologs in HMP2 and PRISM. Table fields indicate homolog ID; the prevalence of each homolog in each disease phenotype; effect size estimated from the linear model’s dysbiosis coefficient; nominal p-values from the linear mixed-effects models (see Methods); and adjusted two-tailed p-values based on the Benjamini-Hochberg procedure with target false discovery rate (FDR) of 0.05.

**PRISM - metagenomics**
| gene | prevalence.CD | prevalence.UC | prevalence.nonIBD | coef.CD | coef.UC | pval.CD | pval.UC | qval.CD | qval.UC |
| --- | --- | --- | --- | --- | --- | --- | --- | --- | --- |
| 5α-reductase | 0.264705882 | 0.41509434 | 0.735294118 | -0.542266911 | -0.383526884 | 0.00062766 | 0.022492652 | 0.006904263 | 0.247419176 |
| 5β-reductase | 0.926470588 | 1 | 1 | -0.308871794 | -0.250959398 | 0.114236071 | 0.230448586 | 0.184603019 | 0.365068971 |
| 3β-HSDH | 0.779411765 | 0.867924528 | 1 | -0.408722525 | -0.348681578 | 0.133902006 | 0.232316618 | 0.184603019 | 0.365068971 |

**HMP2 - metagenomics**
| gene | pre.CD-dys | pre.CD-nondys | pre.UC-dys | pre.UC-nondys | prev.nonIBD-dys | prev.nonIBD-nondys | coef.CD-dys | coef.UC-dys | pval.CD-dys | pval.UC-dys | qval.CD-dys | qval.UC-dys |
| --- | --- | --- | --- | --- | --- | --- | --- | --- | --- | --- | --- | --- |
| 5α-reductase | 0.188405797 | 0.530337079 | 0.261904762 | 0.536977492 | 0.184210526 | 0.638888889 | -0.264286244 | -0.124968947 | 0.00061518 | 0.217896072 | 0.000676698 | 0.217896072 |
| 5β-reductase | 0.934782609 | 0.997752809 | 0.976190476 | 0.996784566 | 0.921052632 | 0.990740741 | -0.284491047 | -0.213675247 | 7.98E-09 | 0.000760302 | 1.10E-08 | 0.000929258 |
| 3β-HSDH | 0.695652174 | 0.979775281 | 0.857142857 | 0.993569132 | 0.578947368 | 0.978395062 | -0.461342324 | -0.371059998 | 1.71E-17 | 7.15E-08 | 4.69E-17 | 1.57E-07 |

